# Microbiome-based disease prediction with multimodal variational information bottlenecks

**DOI:** 10.1101/2021.06.08.447505

**Authors:** Filippo Grazioli, Raman Siarheyeu, Israa Alqassem, Andreas Henschel, Giampaolo Pileggi, Andrea Meiser

## Abstract

Scientific research is shedding light on the interaction of the gut microbiome with the human host and on its role in human health state. Existing machine learning methods have shown great potential in discriminating healthy from diseased microbiome states. Most of them leverage shotgun metagenomic sequencing to extract gut microbial species-relative abundances or strain-level markers. Each of these gut microbial features showed diagnostic potential when tested separately; however, no existing approach combines them in a single predictive framework. Here, we propose the Multimodal Variational Information Bottleneck (MVIB), a novel deep learning model capable of learning a joint representation of multiple heterogeneous data modalities. MVIB achieves competitive classification performance while being faster than existing methods. Additionally, MVIB offers interpretable results. Our model adopts an information theoretic interpretation of deep neural networks and computes a joint stochastic encoding of different input data modalities. We use MVIB to predict whether human hosts are affected by a certain disease by jointly analysing gut microbial species-relative abundances and strain-level markers. MVIB is evaluated on human gut metagenomic samples from 11 publicly available disease cohorts covering 6 different diseases. We achieve high performance (0.80 < ROC AUC < 0.95) on 5 cohorts and at least medium performance on the remaining ones. We adopt a saliency technique to interpret the output of MVIB and identify the most relevant microbial species and strain-level markers to the model’s predictions. We also perform cross-study generalisation experiments, where we train and test MVIB on different cohorts of the same disease, and overall we achieve comparable results to the baseline approach. Further, we evaluate our model by adding metabolomic data derived from mass spectrometry as a third input modality. Our method is scalable with respect to input data modalities and has an average training time of < 1.4 seconds. The source code and the datasets used in this work are publicly available.

**Author summary:** The gut microbiome can be an indicator of various diseases due to its interaction with the human system. Our main objective is to improve on the current state of the art in microbiome classification for diagnostic purposes. A rich body of literature evidences the clinical value of microbiome predictive models. Here, we propose the Multimodal Variational Information Bottleneck (MVIB), a novel deep learning model for microbiome-based disease prediction. MVIB learns a joint stochastic encoding of different input data modalities to predict the output class. We use MVIB to predict whether human hosts are affected by a certain disease by jointly analysing gut microbial species-relative abundance and strain-level marker profiles. Both of these gut microbial features showed diagnostic potential when tested separately in previous studies; however, no research has combined them in a single predictive tool. We evaluate MVIB on various human gut metagenomic samples from 11 publicly available disease cohorts. MVIB achieves competitive performance compared to state-of-the-art methods. Additionally, we evaluate our model by adding metabolomic data as a third input modality and we show that MVIB is scalable with respect to input feature modalities. Further, we adopt a saliency technique to interpret the output of MVIB and identify the most relevant microbial species and strain-level markers to our model predictions.

## Introduction

The human microbiota consist of various microbial communities that live in and on our bodies. These communities are composed of different species of bacteria, archaea, protists, fungi and viruses [1]. Together, they constitute complex and diverse ecosystems that interact with the human host. When we refer to microbiota together with their genomic information, we use the term *microbiome.* Previous research estimated that the genes in the gut microbiome alone outnumber the human genes by two orders of magnitude [2]. Recent studies showed that the human microbiota play key roles in human health state [3]. The presence of microbiota benefits the host as they enable important chemical processes, e.g., they maintain homeostasis, develop the immune system and help in harvesting various nutrients that are otherwise inaccessible [3,4]. Previous research reported that altered states of microbiota can contribute to carcinogenesis and affect therapeutic response in cancer patients [5]. [6] reviewed the role of microbiota in human health and disease and anticipated a potential use of microbiota analysis for disease diagnosis and prediction.

Over the past two decades, several large-scale microbial profiling projects were established, such as the Human Microbiome Project [7] and the MetaHIT (Metagenomics of the Human Intestinal Tract) project [8]. These projects aimed at investigating the nature of the microbial components of the human genetic and metabolic landscape and their link to various diseases. However, despite various attempts to develop unified best practices, truly standardised approaches in microbiome research have not yet been established [9–11]. Therefore, we are in need for various statistical and machine learning models that leverage high throughput metagenomic data with supervised and unsupervised learning techniques.

Shotgun metagenomic sequencing allows comprehensive sampling of all genes in all microorganisms that are present in a given sample. This technology enables researchers to examine microbial diversity and to detect their abundances in different environments. In comparison to 16S rRNA gene sequencing technology, shotgun metagenomic sequencing provides higher resolution profiles at species and strain levels. Existing machine learning methods leverage shotgun metagenomics to extract gut microbial species abundance or strain-level marker profiles to differentiate healthy from diseased human hosts. Both of these gut microbial features showed diagnostic potential [12, 13] and have been used separately in previous research for microbiome-based disease prediction.

Current microbiome-based disease prediction approaches either use species-relative abundance or strain-level marker profiles. [14,15] solve the disease prediction task by applying deep learning to abundance profiles from human gut microbiome. MetAML [12] solves the disease prediction task by applying classical machine learning algorithms to either abundance or marker profiles. MicroPheno [16] sub-samples 16S rRNA sequences via bootstrapping, then computes k-mer representations of the sub-sampled sequences, after that it uses the produced k-mer representations for disease prediction. DeepMicro [17] leverages deep representation learning with autoencoders to compute encodings of either microbiome species abundance or marker profiles, i.e., it transforms high-dimensional microbiome data into a low-dimensional representation, then it applies classical machine learning classification models on the generated representations for disease prediction. [18] solves the prediction of cardiovascular disease by using supervised learning on taxonomic features, i.e., microbial taxa. PopPhy-CNN [19] represents microbial phylogenetic tree and relative abundances of microbial taxa in a single matrix and solves disease prediction via a convolutional neural network (CNN). The SIAMCAT R package [20] provides a toolbox for statistical inference of associations between microbial communities and host phenotypes. Its feature matrix consists of abundances of microbial taxa, genes, or pathways across all samples, in addition to optional meta-variables, such as demographics, lifestyles, donor clinical records.

Also related to our work is [21], which explores various statistical methodologies for the analysis of multi-table heterogeneous data for microbiome research. This work combines the analysis of body mass composition information and 16S rRNA abundances in a single computational framework, but does not specifically address disease prediction.

A rich body of literature evidences the clinical value of microbiome predictive models, e.g., [22]. Hence, our main objective is to improve the current state of the art in microbiome-based disease classification for diagnostic purposes by combining multimodal data sources. To the best of our knowledge, none of the existing approaches is capable of solving this task by efficiently combining features from heterogeneous data modalities. Here, we present the Multimodal Variational Information Bottleneck (MVIB), a novel multimodal generalisation of the Deep Variational Information Bottleneck (Deep VIB) [23]. MVIB is a microbiome-disease classification method. It leverages the theory of the Information Bottleneck (IB) [24] to learn a meaningful joint encoding from different input data modalities, e.g. species-relative abundance, strain-level marker profiles and metabolomic data. The learned joint encoding by MVIB is maximally compressive of the heterogeneous input data modalities and at the same time is maximally expressive of the target class, i.e. diseased or healthy human host. By design, MVIB is scalable with respect to input data modalities.

We evaluate MVIB on 11 different metagenomic datasets from human gut microbiome. In this paper, we show how MVIB performs when combining species relative abundances and strain-level markers. Additionally, we demonstrate how MVIB works in a trimodal setting by adding metabolomic data as a third modality. We benchmark MVIB against state-of-the-art methods, i.e. DeepMicro, PopPhy-CNN and Random Forest. Additionally, we adopt a saliency technique derived from computer vision literature [25] to interpret the output of MVIB and identify most discriminative microbial species and strain-level markers with respect to various human diseases. Furthermore, we perform various transfer learning [26,27] experiments, as well as cross-study generalisation experiments where we train and test MVIB on different cohorts of the same disease.

## Materials and methods

### Datasets

For evaluation and comparative benchmark analysis, we consider publicly available human gut metagenomic samples from 11 different cohorts that cover 6 different diseases. These diseases are inflammatory bowel disease (*IBD*), type 2 diabetes in Europe (women) (*EW-T2D*) and in China (women and men) (*C-T2D*), obesity (*Obesity, Obesity-Joint*), liver cirrhosis (*Cirrhosis*), colorectal cancer (*Colorectal, Colorectal-EMBL, Early-Colorectal-EMBL, Colorectal-Metabolic*) and chronic high blood pressure (*Hypertension*). The names in parentheses indicate the cohort identifiers that we used in this work. The number of affected and control subjects in each cohort are listed in Table 1.

**Table 1.** Datasets overview.

| Dataset name | Samples | Controls | Affected | Data source | Modalities |
| --- | --- | --- | --- | --- | --- |
| IBD | 110 | 85 | 25 | [12], [8] | Abundance, Markers |
| EW-T2D | 96 | 43 | 53 | [12], [29] | Abundance, Markers |
| C-T2D | 344 | 174 | 170 | [12], [30] | Abundance, Markers |
| Obesity | 253 | 89 | 164 | [12], [31] | Abundance, Markers |
| Obesity-Joint | 331 | 117 | 214 | [12], [31], [34], [35] | Abundance, Markers |
| ΔObesity | 78 | 28 | 50 | [34], [35] | Abundance, Markers |
| Cirrhosis | 232 | 114 | 118 | [12], [32] | Abundance, Markers |
| Colorectal | 121 | 73 | 48 | [12], [33] (cohort F) | Abundance, Markers |
| Colorectal-EMBL | 199 | 103 | 96 | [33] (cohorts F/G) | Abundance, Markers |
| ΔColorectal | 78 | 30 | 48 | [33] (cohorts F/G) | Abundance, Markers |
| Early-Colorectal-EMBL | 96 | 52 | 44 | [33] (cohorts F/G) | Abundance, Markers |
| Hypertension | 196 | 40 | 156 | [36] | Abundance, Markers |
| Colorectal-Metabolic | 347 | 127 | 220 | [37] | Abundance, Markers, Metabolites |

We obtained pre-processed human gut metagenomic data for the IBD, EW-T2D, C-T2D, Obesity, Cirrhosis and Colorectal cohorts from the MetAML repository [12]. We also considered five additional disease cohorts, i.e. Obesity-Joint [12, 31, 34, 35], Colorectal-EMBL [33] (cohorts F and G), Early-Colorectal-EMBL [33], Colorectal-Metabolic [37] and Hypertension [36]. For the latter cohorts, we performed the data pre-processing steps highlighted in Fig 1A and B and described later in detail in Section Pre-processing.

**Fig 1.**
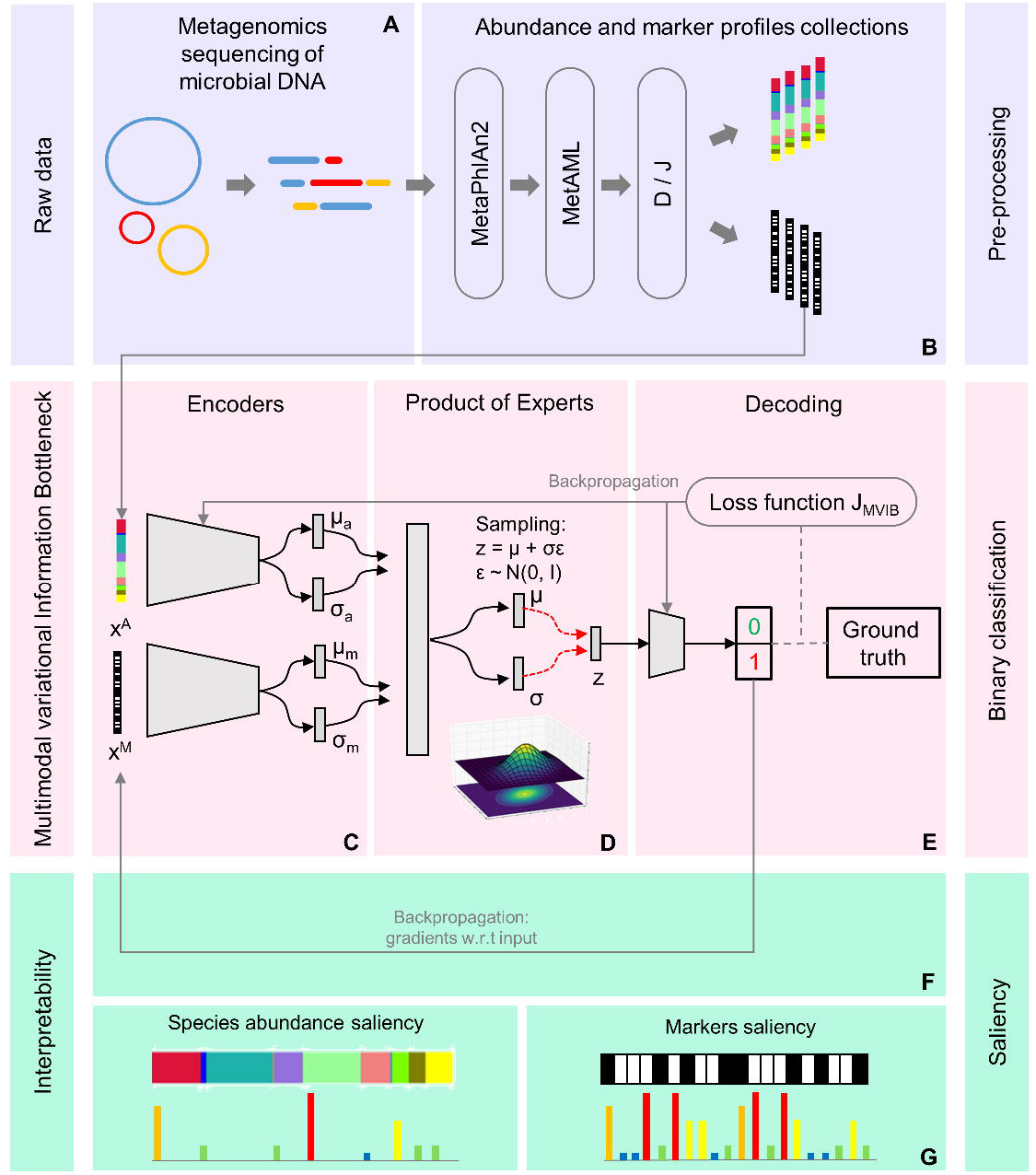
Full workflow. (A) The raw data, i.e., shotgun metagenomic sequencing data of the human gut microbiome. (B) For pre-processing, we leverage MetaPhlAn2 and MetAML to extract species-relative abundances and strain-level markers. We consider two pre-processing schemes to produce two different dataset collections, *default* (D) and *joint* (J). (C) A high-level representation of the probabilistic encoders of the MVIB model. (D) The Product of Experts computes a single joint posterior i.e. ***z***; the joint posterior ***z*** is sampled with the reparametrisation trick [28]. (E) A logistic regression decoder estimates the probability of whether a subject is affected by a certain disease. (F, G) The gradients of the output class are computed with respect to (w.r.t) the input vectors and used to compute saliency maps.

Early-Colorectal-EMBL presents 96 affected samples from the Colorectal-EMBL dataset, where 52 subjects are labelled as “early stage” (colorectal cancer stages 0, I and II) and 44 are labelled as “late stage” (stages III and IV). The Hypertension cohort presents samples from Chinese individuals with pre-hypertension (pHTN) or primary hypertension (HTN), as well as healthy control samples [36]; we consider both pHTN and HTN subjects (156 in total) as affected. The Colorectal dataset from MetAML repository is a subset of Colorectal-EMBL, therefore we create Δ*Colorectal = Colorectal-EMBL – Colorectal* to address the samples which only belong to the larger dataset. Similarly, Obesity is a subset of Obesity-Joint, hence we create Δ*Obesity = Obesity-Joint – Obesity.*

In addition to the datasets discussed so far, which present two data modalities, we performed experiments on a trimodal dataset. To this end, we included metabolite profiles in addition to species-relative abundance and the strain-level marker profiles. We extracted 347 samples with both metabolomic and metagenomic data from [37], which proposes a large cohort of participants who underwent colonoscopy to assess taxonomic and functional characteristics of the gut microbiota and metabolites. The metabolomic data was extracted by means of capillary electrophoresis time-of-flight mass spectrometry (CE-TOFMS). In this dataset, 220 samples belong to subjects affected by colorectal cancer, polypoid adenomas or intramucosal carcinomas, in addition to more advanced lesions. The remaining 127 samples belong to healthy individuals. We refer to this trimodal dataset as *Colorectal-Metabolic*.

All datasets include ground truth labels (i.e., healthy or affected) which refer to the time the microbiome samples were collected. Hence, in this work, we do not predict a future health status.

### Pre-processing

We run all metagenomic samples that were not taken from [12] (i.e. Colorectal-EMBL, Early-Colorectal-EMBL, Obesity-Joint, Hypertension and Colorectal-Metabolic datasets) through a quality control and MetaPhlAn2-based annotation pipeline. This allowed us to get species-relative abundance and strain-level marker profiles in the same format as the datasets taken from the MetAML repository [12].

We downloaded whole-genome shotgun metagenomic sequencing data from the bioproject repositories of the National Center for Biotechnology Information (NCBI). Raw read data for each sequencing run were converted into FASTQ format using fastq-dump version 2.8.0 (NCBI SRA Toolkit [38]) and aggregated by sample identifiers. Afterwards, we used Kneaddata version 0.7.4 [39] with default parameters to perform quality control of the sequencing reads and remove reads of length length < 60 base pairs (bp). Finally, we run MetaPhlAn2 (Metagenomic Phylogenetic Analysis) [40] for profiling the compositions of microbial communities from the quality-controlled data. MetAML `dataset_selection.py` [12] was used to leave only species-level information in the abundance profiles. Obtained species-relative abundances and strain-level markers make up the feature vectors of our machine learning model.

We created two collections of cohort datasets, i.e. *default* and *joint.* Each collection includes all the cohorts. The datasets in the *default* (D) collection are obtained with the MetaPhlAn2+MetAML pre-processing described above. In the *joint* (J) collection, species abundances and strain-level markers are homogeneous across all datasets. We achieved this by taking the union of all features from the various datasets. This guarantees that species abundance and strain-level marker profiles have the same dimensionality across cohorts. We created the *joint* collection for transfer learning (Section Transfer learning) and cross-study generalisation (Section Cross-study generalisation) experiments. We applied feature normalisation to the final sets of species abundances in both collections to obtain species-relative abundances ∈ ℝ^[0,1]^. The dimensionality of the species abundance feature vectors is < 10^3^.

The strain-level marker profiles are represented by a vector of binary variables, where 0 indicates the absence of a certain strain and 1 indicates its presence. The dimensionality of the strain-level markers feature vectors is < 10^5^.

### The Multimodal Variational Information Bottleneck

Let *Y* be a random variable representing a ground truth label associated with a set of multimodal input random variables *X^1^*,…,*X^M^*. In order to provide a more compact notation, let us represent the collection of the available data modalities as a data point *X* = {*X^i^*|*i^th^ modality present*}. Let *Z* be a stochastic encoding of *X* coming from an intermediate layer of a deep neural network and defined by a parametric encoder *p*(***z***|***x***; ***θ***) representing the upstream part of such neural model. For the rest of this manuscript, we adopt the following notation: *X*, *Y*, *Z* are random variables; ***x***, ***y***, ***z*** are multidimensional instances of random variables; *f* (·; ***θ***) are functions parametrised by a vector of parameters ***θ***; 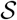 represents a set.

Following the information bottleneck approach [24], our goal consists in learning an encoding *Z* which is (a) maximally informative about *Y* and (b) maximally compressive about *X*. Following an information theoretic approach, objective (a) implies maximising the mutual information *I*(*Z,Y*; ***θ***) between the encoding *Z* and the target *Y*, where:

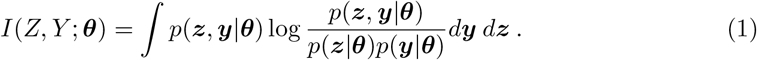

A trivial solution for maximising Eq 1 would be the identity *Z* = *X*. This would ensure a maximally informative representation, but (b) places a constraint on *Z*. In fact, due to (b), we want to “forget” as much information as possible about *X*. This leads to the objective:

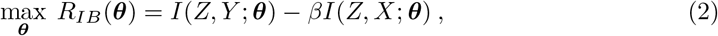

where *β* ≥ 0 is a Lagrange multiplier. The first term on the right hand side of Eq 2 causes *Z* to be predictive of *Y*, while the second term constraints *Z* to be a minimal sufficient statistics of *X*. *β* controls the trade-off.

As derived in [23] for the Deep Variational Information Bottleneck (Deep VIB), assuming *q*(***y***|***z***) and *r*(***z***) are variational approximations of the true *p*(***y***|***z***) and *p*(***z***), respectively, Eq 2 can be rewritten as:

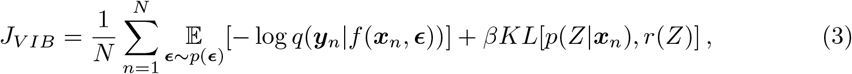

where 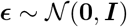 is an auxiliary Gaussian noise variable, *KL* is the Kullback-Leibler divergence and *f* is a vector-valued parametric deterministic encoding function (in this work, a neural network). The introduction of ***ϵ*** consists in the reparameterisation trick [28], which allows to write *p*(***z***|***x***; ***θ***)*d****x*** = *p*(***ϵ***)*d****ϵ***, where ***z*** = *f*(***x***, ***ϵ***) is now treated as a deterministic variable. This formulation allows the noise variable to be independent of the model parameters. This way, it is easy to compute gradients of the objective in Eq 3 and optimise via backpropagation. In this work, we let the variational approximate posteriors be multivariate Gaussians with a diagonal covariance structure 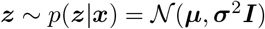; a valid reparameterisation is ***z*** = ***μ*** + ***σϵ***.

We generalise the formulation of the Deep VIB objective of Eq 3 by considering that *X* is a collection of multimodal random input variables s.t.

*X* = {*X^i^*|*i^th^ modality present*}. In light of this, the posterior *p*(*Z*|***x***) of Eq 3 consists actually in the joint posterior *p*(*Z*|***x***^1^,…, ***x***^*M*^), conditioned by the joint *M* available data modalities. Following the approach proposed for the Multimodal Variational Autoencoder [41], assuming conditional independence between the various modalities conditioned on *Z* and approximating *p*(*Z*|***x^i^***) with 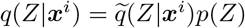, where 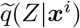 is the stochastic encoder of the *i^th^* data modality and *p*(*Z*) is a prior, the joint posterior can be expressed as a product of single-modality posteriors:

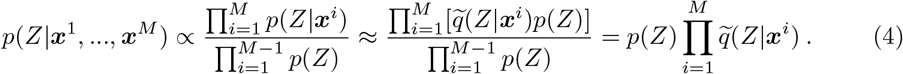

The formulation derived from Eq 4 is addressed as a product of experts (PoE) (see Fig 1D). When the involved probability distributions are Gaussian, the PoE acquires a simple analytical solution, as the product of Gaussian experts is itself a Gaussian [42]. We can now formulate the objective of the Multimodal Variational Information Bottleneck:

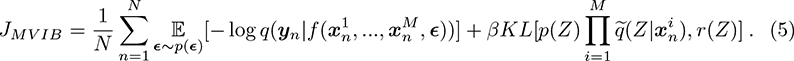

### Implementation details

In this work, there are two main data modalities: the species-relative abundance profiles and the strain-level marker profiles. For each modality, the dedicated stochastic encoder has the form 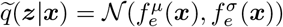 (see Fig 1C). *f_e_* is a multi-layer perceptron (MLP). For abundance profiles, the MLP has 2 layers of the form *input dimension* - *input dimension*/2 - *input dimension*/2, followed by two parallel layers which output 2 vectors of size *K* for ***μ*** and ***σ. K*** is the size of the bottleneck, i.e. the dimension of *Z*. For a more stable computation, we let 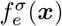 model the logarithm of the variance log ***σ***^2^. For marker profiles, the MLP has 2 layers of the form *input dimension*/2 - 1024 - 1024, followed by two parallel layers which output 2 vectors of size *K* for ***μ*** and ***σ***. SiLU [43] activation functions are used. 0.4 drop-out is used at training time.

As described later, experiments on the Colorectal-Metabolic dataset demand a third additional stochastic encoder for the metabolite profiles. For this data modality, the same encoder architecture adopted for the abundance profiles is used.

The decoder consists in a logistic regression model *q*(***y***|***z***) = *σ*(*f_d_*(***z***)), where *σ*(·) is the logistic sigmoid function and *f_d_*(***z***) = ***w^T^z*** + ***b*** (see Fig 1E). This implements the binary classification. ***y*** models the diagnosis label for a given disease: sick or healthy.

In Eq 5, *r*(*Z*) and *p*(*Z*) are treated as K-dimensional spherical Gaussian distributions, 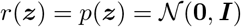. The latent dimension *K* of the encoding is set to 256. *β* is set to 10^-5^.

The networks are trained using the Adam optimiser, with a learning rate of 10^-4^ and a *L*_2_ weight decay with *λ* = 10^-5^. The batch size is set to 256. The training is performed for 200 epochs and, in order to avoid overfitting, the best model is selected by saving the weights corresponding to the epoch where the area under the curve (AUC) of the receiver operating characteristic (ROC) is maximum on the validation set (see Section Validation framework and performance evaluation).

All experiments were performed on a CentOS Linux 8 machine with NVIDIA GeForce RTX 2080 Ti GPUs and CUDA 10.2 installed, with the exception of the transfer learning experiments (see Section Transfer learning), which were performed with an NVIDIA TITAN RTX GPU. Algorithms are implemented in Python 3.6 using PyTorch [44] version 1.7; code is publicly available at https://github.com/nec-research/microbiome-mvib.

### Extending the MVIB objective with the triplet margin loss

The MVIB objective function proposed in Eq 5 consists of two terms: a supervised negative log-likelihood loss and a KL divergence which acts as a regulariser. As presented in Section Results, it was empirically observed that extending the *J_MVIB_* of Eq 5 with an additional triplet margin loss [45] term can lead to a more accurate predictor on various disease datasets.

The triplet margin loss was first introduced in the field of computer vision [45]. The underlying idea consists in explicitly enforcing the latent representation of an input sample (e.g. an image, or a vector) to be close to the latent representations of the samples that belong to the same class, and distant from the latent representations of samples that belong to a different class. The concept of closeness depends on the nature of the latent space. In a Euclidean space, the Euclidean distance can be used as distance metric.

Given a certain batch size *B* of samples on which the loss is meant to be computed, let *B_+_* be the number of sick samples and *B_-_* the number of healthy control samples. Let *B_T_* = *min*{*B*_+_, *B*_-_}. From a batch of size *B*, we randomly sample *B_T_* sick samples and *B_T_* control samples without repetitions (this implies that the smaller set of samples, either sick or control, will be fully considered). These 2*B_T_* samples constitute the *anchors* of the triplet margin loss. We represent the anchors with two sets: 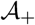 (which contains *B_T_* sick anchors) and 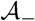 (which contains *B_T_* control anchors). For each anchor sample in 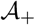 and 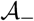, we sample: (a) a subject of the same class (addressed in this context as *positive*) and (b) a sample of the opposite class (addressed as *negative*). In our implementation, this is obtained by shuffling the samples in the opposite-class anchor set. This allows to constitute a set of positive samples 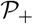 (i.e. of the same class) and a set of negative samples 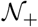 (i.e. of the opposite class) for the sick anchors 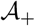 Analogously, a set of positive samples 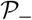 and a set of negative samples 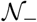 are obtained for the control anchors 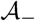.

Samples contained in the 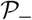 set are healthy control subjects, while samples contained in 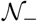 belong to sick ones. Analogously, samples contained in the 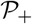 set belong to sick subjects, while samples contained in 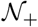 come from heathy control ones. This is because, in the context of the triplet margin loss, *positive* and *negative* mean of the *same* and of *opposite* ground truth class, respectively.

We define the triplet margin loss as:

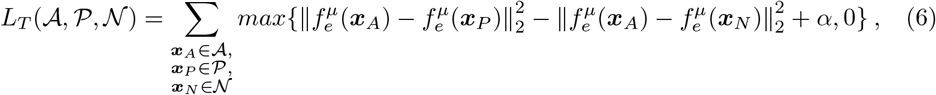

where *α* is a tunable margin which we set to 1 and 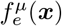 is the MLP that computes the mean of 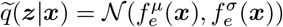 (see Section Implementation details). It follows that, for our 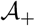 and 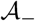 anchors sets, the triplet loss objective can be written as:

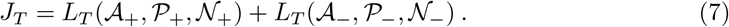

Intuitively, the first term of the right hand side of Eq 7 encourages the encodings of sick samples to be closer to each other in their K-dimensional Euclidean space and far away from the encodings of healthy control samples. In the same fashion, the second term of the equation encourages the encodings of healthy samples to be clustered in the same region of the latent space and to be distant from the encodings of sick samples.

With the definition of the triplet margin loss objective of Eq 7, we can extend the MVIB objective presented in Eq 5 and introduce the MVIB-T objective:

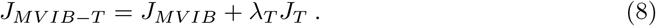

*λ_T_* is a multiplying constant which we set to 1 for all experiments.

### Full multimodal objective

For the training of MVIB, we adopt the same training paradigm proposed for the Multimodal Variational Autoencoder [41]. The MVIB objective presented in Eq 5 assumes that all *M* data modalities are present. This has the unfortunate consequence of not training the single-modality encoders 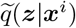 individually. This implies that the model cannot be used if certain data modalities are not available at test time. In order to circumvent this limitation and allow the MVIB to work at test time with missing data modalities, we need to compute the MVIB objective for the combined data modalities, as well as for the individual modalities.

Eq 5 can be reformulated as a full multimodal objective, which allows the model to be optimal in all multimodal and single-modality settings:

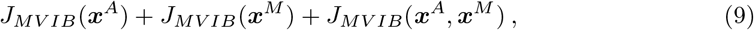

where ***x^A^*** represents the species-relative abundance profiles and ***x^M^*** the strain-level marker profiles. This extension holds also for the MVIB-T objective of Eq 8. For all experiments performed in this work, the objective functions are computed in their full multimodal form, as described in this section.

### Validation framework and performance evaluation

With the goal of providing a non-biased estimate of the model’s classification performance, we implemented a performance evaluation scheme inspired by DeepMicro [17]. We split each dataset into training and test sets with a 8:2 ratio. In particular, for the IBD, EW-T2D, C-T2D, Obesity, Cirrhosis and Colorectal datasets, we implemented the random train-test split using the same random partition seeds used by DeepMicro; this ensures that our test samples are the same ones considered by DeepMicro and allows a fair benchmark of the models’ performances. The same procedure was adopted for Obesity-Joint, Colorectal-EMBL, Early-Colorectal-EMBL, Hypertension and Colorectal-Metabolic too.

Considering only the training set, we performed a stratified 5-fold cross-validation and used the validation sets to compute a validation ROC AUC score for selecting the epoch with the best model parameters. The five best models obtained via the 5-fold cross-validation were then tested on the left-out test set and their predictions were ensembled via a majority vote. This procedure was repeated five times, each time with a different random partition seed, in order to ensure that the five experiments were conducted with independent random training-test splits. The resulting test performance metrics coming from the five independent experiments were then averaged and their mean was used for comparing model performance.

### Transfer learning

Motivated by the hypothesis that altered microbiome states caused by two different diseases might in fact share some common patterns, we performed experiments following a transfer learning paradigm [26, 27]. Iteratively, we first select a target disease from the set of considered datasets. This allows to define a target domain 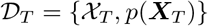, where 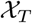 is the target feature space and *p*(***X***_*T*_) is the marginal distribution of the set of samples of the target dataset 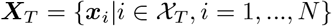. A target task 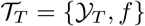 is also defined, where 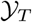 is the label space of the target dataset and *f* the decision function which is expected to be learned. The target task shall therefore be interpreted as the prediction of the disease which we mostly care about.

Merging the non-target datasets, we constitute a source domain and task 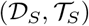. For the source task, all sick subjects are treated equally and they are assigned a positive ground truth label, independently on what pathology they actually have. We first train MVIB on the source domain and task 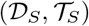 and fine-tune it on the target domain and task 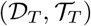.

For the transfer learning experiments, we adopted the *joint* datasets collection, as the microbial species and strain markers share the same positional indexes across the various datasets and present the same dimensionality.

As Colorectal-EMBL is an extension of the Colorectal dataset, and Obesity-Joint is an extension of Obesity, applying the procedure described above would allow the model to observe the samples shared by more than one dataset during both the source and the target task. This would lead to biased results. Therefore, in the transfer learning experiments, ΔObesity and ΔColorectal are considered during the source task instead of Obesity-Joint and Colorectal-EMBL, respectively.

### Cross-study generalisation

In order to further evaluate the generalisation capabilities of MVIB across different studies, and motivated by the fact of having various datasets for the same disease, we performed cross-study generalisation experiments. First, we identified the following ordered pairs of datasets: (EW-T2D, C-T2D), (C-T2D, EW-T2D), (ΔObesity, Obesity), (ΔColorectal, Colorectal), (Obesity, ΔObesity), (Colorectal, ΔColorectal). Then, for each pair, we trained MVIB on the first source dataset, and tested it on the second target one. No fine-tuning was performed on the test dataset.

### Explaining predictions with saliency

Following the same approach proposed by [25], given a multimodal pair of feature vectors 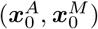, where ***x^A^*** represents the species-relative abundance profile and ***x^M^*** the strain-level marker profile, we would like to rank the strain-level markers of ***x^M^*** and the microbial species of ***x^A^*** based on their influence on the MVIB prediction. More formally, we compute the derivate of the MVIB class prediction with respect to both input vectors ***x^A^*** and ***x^M^***:

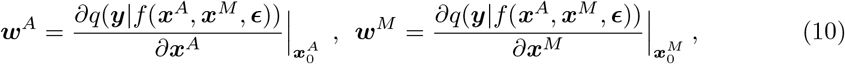

where *q*(·) is the parametric probabilistic decoder of MVIB and *f*(·) represents its whole multimodal encoding block, following the same notation of Eq 5.

The magnitude of the derivatives presented in Eq 10 indicates which strain-level markers and species-relative abundances need to be changed the least to affect the class score the most. For each disease dataset, we compute the saliency maps for all samples correctly classified as positive (i.e. sick) and compute their average. This allows to discover a ranking of the most influential strain-level markers and microbial species for each disease dataset (see Fig 1F,G). The computation of saliency maps is extremely quick and it only requires a single backpropagation pass.

### Trimodal MVIB: combining metabolomics and metagenomics

In order to further investigate the multimodal learning capabilities of MVIB, we performed trimodal experiments on the Colorectal-Metabolic dataset [37]. This dataset includes three data modalities for all samples: species-relative abundance, strain-level marker and metabolite profiles. For training the model in the trimodal setting, the same full multimodal training paradigm presented in Section Full multimodal objective was adopted. This allows to compute the MVIB objective for all possible data modalities combinations, as well as for the individual modalities.

In this trimodal setting, the objective function is computed in the following fashion:

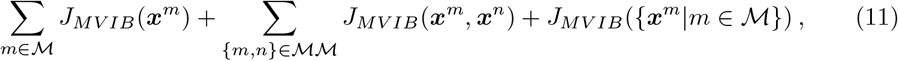

where 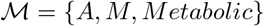 is the set of the three considered data modalities and 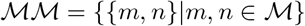 is the set of all possible (unordered) modality pairs. S1 Fig depicts the trimodal architecture of MVIB used for the experiments on the Colorectal-Metabolic dataset.

For the trimodal experiments, the learning rate has been set to 10^-5^ and the bottleneck dimension *K* to 128, as we observed that this slightly improves learning.

## Results

### MVIB achieves competitive results on the multimodal microbiome-based disease prediction task

We assess the performance of MVIB in comparison to existing state-of-the-art methods for microbiome-based disease prediction, i.e. Random Forest, DeepMicro [17] and PopPhy-CNN [19]. In Table 2, we report the AUC ROC values for our benchmark analysis on various disease cohorts. The results of MVIB are shown for both dataset collections *default* and *joint* (which we describe previously in Section Pre-processing). MVIB results are derived from the two input feature modalities, species abundances and strain-level markers (A+M). A complete summary of MVIB results for all bimodal dataset collections is available in S2 Table.

**Table 2.** Classification performance of MVIB and competing methods.

| Dataset | Random Forest |  |  | DeepMicro<sub>VAE</sub><sup>‡</sup><br>[17] |  |  | PopPhy-CNN<br>[19] | MVIB (A+M) |  |
| --- | --- | --- | --- | --- | --- | --- | --- | --- | --- |
|  | A | M | A+M | A | M | A+M | A | D | J |
| IBD | 0.899<br>(0.037) | 0.932<br>(0.025) | 0.936<br>(0.017) | 0.779<br>(0.032) | 0.899<br>(0.039) | 0.833<br>(0.034) | 0.799<br>(0.052) | 0.922<br>(0.020) | **0.936**<br>**(0.014)** |
| EW-T2D | 0.825<br>(0.020) | 0.812<br>(0.017) | 0.789<br>(0.024) | 0.640<br>(0.051) | 0.853<br>(0.041) | 0.741<br>(0.080) | 0.591<br>(0.037) | **0.859**<br>**(0.023)** | 0.853<br>(0.025) |
| C-T2D | 0.717<br>(0.018) | 0.736<br>(0.023) | 0.734<br>(0.017) | 0.715<br>(0.031) | 0.719<br>(0.019) | 0.714<br>(0.030) | 0.641<br>(0.019) | 0.750<br>(0.009) | **0.758**<br>**(0.012)** |
| Obesity | 0.662<br>(0.022) | 0.634<br>(0.019) | 0.619<br>(0.017) | 0.600<br>(0.030) | 0.599<br>(0.014) | 0.576<br>(0.040) | 0.630<br>(0.018) | 0.662<br>(0.024) | **0.666**<br>**(0.027)** |
| Cirrhosis | 0.897<br>(0.013) | 0.896<br>(0.007) | 0.905<br>(0.009) | 0.781<br>(0.021) | 0.891<br>(0.016) | 0.867<br>(0.018) | 0.893<br>(0.008) | **0.925**<br>**(0.005)** | 0.924<br>(0.005) |
| Colorectal | 0.803<br>(0.048) | 0.771<br>(0.053) | 0.764<br>(0.053) | 0.739<br>(0.070) | 0.737<br>(0.068) | 0.548<br>(0.040) | **0.803**<br>**(0.023)** | 0.780<br>(0.071) | 0.777<br>(0.069) |
| Obesity-Joint | 0.806<br>(0.026) | 0.795<br>(0.024) | 0.799<br>(0.025) | 0.662<br>(0.015) | 0.637<br>(0.021) | 0.600<br>(0.017) | 0.697<br>(0.026) | 0.815<br>(0.019) | **0.818**<br>**(0.018)** |
| Colorectal-EMBL | **0.890**<br>**(0.015)** | 0.866<br>(0.024) | 0.860<br>(0.016) | 0.662<br>(0.027) | 0.725<br>(0.049) | 0.693<br>(0.034) | † | 0.811<br>(0.010) | 0.814<br>(0.013) |
| Early-Colorectal-EMBL | 0.577<br>(0.046) | 0.533<br>(0.039) | **0.582**<br>**(0.024)** | 0.562<br>(0.070) | 0.557<br>(0.048) | 0.5616<br>(0.048) | † | 0.535<br>(0.050) | 0.543<br>(0.048) |
| Hypertension | 0.662<br>(0.035) | **0.677**<br>**(0.023)** | 0.631<br>(0.031) | 0.523<br>(0.030) | 0.561<br>(0.028) | 0.658<br>(0.054) | † | 0.603<br>(0.045) | 0.591<br>(0.041) |

For Random Forest and DeepMicro, we report the results on various feature combinations, i.e. only species abundance (A), only strain-level markers (M) and the concatenation of abundances and markers (A+M). Benchmark methods do not explicitly model multiple input data modalities, hence we adopted a simple feature concatenation in the (A+M) setting. The results of Random Forest shown in Table 2 are derived after fine-tuning the model through cross-validated grid-search over the hyperparameter space summarised in S3 File. For PopPhy-CNN, we only report the results using species abundances (A) modality, since the current implementation of the method does not scale to include strain-level markers.

The DeepMicro method consists of a two-steps mechanism: (1) train an autoencoder to generate input embeddings, (2) train a downstream classifier on the embeddings computed in the first step. Various autoencoder architectures and downstream classifiers are introduced in [17] and there is no clear criteria on which combination has an optimal performance. For our benchmark analysis, we trained DeepMicro with the Variational Autoencoder (VAE) [28]. We believe this choice offers fair comparisons, since MVIB is also based on variational inference. Choosing VAE for DeepMicro allows us to highlight the key improvements of our end-to-end multimodal architecture when compared to the demanding two-step mechanism offered by DeepMicro. For the classification step of DeepMicro, we trained multiple downstream classifiers, i.e. SVM, MLP and Random Forest, then we report the best test ROC AUC values.

We modified the source code of PopPhy-CNN to make the model validation and testing consistent with our validation framework. The original PopPhy-CNN implementation offers a slightly different validation procedure, hence we made the necessary changes to ensure that all of the evaluation results (see Table 2) are computed in the same manner.

From the results of Table 2, we see that MVIB outperforms Random Forest on all datasets except Colorectal-EMBL, Early-Colorectal-EMBL and Hypertension. MVIB consistently outperforms DeepMicro on all datasets in both settings, i.e. single-modality (A or B) and multimodality (A+M). In comparison to PopPhy-CNN, MVIB achieves better results on all datasets, except for Colorectal, where PopPhy-CNN achieves 2% higher ROC AUC. For the Colorectal-EMBL, Early-Colorectal-EMBL and Hypertension datasets, we could not produce PopPhy-CNN results, as the original implementation seems to generate an infinite loop when the phylogenetic tree is pruned.

Early-Colorectal and Hypertension present the hardest diseases to predict for all classifiers including MVIB. MVIB achieves approximately 55% and 60% ROC AUC on each of the aforementioned datasets, respectively. However, these values are still above random baseline. In summary, the discrimination capabilities of various classifiers including MVIB vary among different datasets which may indicate less measurable microbial changes in subjects with certain diseases.

### Multimodal ablation study

To further evaluate the multimodal learning capabilities of MVIB, we perform an ablation study to compare classification performance of single-modality and bimodal settings. To this end, we train MVIB by optimising the full multimodal objective (see Section Full multimodal objective), which ensures that all encoders are trained individually as well as jointly. At test time, for the single-modality setting, we tested the model by considering either species abundances (MVIB-A) or strain-level markers (MVIB-M) as inputs. While for the bimodal setting, we passed both data modalities simultaneously as inputs to our model (MVIB-A+M). Fig 2 summarises our results.

**Fig 2.**
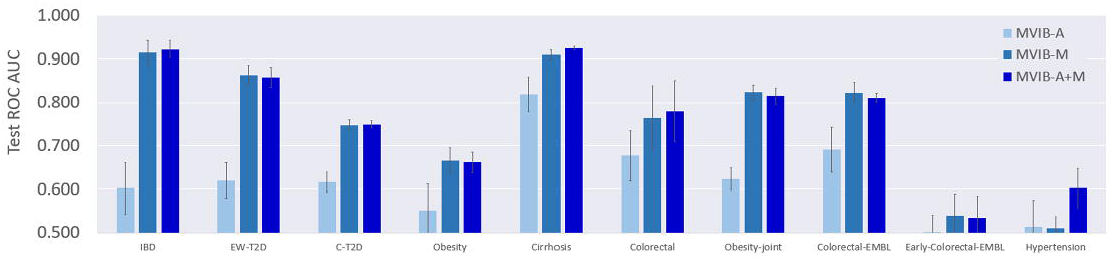
Ablation study results. Comparisons between single-modality and bimodal MVIB. The shown values are ROC AUC on test sets, error bars represents standard error computed by repeating each experiment five times on different random train/test splits. We leveraged the *J_MVIB_* objective (Eq 5) for optimisation. MVIB-A indicates the model performance only on species-relative abundances at test time. MVIB-M indicates the model performance only on strain-level markers at test time. MVIB-A+M indicates the model performance on both species-relative abundances and strain-level markers at test time.

Fig 2 shows that MVIB-M results are consistently better than those obtained from MVIB-A. In the bimodal setting, MVIB-A+M, the performance remains comparable to MVIB-M results (i.e. the best performance). One can notice that on the datasets of IBD, Cirrhosis, Colorectal and Hypertension, the results of MVIB-A+M are (slightly) better than the results reported by MVIB-A or MVIB-M.

MVIB can efficiently combine heterogeneous input data modalities. Although ROC AUC results from MVIB-A are consistently lower than the ROC AUC values from MVIB-M, the combination of the two modalities does not lead to a performance drop that may occur in other methods due to an increased feature space, i.e. the curse of dimensionality. In summary, MVIB can guarantee classification performance which is at least as good as the best single-modality performance.

### The triplet margin loss can improve classification

The first term of the original MVIB objective *J_MVIB_* (Eq 5) is a negative log-likelihood, which acquires the shape of a binary cross-entropy in the binary classification setting. We explored an extension of this objective, namely *J_MVIB–T_*, by adding a triplet margin loss term (Eq 8).

The triplet margin loss aims to encourage the latent distribution of samples which belong to the same class to cluster in a dedicated region of the latent Euclidean space. At the same time, the triplet margin loss encourages the distributions of samples of different classes to depart from each other in the latent space. This aims at increasing the separability of different classes and facilitating the classification task.

S4 Table presents a comparison of the effects of the triplet margin loss on the MVIB classification performance. For the IBD, EW-T2D, C-T2D and Early-Colorectal-EMBL datasets, the highest ROC AUC is achieved by optimising the *J_MVIB–T_* objective, which includes the triplet margin loss. The *J_MVIB_* objective leads to best results on the remaining datasets.

Fig 3A depicts the 95% confidence interval of the subjects’ stochastic encodings 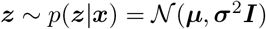 deriving from the optimisation of the *J_MVIB_* (Eq 5). Such Gaussian distributions are the output of the PoE (Eq 4) and consist in fact in the joint posterior distribution of the latent encoding, conditioned on all input data modalities. In comparison with the curves of Fig 3C, obtained with the *J_MVIB–T_* objective (Eq 8), we observe that the triplet margin loss leads to Gaussian distributions which present a higher variance. Conversely, stochastic encodings deriving from *J_MVIB_* (Eq 5) present smaller variance (see Fig 3A).

**Fig 3.**
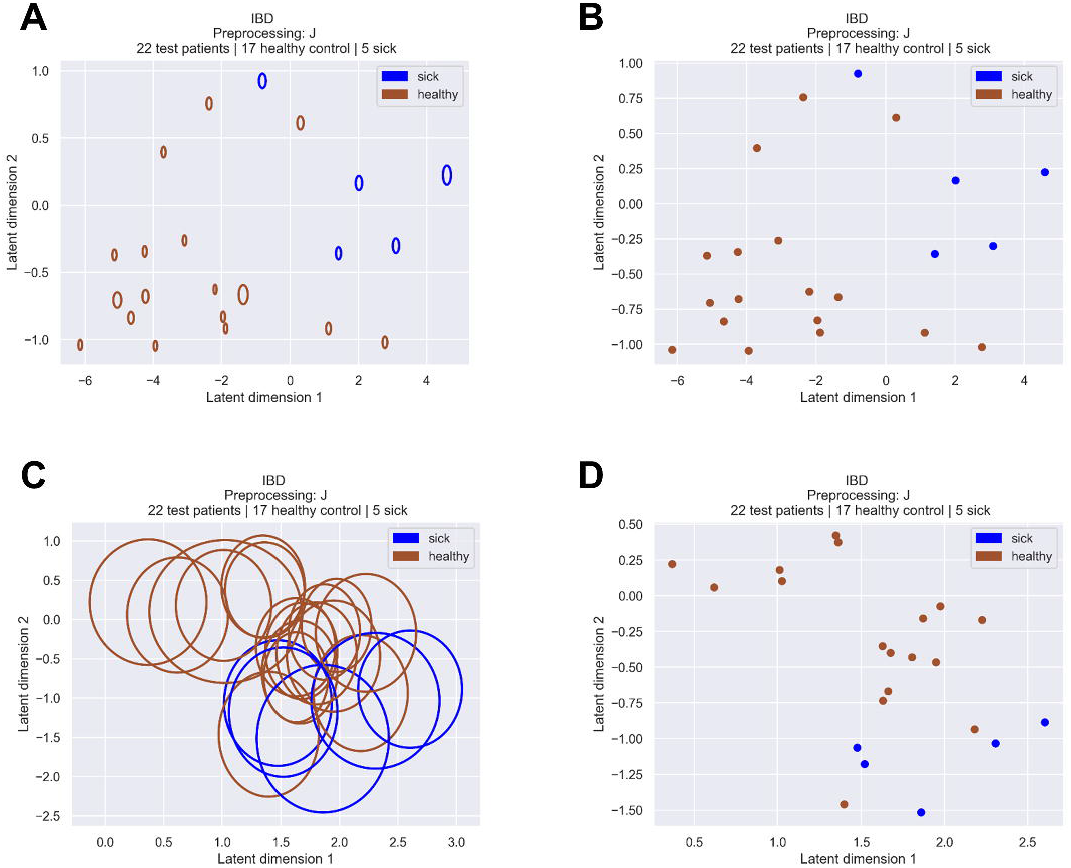
Effect of the triplet margin loss on the stochastic encodings of the microbiome samples. The depicted curves are the 95% confidence intervals of the samples’ stochastic encodings 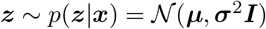; the points are their means ***μ***. The displayed encodings consist in a set of 22 test samples obtained from a random training-test split of the IBD dataset (i.e. the 20% of the dataset not used for training). The *K* dimension of the latent space has been set to 2 in order to allow a 2D visualisation. (A, B) optimisation of the *J_MVIB_* objective (Eq 5). (C, D) optimisation of the *J_MVIB–T_* objective (Eq 8).

Methods such as simple autoencoders, or the PCA, which are commonly used for dimensionality reduction, provide deterministic encodings, i.e. compressed representations of the input without a rigorous estimate of the uncertainty. Our method allows to compute *stochastic* encodings, i.e. to represent the input not only as a point in a latent space, but as a probability distribution. Computing probability distributions allows to estimate the confidence of the stochastic encodings.

Intuitively, it is preferable to obtain stochastic encodings which present a small variance, as this allows for a better separability and classification. Fig 3 allows to visually interpret the effect of including the triplet margin loss (Eq 8) in the objective function and to visualise the results of the different losses on the samples’ stochastic encodings. We conclude that, although adding the triplet margin loss can lead to better classification results on some datasets (see S4 Table), the stochastic encodings derived from the optimisation of the *J_MVIB–T_* objective (Eq 8) present higher variance with respect to those obtained from the optimisation of the *J_MVIB_* objective (Eq 5).

In addition to the plots of Fig 3, which only refer to the IBD dataset, S5 File contains the plots of the MVIB stochastic encodings for all the datasets considered in this work. The stochastic encodings depicted in S5 File are obtained from the *joint* datasets collection. They are available for models trained by optimising the *J_MVIB–T_* objective, as well as models trained by optimising the *J_MVIB_* objective, in order to allow comparison.

S6 File presents for each dataset a comparison between the PCA 2D projections and the mean of the MVIB 2D stochastic encodings. For both the PCA and the MVIB plots presented in S6 File, the *default* datasets collection has been adopted. It is possible to observe that the PCA projections completely fail at clustering the two samples classes (healthy and sick) in dedicated areas of the 2D plane. Conversely, MVIB encodes sick and healthy samples in two clearly separated areas of the 2D plane, creating two distinct clusters.

### Transfer learning across different diseases can improve disease prediction

Driven by the hypothesis that different diseases might lead to some common altered patterns in the subjects’ microbiome, we performed various transfer learning experiments. As described in Section Transfer learning, we first pre-trained MVIB on all non-target diseases, then we fine-tuned the model on target diseases. For these experiments, we only consider the *joint* datasets collection, as it presents feature vectors with the same dimensions across all disease cohorts for both the species-relative abundance and strain-level marker profiles. S7 Table presents the MVIB classification performance achieved by performing a pre-training on the source task followed by a fine-tuning on the target diseases. The ROC AUC slightly decreases on the Cirrhosis and Colorectal target datasets. A consistent performance drop is observed on the IBD and EW-T2D target datasets. Improvements on randomly initialised models can be observed on the remaining datasets, i.e. C-T2D, Obesity and Hypertension.

We did not consider Colorectal-EMBL or Obesity-Joint datasets here, since the former is just an extension to Colorectal dataset, and the latter is an extension to Obesity dataset. Thus, we ensured that the samples that are present in more than a single dataset will not observed twice, i.e. during both source and target tasks. Moreover, ΔObesity and ΔColorectal are only used during the source task (i.e. pre-training).

### Cross-study generalisation results and benchmark

Generalisation is a fundamental requirement for machine learning models. It consists in the capacity to make correct predictions on samples which were not observed at training time. In order to further investigate how well MVIB can generalise, we performed cross-study experiments as described in Section Cross-study generalisation. First, six ordered pairs of datasets were identified: (EW-T2D, C-T2D), (C-T2D, EW-T2D), (ΔObesity, Obesity), (ΔColorectal, Colorectal), (Obesity, ΔObesity), (Colorectal, ΔColorectal). The first dataset of each pair was exclusively used for training, while the second one for testing without fine-tuning. In order to guarantee that abundance and marker features share the same positional indexes across the source and target datasets, the *joint* dataset collection was employed for all cross-study experiments (see Section Pre-processing).

In order to compare the generalisation capabilities of MVIB with a state-of-the-art machine learning model used in microbiome research, we performed benchmark experiments with Random Forest. The Random Forest was trained on the concatenation of the abundance and marker profiles (multimodal setting). For the Random Forest, a cross-validated grid-search over a defined hyperparameter space was adopted for fine tuning (see S3 File). MVIB did not undergo hyperparameter tuning, but employed the default implementation described in Section Implementation details, with the exception of a lower learning rate, which was set to 10^-5^, as we observed this improves convergence.

Fig 4 shows the cross-study results. MVIB performed better than the Random Forest on the (EW-T2D, C-T2D) and (ΔColorectal, Colorectal) experiments. Conversely, the Random Forest achieved better generalisation on (C-T2D, EW-T2D), (ΔObesity, Obesity) and (Obesity, ΔObesity). The two methods obtained the same results on (Colorectal, ΔColorectal).

**Fig 4.**
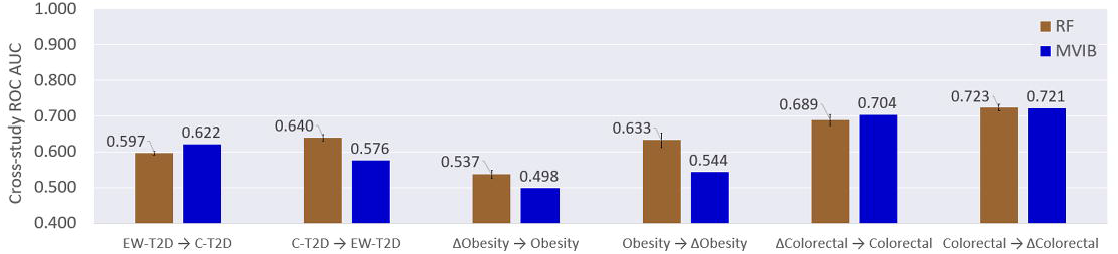
MVIB cross-study generalisation and benchmark. Values are test ROC AUC computed by first training the models on a source dataset and then testing it on a target dataset. RF: Random Forest. All datasets used for cross-study experiments belong to the *joint* collection. For MVIB, the *J_MVIB–T_* objective has been adopted for the optimisation (Eq 8). The datasets reported on the x-axis shall be interpreted as: *train →test.* For the Random Forest, the error bars represent the standard error over five repeated experiments and account for the stochasticity of the Scikit-learn implementation. The standard error is missing for the MVIB results, as our PyTorch implementation has been made deterministic.

Based on the empirical results which we obtained, it is possible to conclude that MVIB offers competitive cross-study generalisation when benchmarked with a fine-tuned Random Forest.

### Discovering the most salient microbial species and strain-level markers

In order to achieve interpretability, we implemented a method which allows to compute saliency, i.e. detecting the areas of the input vectors which are most discriminative with respect to the predicted class (see Section Explaining predictions with saliency). For each disease dataset in the *default* dataset collection, we obtained saliency maps of both abundance and marker profiles. Such saliency maps were obtained by computing the derivative of the model’s positive predictions with respect to the inputs (Eq 10). Since we were mostly interested in the magnitude of such gradients, rather than the sign, the absolute value was considered.

The aim of this analysis is the discovery of the most salient microbial species and strain-level markers for each considered dataset, i.e. the features which mostly affect the outcome of positive disease predictions. In order to discover true biological insights about the relationships between the microbiome and the analysed diseases, we considered saliency maps derived from true positive predictions. After having trained MVIB, each dataset was passed through the model to compute predictions for all samples. Saliency maps were then computed (Eq 10). Saliency vectors coming from true positive predictions where then extracted from the dataset batch. As described in Section Validation framework and performance evaluation, experiments were repeated five times with different independent train-test splits. Furthermore, for each of the five train-test splits, the five different best models derived from a 5-fold cross-validation were ensembled. This leads to 25 different models, and therefore 25 saliency maps, which were then averaged.

Fig 5 depicts the microbial species sorted by mean saliency for the Colorectal-EMBL dataset. Fig 5A only depicts the top 25 species, while Fig 5B depicts the full distribution.

**Fig 5.**
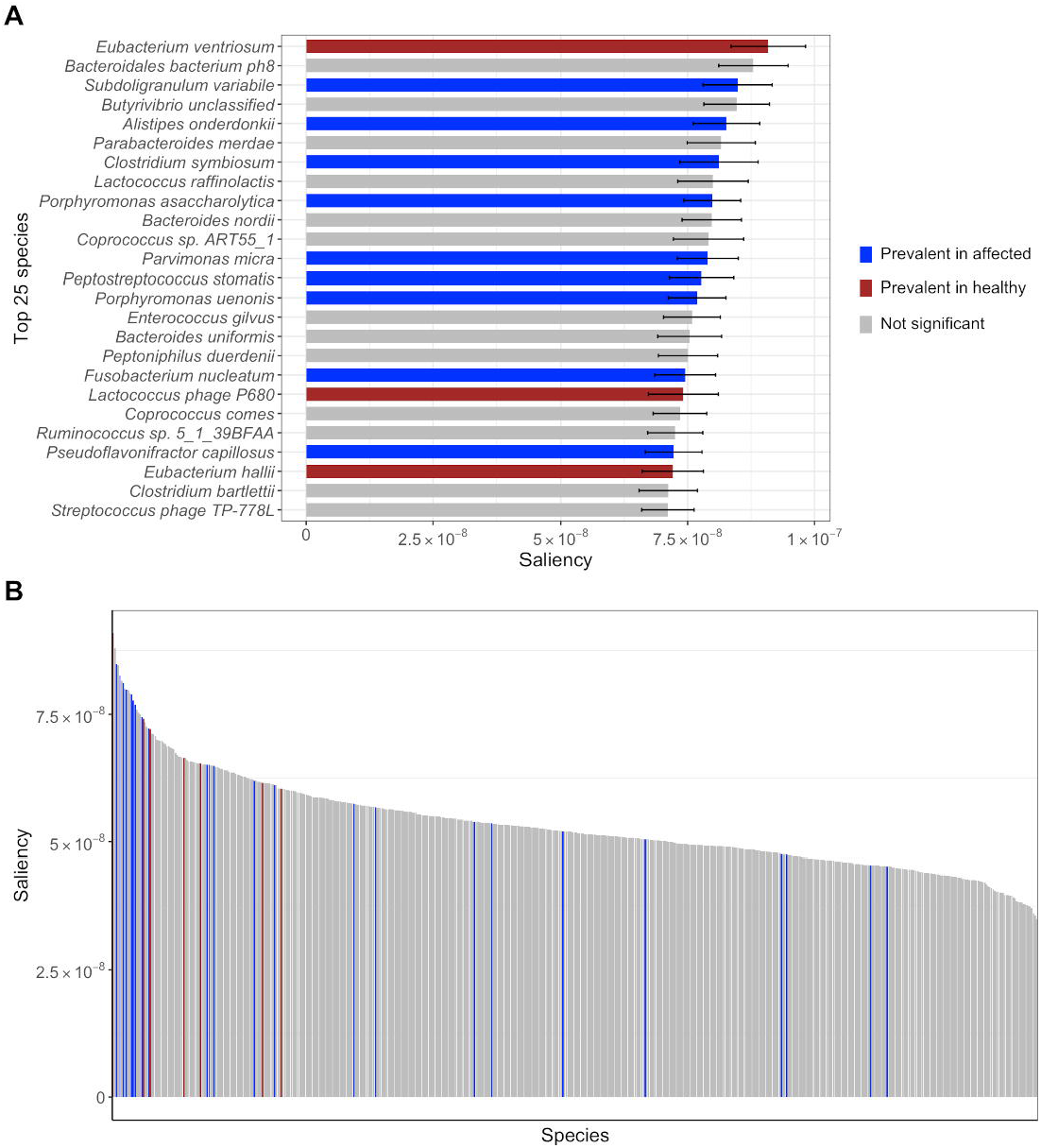
Microbial species sorted by saliency for the Colorectal-EMBL dataset. Saliency maps computed for the Colorectal-EMBL dataset using Eq 10. (A) Top 25 microbial species sorted by mean saliency. Species prevalence in healthy (red) and affected (blue) individuals were calculated using a Wilcoxon test for each microbial species for two unpaired samples: healthy and affected individuals. Error bars represent the standard error over the five repeated experiments and the five ensembled MVIB models for each experiment. (B) Full distribution of the mean saliency across the microbial species of the Colorectal-EMBL dataset.

For the microbial species, the prevalence in healthy and affected individuals was calculated by means of a Wilcoxon test. For each microbial species, two samples were constructed from the abundance values: (1) healthy individuals, (2) affected individuals.

False discovery rate (FDR) correction was applied to adjust for multiple hypotheses testing. Species prevalence supported by adjusted p-values <0.1 was reported as significant. In Fig 5, species which significantly prevail in healthy individuals are marked in red, while prevalent specie in positive individual are marked in blue.

S8 File contains the plots with the top 25 microbial species and strain markers sorted by mean saliency for all datasets considered in this work, analogous to what Fig 5A depicts for the Colorectal-EMBL dataset.

As depicted in Fig 5 and S8 File, many species with significant prevalence in either healthy (red) or affected individuals (blue) appear among the most salient ones. However, in multiple instances, non-prevalent species (grey) are also listed among the most salient ones. We believe this has the following explanation. On one hand, the Wilcoxon test used to compute the significance of species prevalence considers one single species at a time. Hence, the Wilcoxon test is not capable to capture complex multi-dimensional patterns in the abundance distribution. On the other hand, MVIB can learn complex non-linear dependencies between the input and a given label. As non-prevalent microbial species appear be salient for the model, this seems to point out the existence of inter-dependencies among microbial species, which MVIB could capture. Additionally, in the Wilcoxon test, p-values < 0.1 were reported as significant. The selection of this threshold for the p-values intrinsically affects what species are labelled as prevalent.

S9 File contains the histogram plots of the full saliency distributions over species (analogous to what Fig 5B depicts for the Colorectal-EMBL dataset), as well as the violin plots of the saliency distributions grouped by prevalence. The species in the histograms are sorted by mean saliency. As it is possible to observe in the histograms plots, saliency maps are not sparse. This is possibly due to how they are computed: as described above, the final saliency maps are obtained by averaging the saliency computed from 25 models. Additionally, as displayed in the plots, values are very low. Saliency maps are in fact the gradients of the model output with respect to the input. In deep learning, gradients are in general very low to allow computational stability, but also for theoretical reasons (e.g. regularization). Although we cannot escape these drawbacks which accompany gradients, we still observe that the average saliency maps let species emerge, which are under-/over-represented in the affected and healthy individuals. We can conclude that, although models e.g. RF or penalised linear regression could allow for simpler and more intuitive feature selection, deep-learning-based methods like MVIB can also allow for explainable predictions.

### Metabolomics can improve colorectal cancer prediction when combined with metagenomics

With the aim of further investigating the multimodal learning capabilities of MVIB, we performed experiments on the Colorectal-Metabolic dataset extracted from [37]. For each sample, this dataset presents three modalities: species-relative abundance, strain-level marker and metabolite profiles.

First, we trained MVIB following the full multimodal training paradigm presented in Section Trimodal MVIB: combining metabolomics and metagenomics. This ensures that all encoders are individually and jointly trained and that MVIB can perform single-modality and multimodal predictions at test time. We then tested the model in various multimodal and single-modality fashions: abundance profiles only (A), marker profiles only (M), abundance + marker profiles (A+M), abundance + marker + metabolite profiles (A+M+Metabolic).

We compared MVIB with a fine-tuned Random Forest. For fine-tuning the Random Forest, a cross-validated grid-search over a defined hyperparameter space was used (see S3 File). In the A+M and A+M+Metabolic multimodal settings, the various input modalities have been concatenated and then fed into the Random Forest.

Fig 6 shows the obtained experimental results. It is possible to observe that the multimodal settings A+M and A+M+Metabolic allow MVIB to reach higher test ROC AUC with respect to Random Forest. Furthermore, adding metabolomic data (A+M+Metabolic) allows to achieve the best classification results. Results obtained in the single-modality M setting are comparable to those obtained in the trimodal setting.

**Fig 6.**
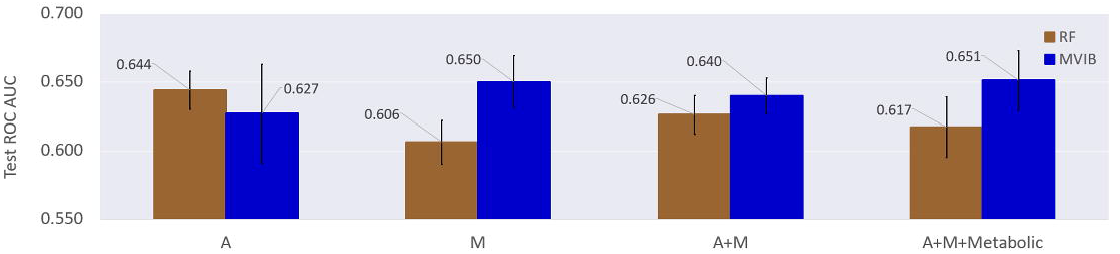
Comparison of different multimodal and single-modality models for colorectal cancer prediction. Values are test ROC AUC, error bars are standard error obtained by repeating the experiment five times on different random train/test splits. RF: Random Forest. A: species-relative abundance profiles. M: strain-level marker profiles. Metabolic: metabolite profiles. For MVIB, the *J_MVIB_* objective has been adopted for the optimisation (Eq 5). Experiments are executed on the Colorectal-Metabolic dataset extracted from [37].

The Random Forest performs better than MVIB only in the abundance-only single-modality setting (A). Most notably, the Random Forest performance tends to degrade when further data modalities are added. This shows the superiority of MVIB in combining multiple heterogeneous data modalities.

### Empirical analysis of the training time

In this section, we report the analysis of the training time of various machine learning models trained on the trimodal Colorectal-Metabolic dataset from [37]. We choose this dataset since it has three data modalities, and the highest number of samples compared with the other datasets considered in this work.

For benchmarking, we compared the training time of MVIB with the training time of Random Forest (RF-DEF in Fig 7), Support Vector Machine (SVM-DEF in Fig 7) and Random Forest with hyperparameter optimisation (RF-HPO in Fig 7). For both RF-DEF and SVM-DEF, we considered the default implementation from Scikit-learn [46] library version 0.23.2, which runs on CPU. We labelled the two methods with *-DEF* suffix in order to highlight that the default Scikit-learn implementation was used without any hyperparameter optimisation.

**Fig 7.**
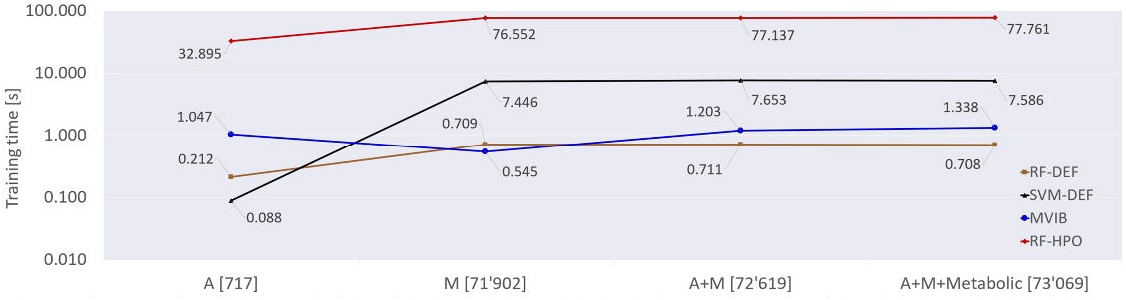
Training time comparison across different machine learning models and various feature space dimensions. The values reported in this graph consist in the training time measured in seconds. The Colorectal-Metabolic dataset extracted from [37] has been used for training all models. The depicted training times are obtained by averaging the run times of five experiments with different random train/test splits. A: species-relative abundance profiles. M: strain-level marker profiles. Metabolic: metabolite profiles. RF-DEF: Random Forest with default Scikit-learn implementation. SVM-DEF: Support Vector Machine with default Scikit-learn implementation. RF-HPO: Random Forest with hyperparameter optimisation (see S3 File). For MVIB, the *J_MVIB_* objective has been adopted for the optimisation (Eq 5). On the x-axis, next to the modality name, the feature space dimension is reported in square brackets.

For RF-HPO, we implement a cross-validated grid-search over a defined hyperparameter space (see S3 File). We used 16 CPU cores to parallelise the grid-search. The hyperparameter optimisation adopted for RF-HPO is the same one which we used to obtain the results in Table 2.

For MVIB, we set the learning rate to 10^-4^ and the latent dimension *K* to 256. We trained MVIB with a batch size of 256 for up to 50 epochs, as convergence (i.e. best validation ROC AUC) is normally observed within the first 20 epochs. We considered the training complete at the epoch in which the model achieves the highest ROC AUC on the validation set. No ensembling was performed, i.e. a single model was trained in this set of experiments. MVIB experiments were executed on a single NVIDIA GeForce RTX 2080 Ti GPU.

We performed experiments in various multimodal and single-modality fashions: abundance profiles only (A), marker profiles only (M), abundance + marker profiles (A+M), abundance + marker + metabolite profiles (A+M+Metabolic). For RF-DEF, SVM-DEF and RF-HPO, in the multimodal settings, the input feature vectors of the various modalities were concatenated. Fig 7 depicts the empirical results of the training time analysis.

Although the SVM-DEF is the fastest model to complete training in the abundance-only setting (A), its training time dramatically increases in the markers-only setting (M), as well as in the the two multimodal settings (A+M and A+M+Metabolic), reaching an average training time of more than 7 seconds. The feature space dimensions in the various settings are reported in Fig 7. The RF-DEF exhibits the same trend: it achieves the fastest training in the A setting, but requires a slightly longer time in all other settings. Compared with the SVM-DEF, the RF-DEF is always much faster and never requires more than 1 second to complete training. The training of RF-HPO requires consistently more time due to the cross-validated grid-search: around 30 seconds in the A setting, and more than 70 seconds for all other settings.

For MVIB we observe a different behaviour. The training time does not appear to correlate with the feature space dimension. In fact the shortest training time is achieved in the M setting, which presents a much higher feature space dimension with respect to A. In all other settings, MVIB converges in around 1 second.

In the A setting, despite the smaller feature space dimension, MVIB requires a longer training time with respect to the M setting. This is because the training duration of MVIB is mainly determined by the number of epochs needed to reach the highest validation ROC AUC. The number of required epochs is mainly a function of the learning rate and the batch size. The single forward pass and backpropagation steps are extremely quick and highly parallelised, hence the feature space dimension plays a less relevant role in the training time of MVIB w.r.t. RF-DEF, SVM-DEF and RF-HPO.

It is additionally worth mentioning that RF-DEF is consistently worse than MVIB when it comes to classification results (see S10 Table and Table 2, respectively). RF-HPO provides competitive classification performance with respect to MVIB (see Random Forest results in Table 2), but its training time is consistently higher (see Fig 7). Hence, we can conclude that the Random Forest only competes with MVIB when it undergoes an extensive hyperparameter optimisation. Our implementation of MVIB requires very short training time and provides high classification performance without fine-tuning.

## Discussion

Microbiome-based disease prediction is a challenging task due to several reasons. First, microbial communities present high complexity in their composition. Second, microbial features (e.g. species abundance, strain-level markers and metabolites) are heterogeneous data modalities and sometimes they are generated using different technologies (e.g. shotgun metagenomic sequencing and mass spectrometry, respectively). Third, human gut microbiome is in a state of constant change, not only when a host is affected by a certain disease, but also as a function of lifestyle, e.g. diet [2], stress [47] and sleep [48]. Therefore, it is not surprising that a “healthy” microbiome can not be determined by simple rules of thumb, e.g., high or low count of individual taxa [49, 50].

In this work, we have introduced MVIB, a novel multimodal deep learning approach for microbiome-based disease prediction. MVIB computes a joint stochastic encoding of species-relative abundance and strain-level marker profiles. Both of these microbial features were obtained from shotgun metagenomic sequencing. MVIB stochastic encodings are maximally compressive of the various input data modalities and are simultaneously maximally informative about the output labels. We demonstrate that MVIB scales well in the trimodal setting where the learned encoding combines information from the species-relative abundances, the strain-level markers and the metabolites (from mass spectrometry). When the various input data modalities are considered jointly, MVIB computes a more complete representation of the host microbiome.

Our results show that MVIB competes with state-of-the-art methods for microbiome-based disease prediction, e.g. Random Forest, DeepMicro and PopPhy-CNN. MVIB achieves the highest ROC AUC on eight out of the eleven cohorts, although the improvement in performance can be marginal in certain cases. We also noticed that the discrimination capabilities of various classifiers including MVIB vary among different datasets, which may indicate less measurable microbial changes in subjects with certain diseases. Compared to DeepMicro, MVIB has the advantage of being an end-to-end approach, i.e. it learns a mapping from inputs to outputs and it does not require hyperparameter tuning. Conversely, DeepMicro requires a 2-step training approach, and it demands a complex fine-tuning process due to a lack of well-defined criteria about which autoencoder architecture and downstream classifier shall be used with a given dataset. Similarly, Random Forest requires a time-consuming hyperparameter optimisation step to achieve a good performance. Therefore, the training time of MVIB is ~70 times faster than Random Forest training time (including the hyperparameter optimisation).

Furthermore, we adopted a saliency technique derived from computer vision literature to interpret the output of MVIB and we identified the most relevant microbial species and strain-level markers to MVIB predictions. We performed cross-study generalisation experiments, where we trained and tested MVIB on different cohorts of the same disease. Our results show that MVIB is able to generalise in some cases like colorectal cancer, however, the results were not always satisfactory. Poor generalisation performance may be due to other environmental factors that are specific to each cohort which we do not consider in our model. For example, nutritional differences between Europe and China could play an important role in shaping the gut microbiome in the two different type 2 diabetes cohorts, i.e. EW-T2D and C-T2D.

Finally, one possible area for future work is to extend MVIB from microbiome-based disease prediction to microbiome-based disease prevention, i.e. to predict the presence of a certain disease in its early stages, e.g. by identifying specific microbiome patterns that indicate the potential for developing that disease. We will also consider extending the current implementation to accommodate temporal longitudinal multimodal microbiome data; the current architecture can support such an extension using for example Long Short-Term Memory networks (LSTMs) [51] as encoders to capture temporal dynamics.

## Acknowledgments

We would like to acknowledge Dr. Carolin Lawrence and Dr. Francesco Alesiani for valuable discussion.

## Supporting information

**S1 Fig. Trimodal MVIB architecture.** This figure depicts the trimodal architecture of MVIB used for combining species-relative abundance, strain-level marker and metabolite profiles for colorectal cancer prediction.

**S2 Table. Complete experimental results for the multimodal microbiome-based disease prediction task with MVIB.** Results obtained optimising the *J_MVIB–T_* objective (Eq 8). Experiments are executed five times with random independent training-test splits. Values in brackets refer to the standard error over the repeated experiments. All values in the table refer to metrics computed on the test sets. ROC AUC: area under the receiver operating characteristic curve. AC: classification accuracy. F1: F1 score. P: precision. R: recall. D and J refer to the two pre-processing techniques adopted and the two collections of datasets obtained: *default* (D) and *joint* (J).

**S3 File. Random Forest hyperparameter space for cross-validated grid-search.** The file contains the hyperparameter space considered for Random Forest in cross-validated grid-search.

**S4 Table. Comparison of different objective functions and pre-processing techniques.** All values are ROC AUC computed on the test sets. Values in brackets refer to the standard error over five repeated experiments. The first group of columns presents the results obtained optimising the *J_MVIB–T_* objective (Eq 8), which includes the triplet margin loss. The second group of columns presents the results obtained optimising the original objective function *J_MVIB_* (Eq 5). D and J refer to the two pre-processing techniques adopted and the two collections of datasets obtained: *default* (D) and *joint* (J).

**S5 File. Effect of the triplet margin loss on the stochastic encodings of the microbiome samples.** For all datasets considered in this work, this file presents plots of the 2D MVIB stochastic encodings analogous to Fig 3. The depicted curves are the 95% confidence intervals of the samples’ stochastic encodings 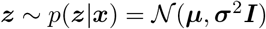; the points are their means ***μ***. The displayed encodings consist only in the test samples obtained from random training-test splits (i.e. the 20% of the dataset not used for training). The *K* dimension of the latent space has been set to 2 in order to allow a 2D visualisation. Plots derived from both the optimisation of the *J_MVIB–T_* objective (Eq 8) and the optimisation of the *J_MVIB_* objective (Eq 5) are included. Five copies of all plots are available, as they are obtained by training the model with five different independent train-test random splits.

**S6 File. PCA 2D projections and MVIB 2D stochastic encoding.** This file presents, for each dataset, the plots of the PCA 2D projections, as well as the plots of the mean of the MVIB 2D stochastic encodings. For the MVIB stochastic encodings 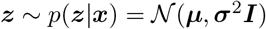, the depicted points represent the mean ***μ***. The *K* dimension of the latent space has been set to 2 in order to allow a 2D visualisation of the encodings. For training MVIB, the *J_MVIB–T_* objective (Eq 8) has been optimised. For MVIB, five copies of the means plots are available, as they are obtained by training the model with five different independent train-test random splits. Both the PCA and the MVIB plots have been created starting from the *default* datasets collection.

**S7 Table. Comparison of pre-trained models against randomly initialised models.** The first column presents the results obtained with randomly initialised models. The second column displays classification results obtained by first pre-training the models on all source datasets, and then fine-tuning them on the target disease (see Section Transfer learning). The *J_MVIB–T_* objective was adopted for both columns (Eq 8). J refers to the adopted pre-processing technique: *joint*. Reported values are ROC AUC computed on the test sets. In brackets, the standard error over five repeated experiments is reported.

**S8 File. Top microbial species and strain markers sorted by saliency for all datasets.** These files present the plots of the top 25 microbial species and strain markers for all datasets considered in this work, analogous to what Fig 5A depicts for the species from the Colorectal-EMBL dataset. Additionally, the scripts used to create the plots are included.

**S9 File. Saliency distributions over microbial species for all datasets.** For each dataset, two different kinds of plots are available. (A) the histogram of the average saliency distribution over microbial species. Species are sorted from left to right by decreasing saliency. Species prevalence in healthy (red) and affected (blue) individuals were calculated using a Wilcoxon test for each microbial species for two unpaired samples: healthy and affected individuals. (B) violin plots of the saliency distributions for microbial species grouped by prevalence: prevalent in affected (blue), prevalent in healthy (red), no prevalence (grey).

**S10 Table. Experimental results for the Random Forest with default Scikit-learn implementation.** Experiments are executed five times with random independent training-test splits. Values in brackets refer to the standard error over the repeated experiments. Reported values are test ROC AUC.

