## Supplementary material for "Microbiome-based disease prediction with multimodal variational information bottlenecks": S2 Table

**S2 Table. Complete experimental results for the multimodal microbiome-based disease prediction task with MVIB.**

| Dataset | Metrics | MVIB | |
|---|---|---|---|
| | | D | J |
| IBD | ROC AUC | 0.922 (0.02) | 0.932 (0.021) |
| | AC | 0.827 (0.022) | 0.827 (0.022) |
| | F1 | 0.8 (0.122) | 0.8 (0.122) |
| | P | 0.32 (0.049) | 0.32 (0.049) |
| | R | 0.457 (0.07) | 0.457 (0.07) |
| EW-T2D | ROC AUC | 0.859 (0.023) | 0.863 (0.02) |
| | AC | 0.76 (0.029) | 0.75 (0.022) |
| | F1 | 0.8 (0.029) | 0.797 (0.028) |
| | P | 0.764 (0.062) | 0.745 (0.053) |
| | R | 0.773 (0.034) | 0.763 (0.028) |
| C-T2D | ROC AUC | 0.75 (0.009) | 0.751 (0.013) |
| | AC | 0.67 (0.018) | 0.667 (0.018) |
| | F1 | 0.671 (0.009) | 0.662 (0.006) |
| | P | 0.647 (0.064) | 0.659 (0.067) |
| | R | 0.652 (0.035) | 0.654 (0.037) |
| Obesity | ROC AUC | 0.662 (0.024) | 0.667 (0.026) |
| | AC | 0.667 (0.019) | 0.659 (0.016) |
| | F1 | 0.682 (0.011) | 0.678 (0.009) |
| | P | 0.909 (0.017) | 0.903 (0.02) |
| | R | 0.779 (0.012) | 0.774 (0.011) |
| Cirrhosis | ROC AUC | 0.925 (0.005) | 0.925 (0.006) |
| | AC | 0.838 (0.014) | 0.838 (0.014) |
| | F1 | 0.893 (0.021) | 0.893 (0.021) |
| | P | 0.783 (0.048) | 0.783 (0.048) |
| | R | 0.829 (0.022) | 0.829 (0.022) |

(The table continues in the next page)

| Dataset | Metrics | MVIB | |
|---|---|---|---|
| | | D | J |
| Colorectal | ROC AUC | 0.78 (0.071) | 0.779 (0.073) |
| | AC | 0.728 (0.027) | 0.72 (0.022) |
| | F1 | 0.852 (0.078) | 0.848 (0.078) |
| | P | 0.42 (0.058) | 0.4 (0.045) |
| | R | 0.545 (0.051) | 0.529 (0.039) |
| Obesity-Joint | ROC AUC | 0.815 (0.019) | 0.818 (0.018) |
| | AC | 0.767 (0.022) | 0.758 (0.022) |
| | F1 | 0.768 (0.019) | 0.759 (0.018) |
| | P | 0.916 (0.012) | 0.916 (0.012) |
| | R | 0.835 (0.013) | 0.83 (0.014) |
| Colorectal-EMBL | ROC AUC | 0.811 (0.01) | 0.814 (0.013) |
| | AC | 0.745 (0.024) | 0.745 (0.017) |
| | F1 | 0.789 (0.042) | 0.813 (0.035) |
| | P | 0.642 (0.031) | 0.611 (0.027) |
| | R | 0.705 (0.027) | 0.694 (0.019) |
| Early-Colorectal-EMBL | ROC AUC | 0.535 (0.05) | 0.543 (0.048) |
| | AC | 0.55 (0.042) | 0.56 (0.04) |
| | F1 | 0.513 (0.063) | 0.527 (0.06) |
| | P | 0.378 (0.027) | 0.4 (0.027) |
| | R | 0.434 (0.041) | 0.453 (0.038) |
| Hypertension | ROC AUC | 0.603 (0.045) | 0.591 (0.041) |
| | AC | 0.8 (0.008) | 0.8 (0.008) |
| | F1 | 0.803 (0.004) | 0.803 (0.004) |
| | P | 0.994 (0.006) | 0.994 (0.006) |
| | R | 0.888 (0.004) | 0.888 (0.004) |

Results obtained optimising the $J_{MVIB-T}$ objective (see Equation **??**). Experiments are executed five times with random independent training-test splits. Values in brackets refer to the standard error over the repeated experiments. All values in the table refer to metrics computed on the test sets. ROC AUC: area under the receiver operating characteristic curve. AC: classification accuracy. F1: F1 score. P: precision. R: recall. D and J refer to the two pre-processing techniques adopted and the two collections of datasets obtained: *default* (D) and *joint* (J).
