## Supplementary material for "Microbiome-based disease prediction with multimodal variational information bottlenecks": S4 Table

**S4 Table. Comparison of different objective functions and pre-processing techniques.**

| Dataset | $J_{MVIB-T}$ | | $J_{MVIB}$ | |
|---|---|---|---|---|
| | D | J | D | J |
| IBD | 0.922 (0.020) | **0.936 (0.014)** | 0.915 (0.018) | 0.915 (0.018) |
| EW-T2D | **0.859 (0.023)** | 0.853 (0.025) | 0.859 (0.024) | 0.855 (0.027) |
| C-T2D | 0.750 (0.009) | **0.758 (0.012)** | 0.754 (0.014) | 0.756 (0.015) |
| Obesity | 0.662 (0.024) | 0.666 (0.027) | **0.673 (0.028)** | 0.672 (0.030) |
| Cirrhosis | 0.925 (0.005) | 0.924 (0.005) | **0.930 (0.002)** | 0.928 (0.002) |
| Colorectal | 0.780 (0.071) | 0.777 (0.069) | 0.788 (0.059) | **0.796 (0.055)** |
| Obesity-Joint | 0.815 (0.019) | 0.818 (0.018) | 0.825 (0.019) | **0.825 (0.018)** |
| Colorectal-EMBL | 0.811 (0.010) | 0.814 (0.013) | 0.827 (0.012) | **0.830 (0.011)** |
| Early-Colorectal-EMBL | 0.535 (0.050) | **0.543 (0.048)** | 0.525 (0.051) | 0.533 (0.053) |
| Hypertension | 0.602 (0.045) | 0.608 (0.043) | 0.622 (0.048) | **0.624 (0.049)** |
