## Supplementary figures and images for "Microbiome-based disease prediction with multimodal variational information bottlenecks"

### 0_embeddings.png

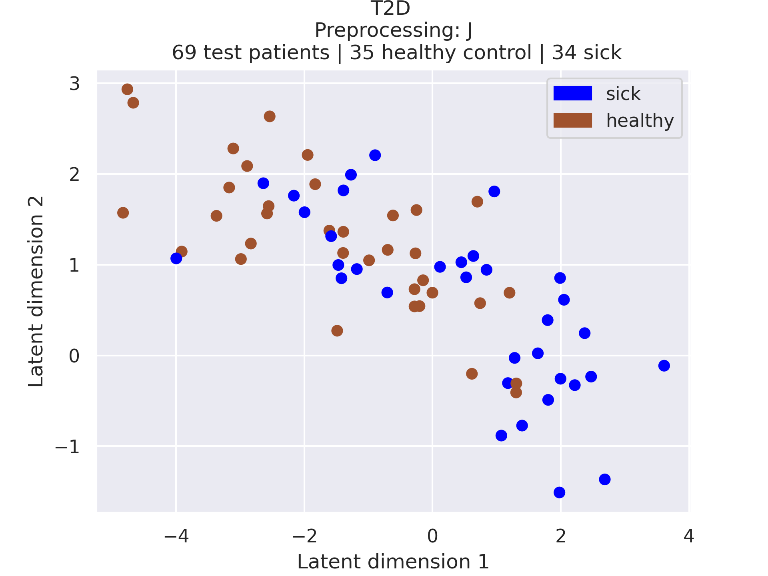

### 0_embeddings.png

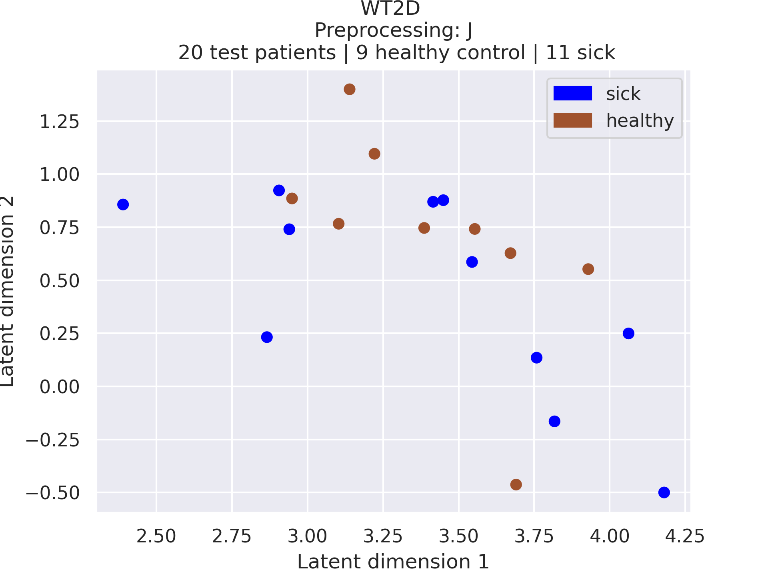

### 0_embeddings.png

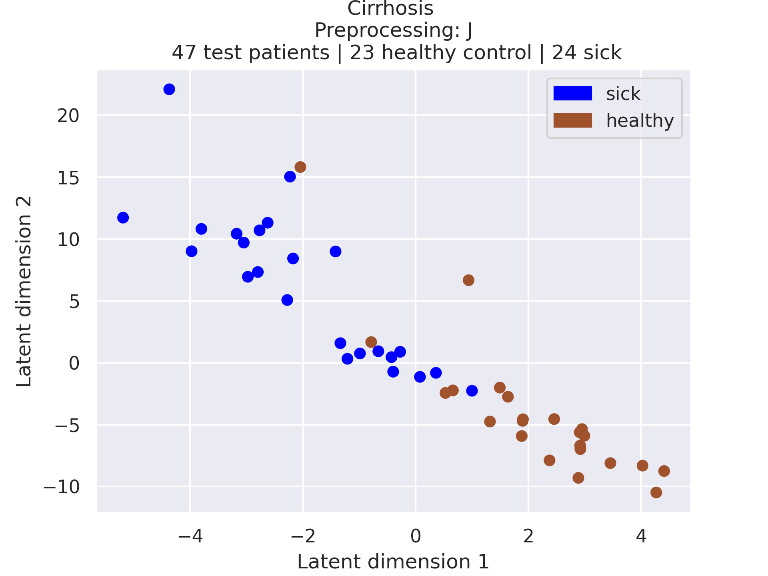

### 0_embeddings.png

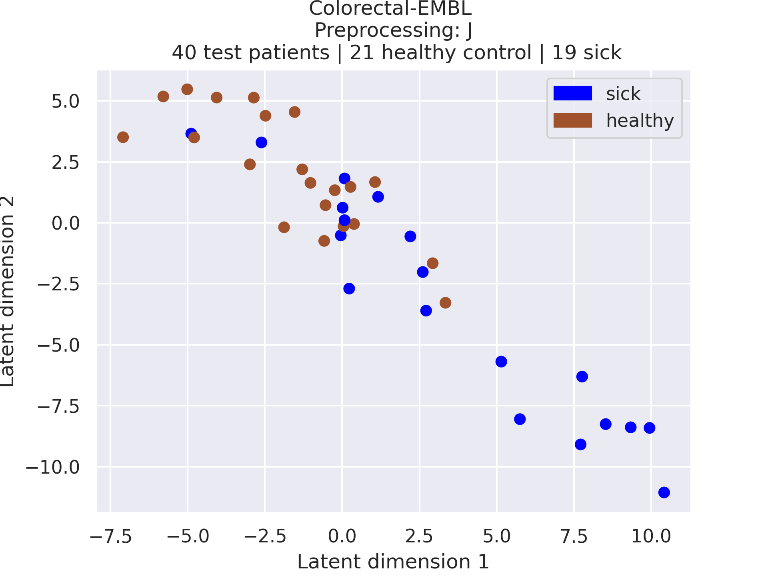

### 0_embeddings.png

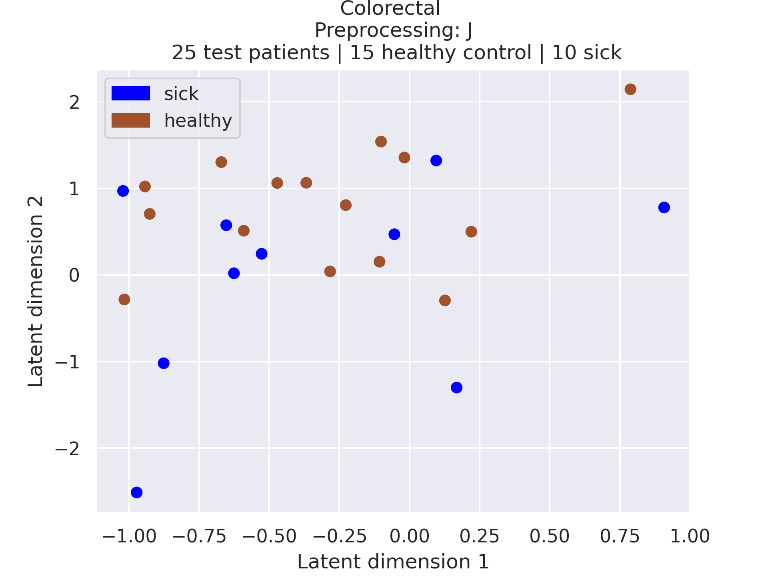

### 0_embeddings.png

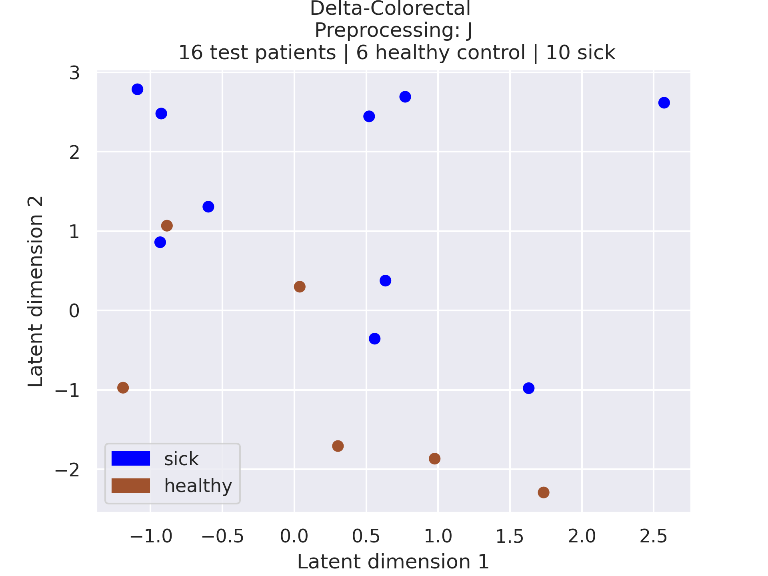

### 0_embeddings.png

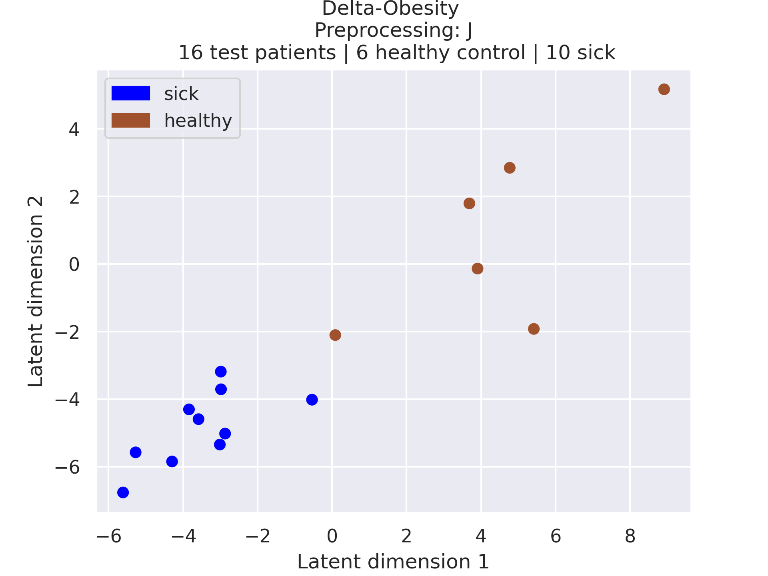

### 0_embeddings.png

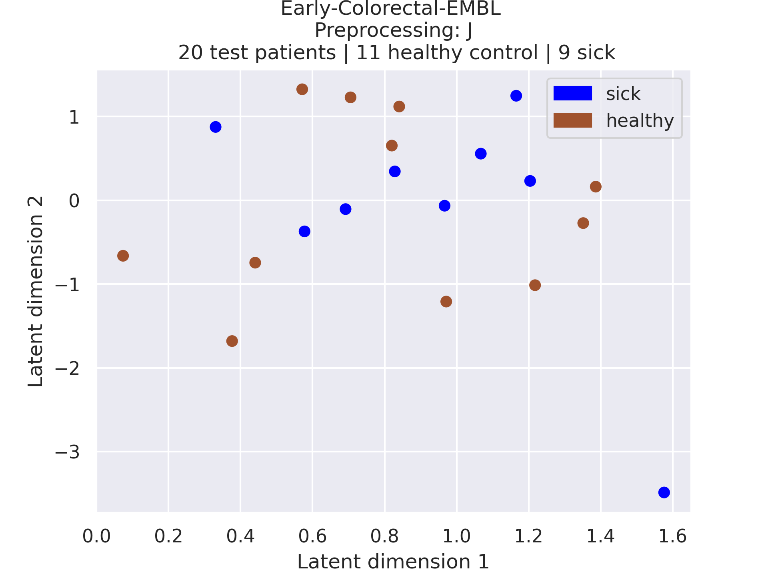

### 0_embeddings.png

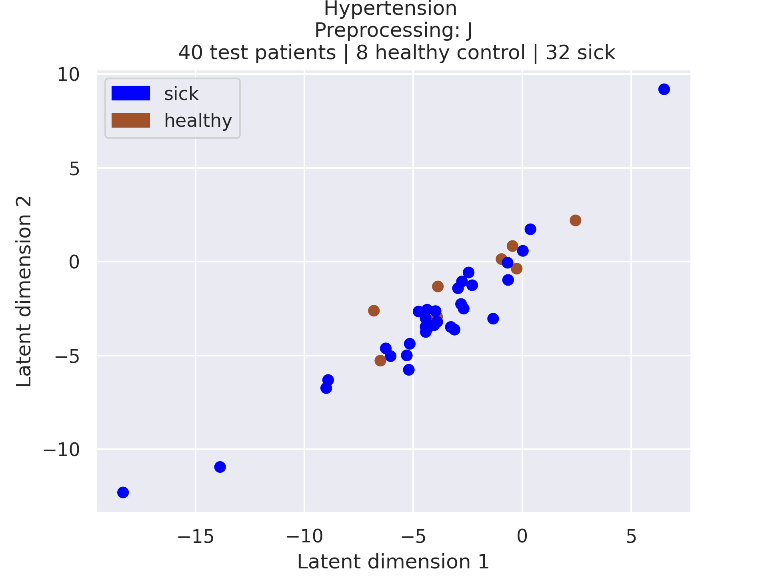

### 0_embeddings.png

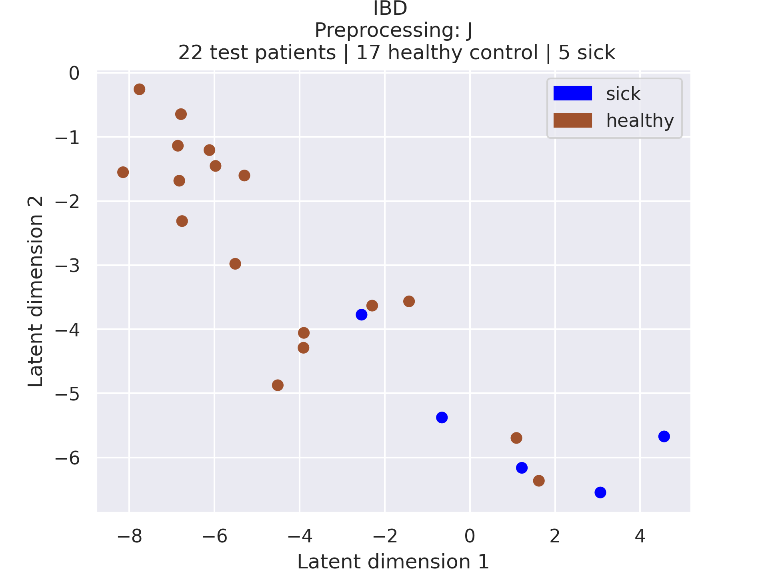

### 0_embeddings_95_confidence.png

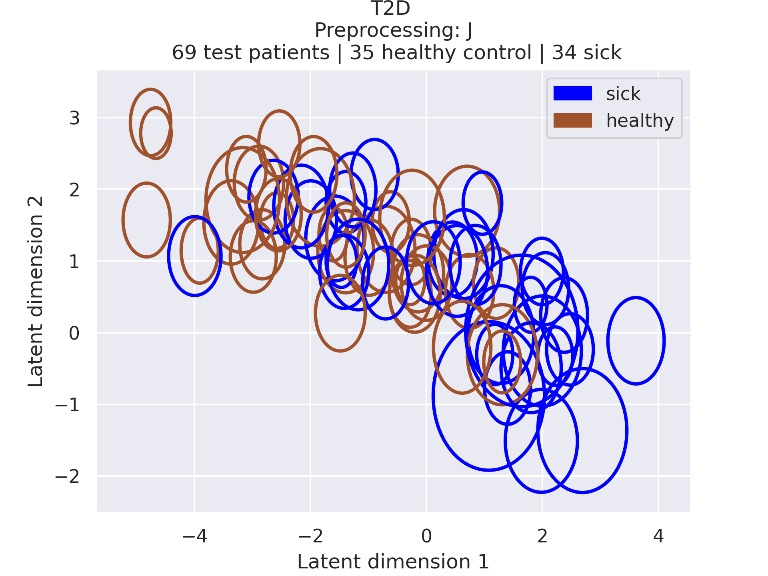

### 0_embeddings_95_confidence.png

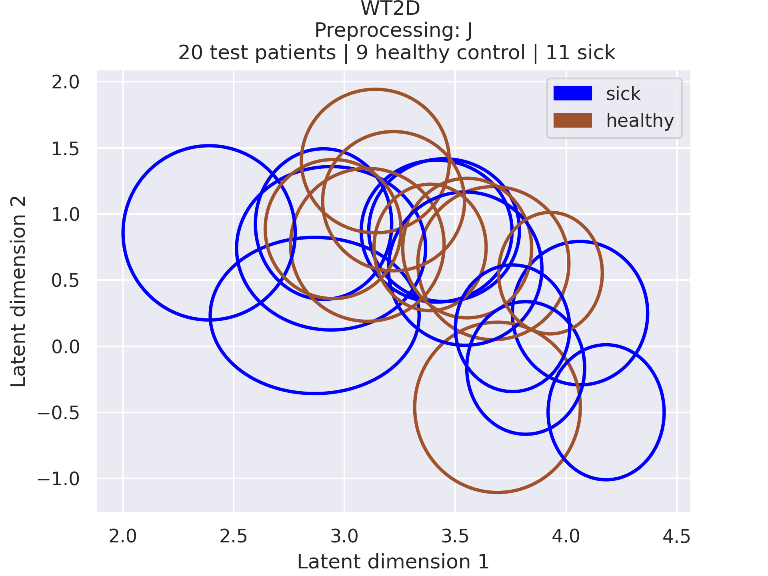

### 0_embeddings_95_confidence.png

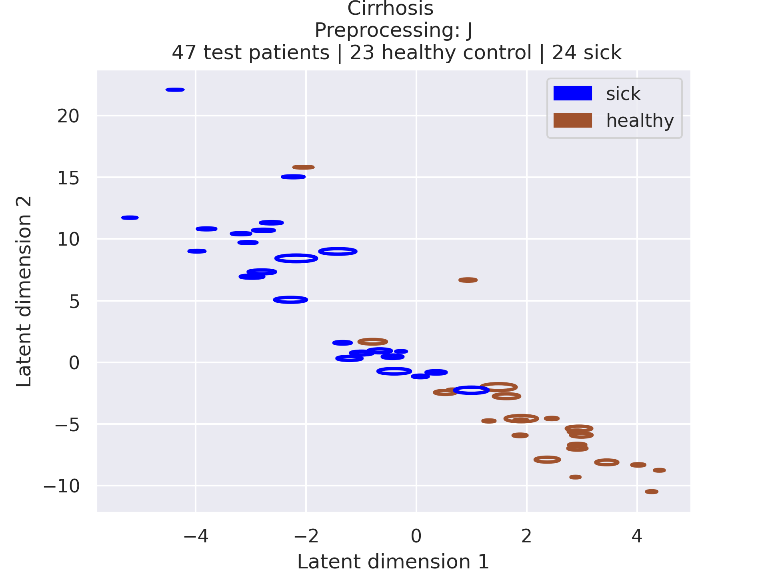

### 0_embeddings_95_confidence.png

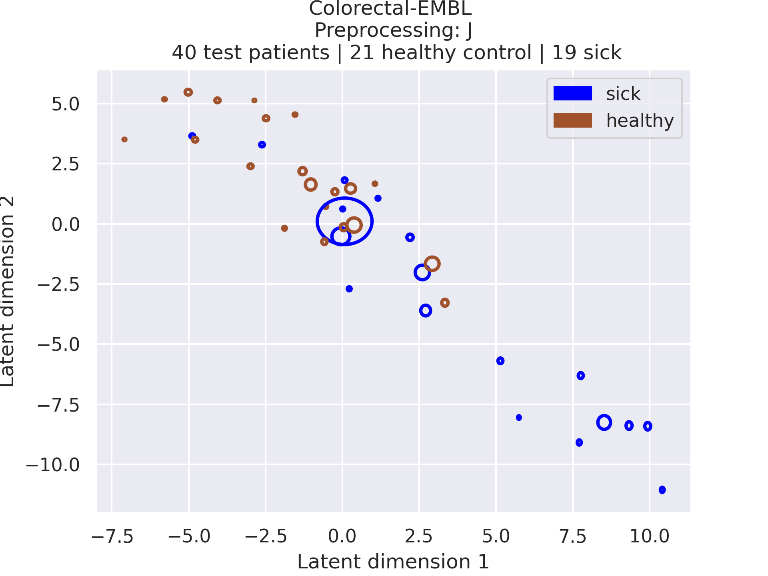

### 0_embeddings_95_confidence.png

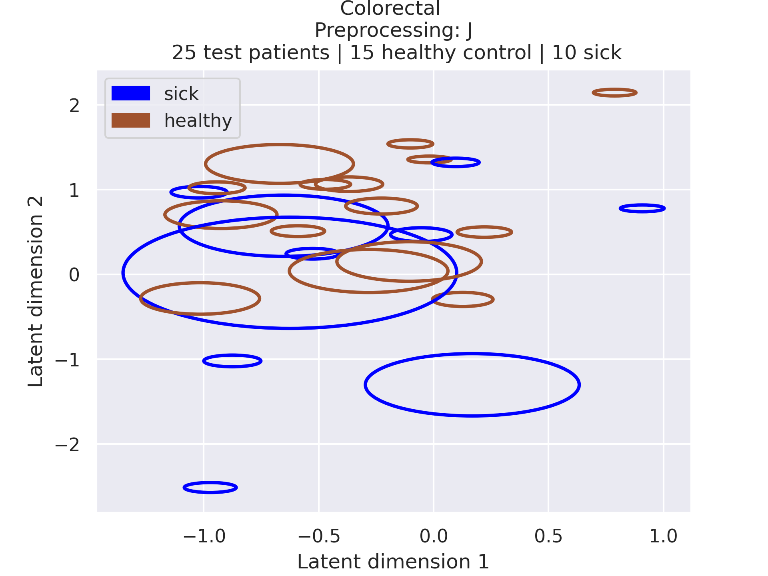

### 0_embeddings_95_confidence.png

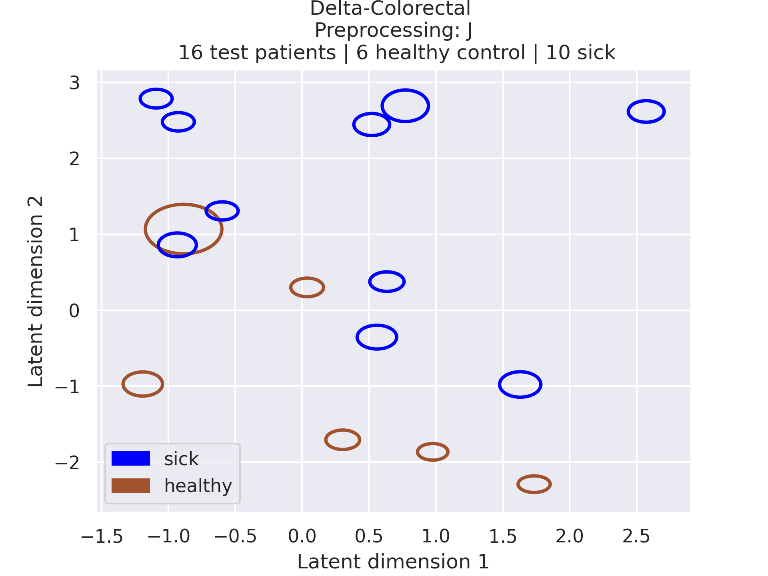

### 0_embeddings_95_confidence.png

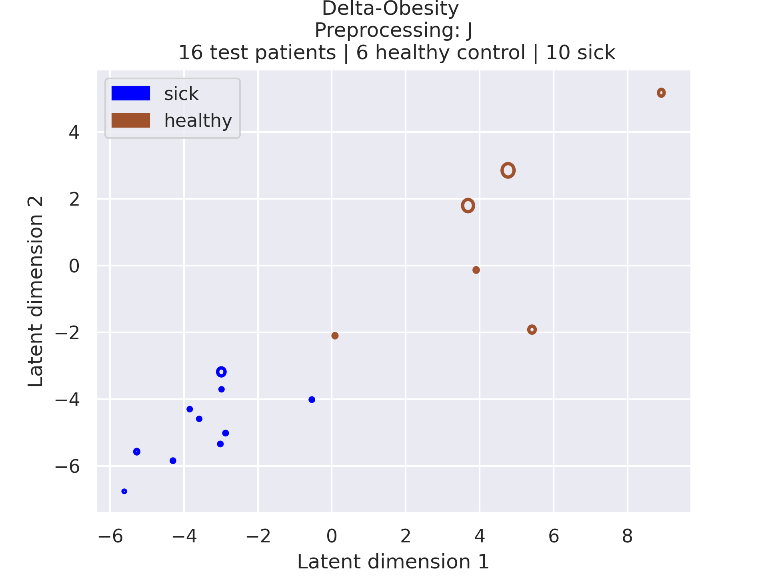

### 0_embeddings_95_confidence.png

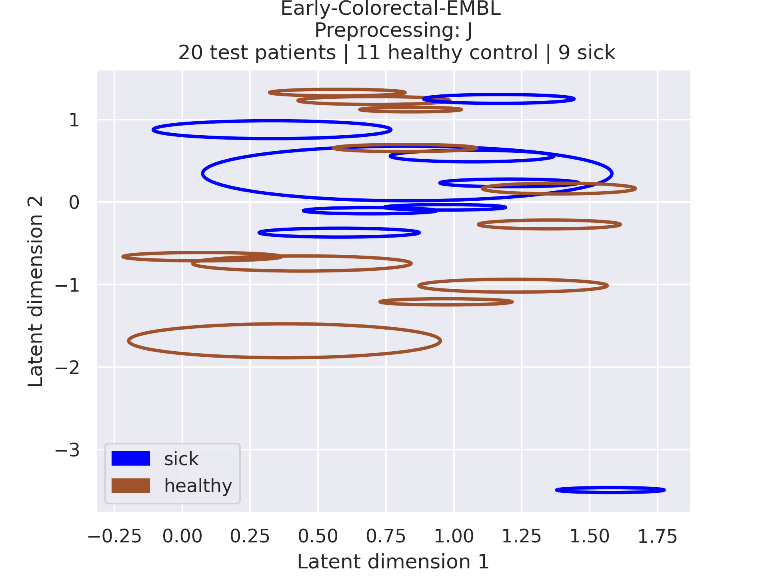

### 0_embeddings_95_confidence.png

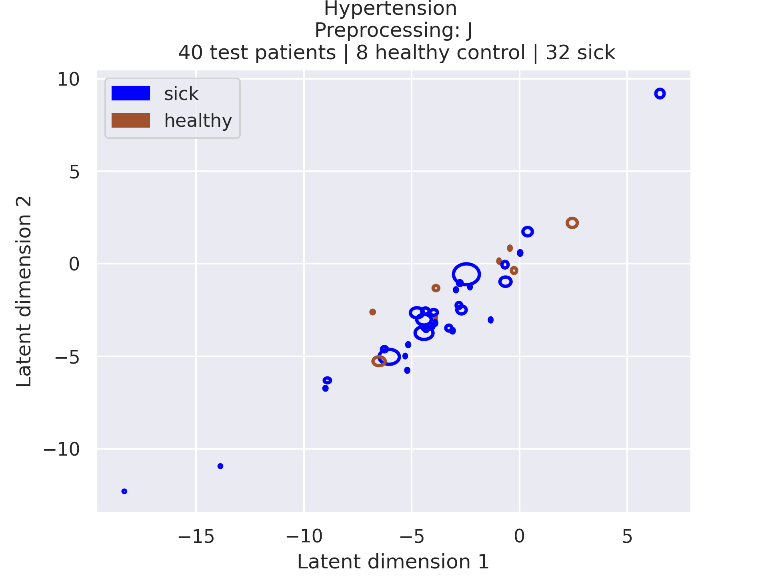

### 0_embeddings_95_confidence.png

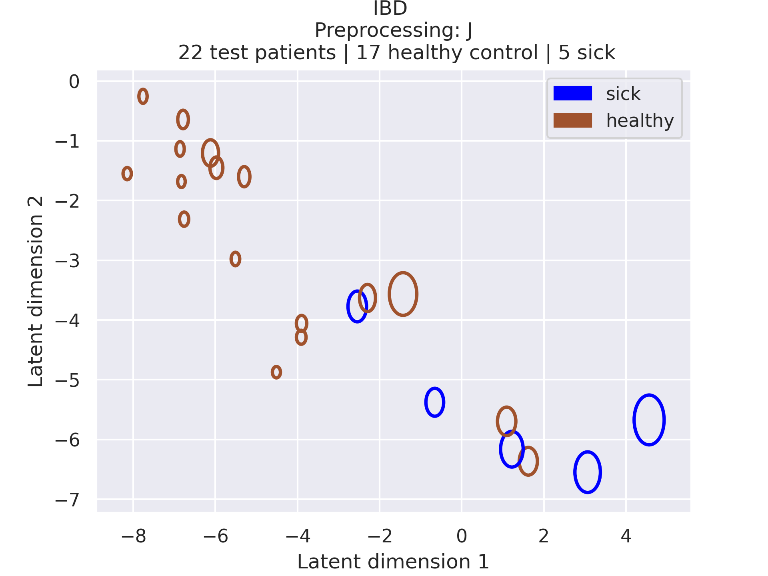

### 1_embeddings.png

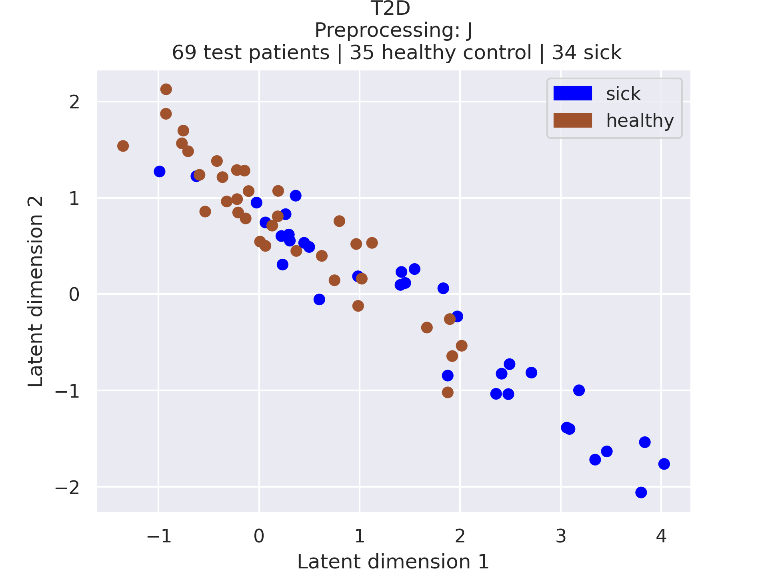

### 1_embeddings.png

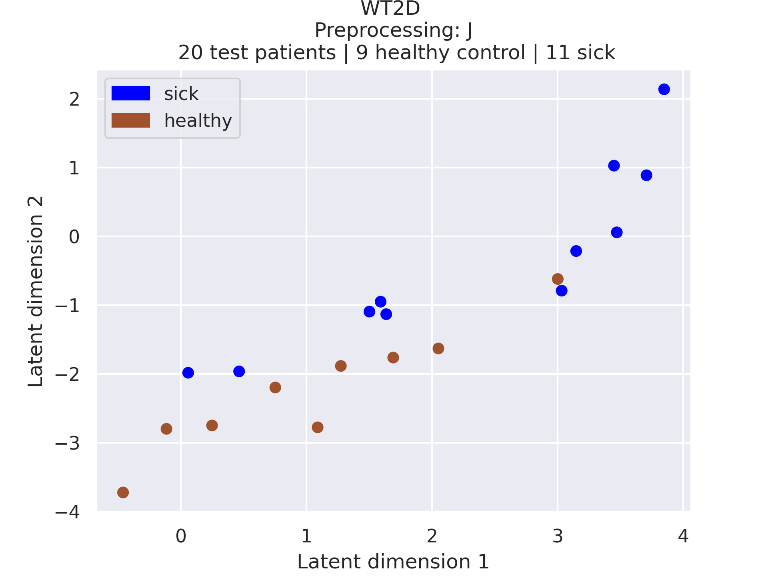

### 1_embeddings.png

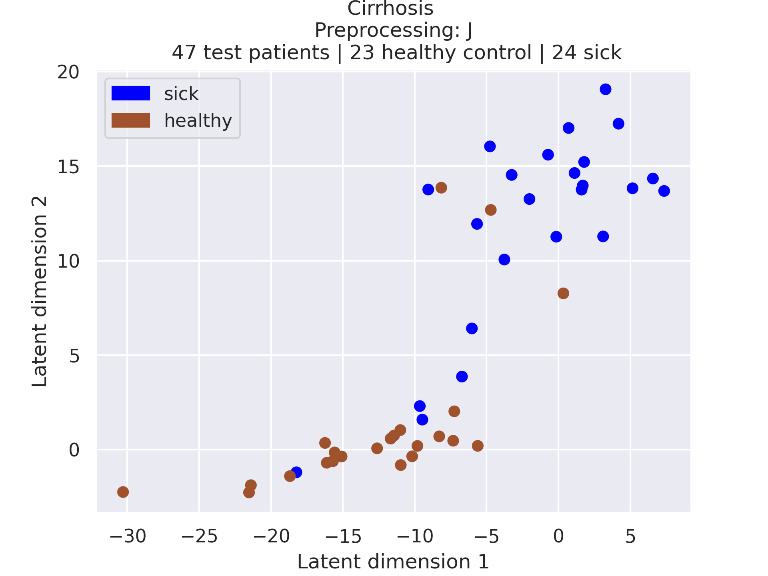

### 1_embeddings.png

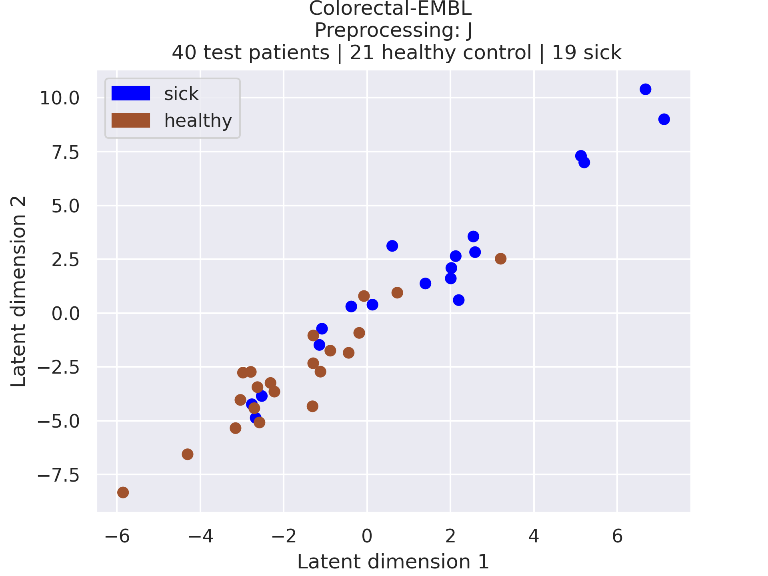

### 1_embeddings.png

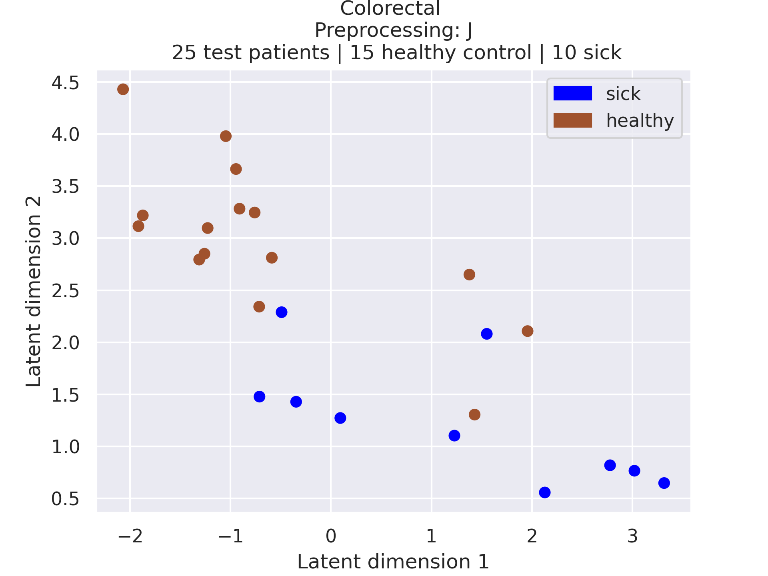

### 1_embeddings.png

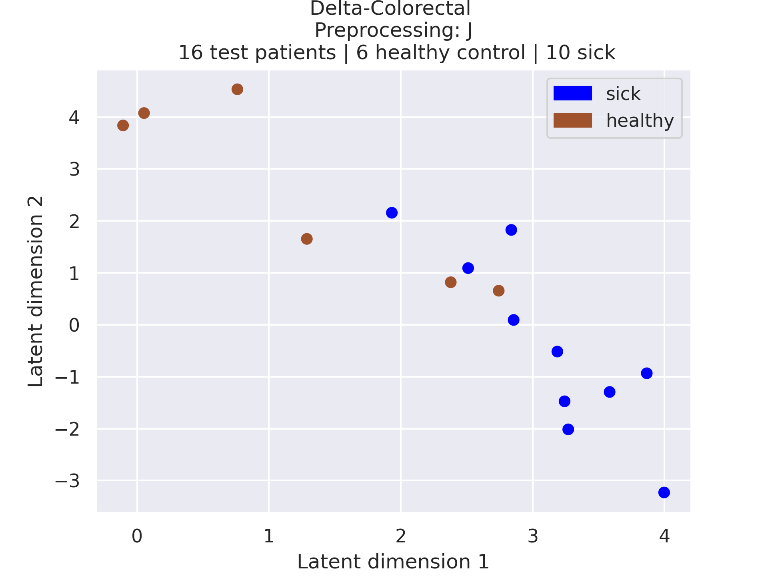

### 1_embeddings.png

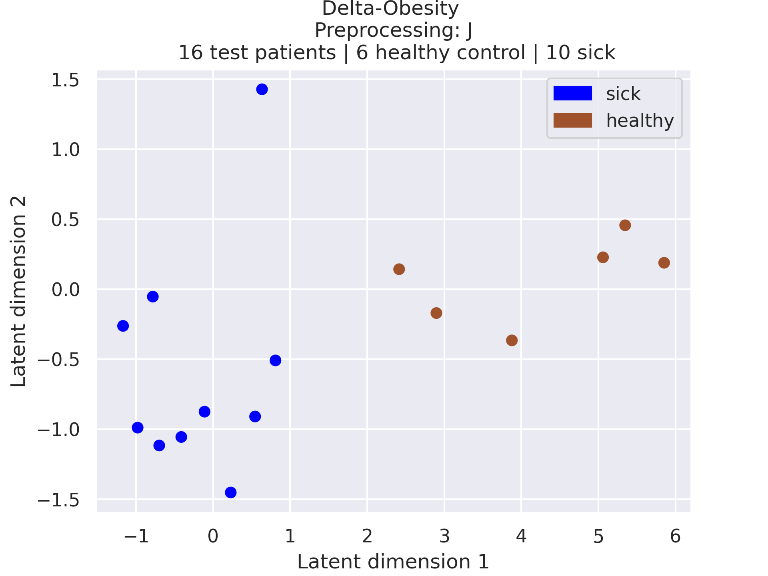

### 1_embeddings.png

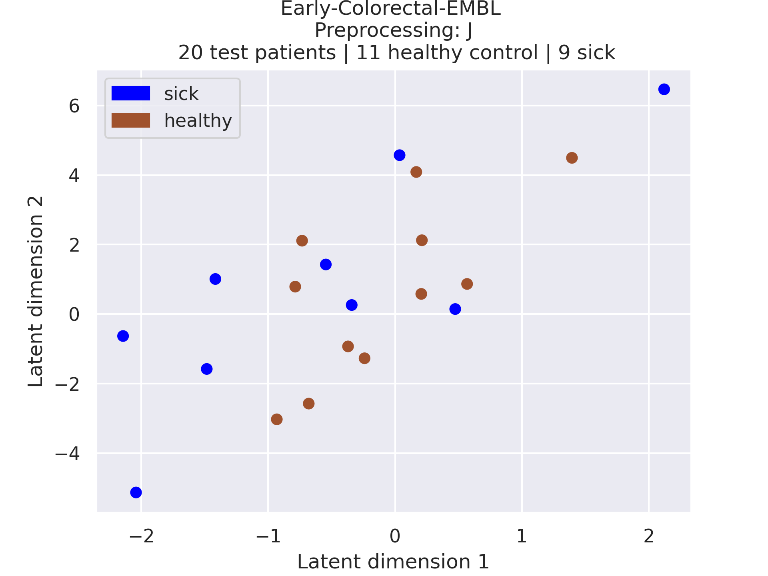

### 1_embeddings.png

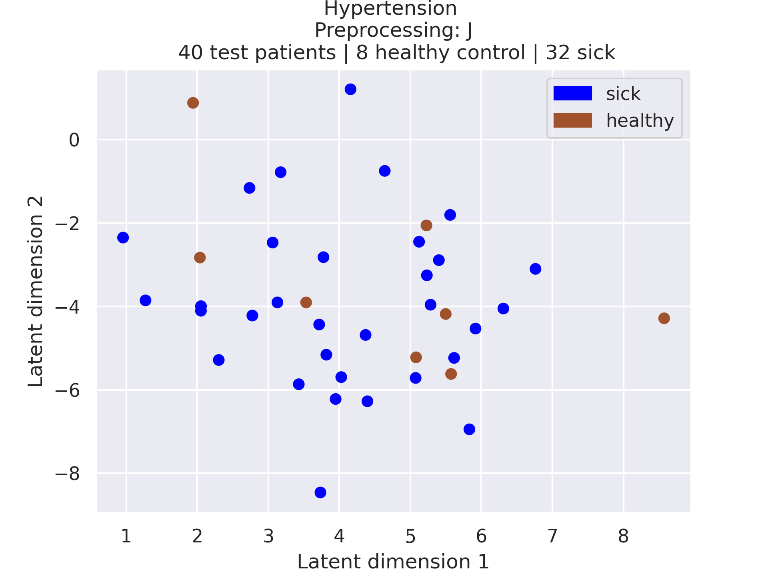

### 1_embeddings.png

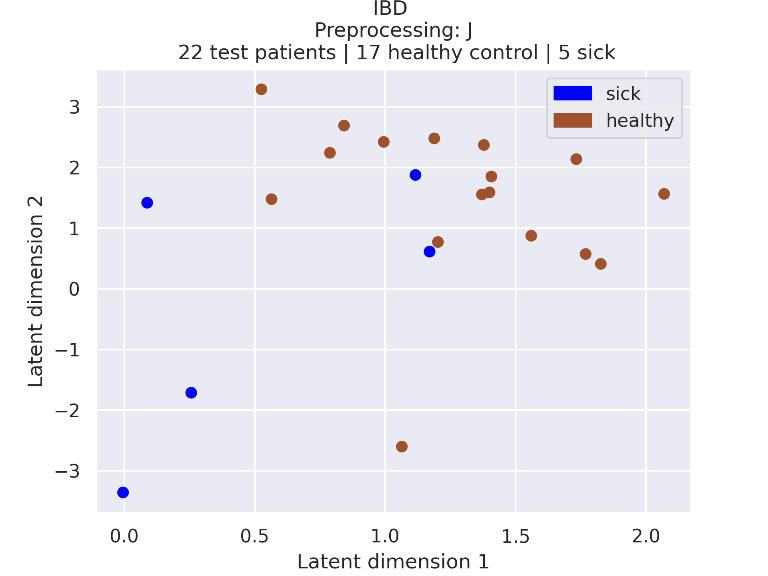

### 1_embeddings_95_confidence.png

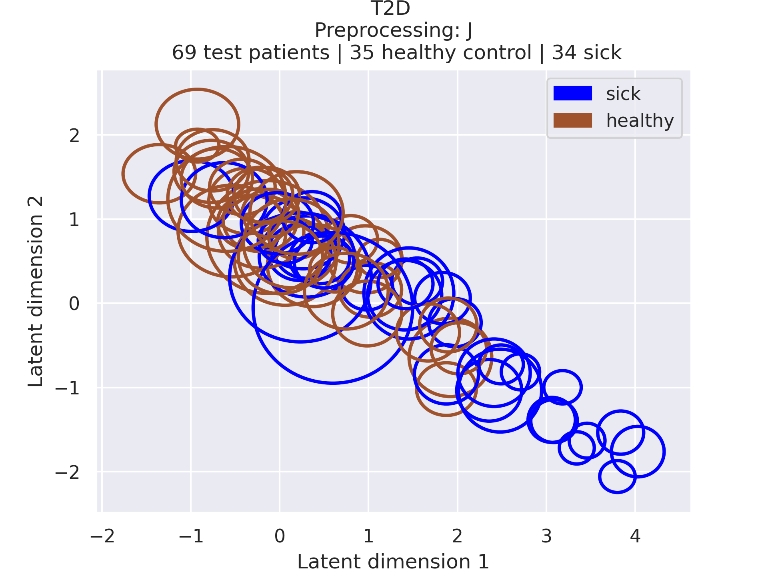

### 1_embeddings_95_confidence.png

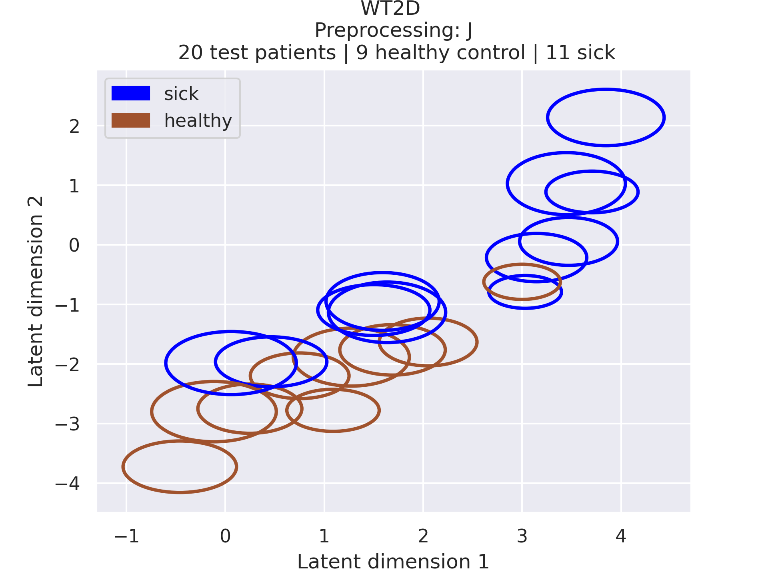

### 1_embeddings_95_confidence.png

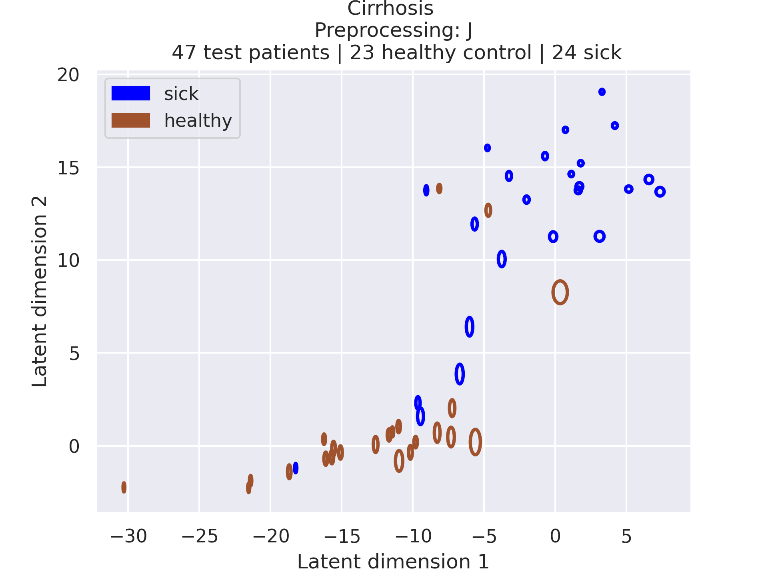

### 1_embeddings_95_confidence.png

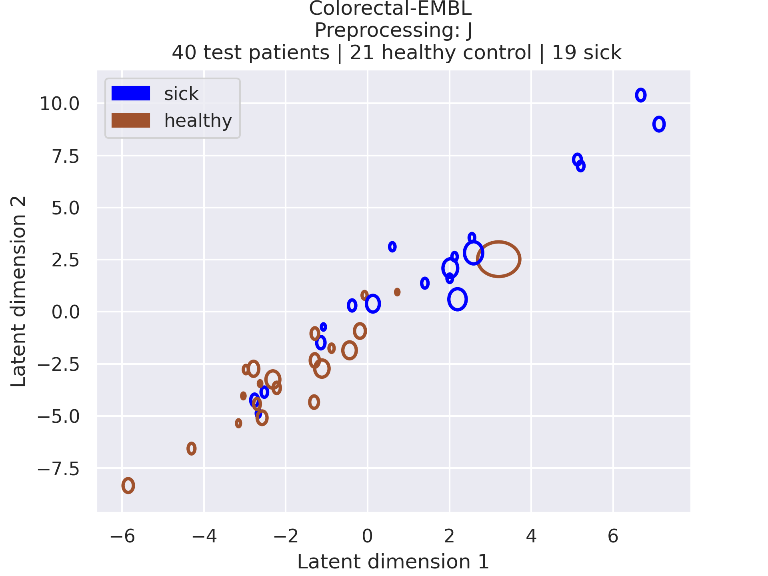

### 1_embeddings_95_confidence.png

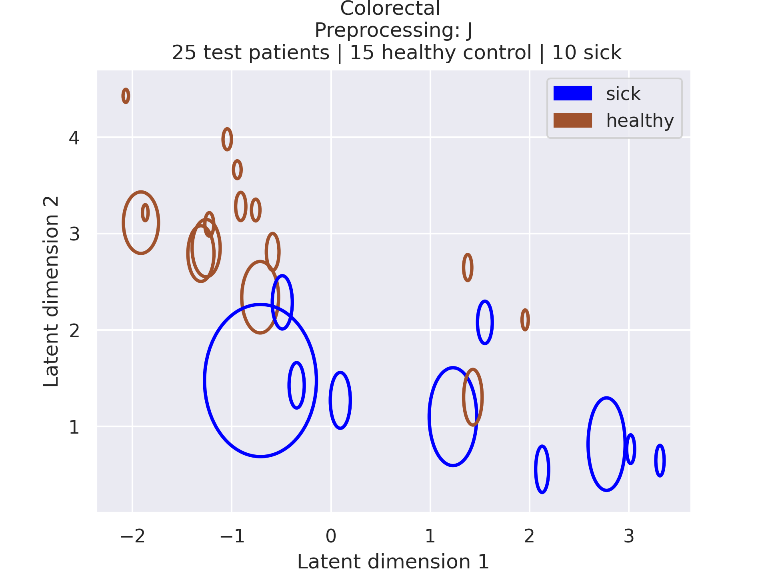

### 1_embeddings_95_confidence.png

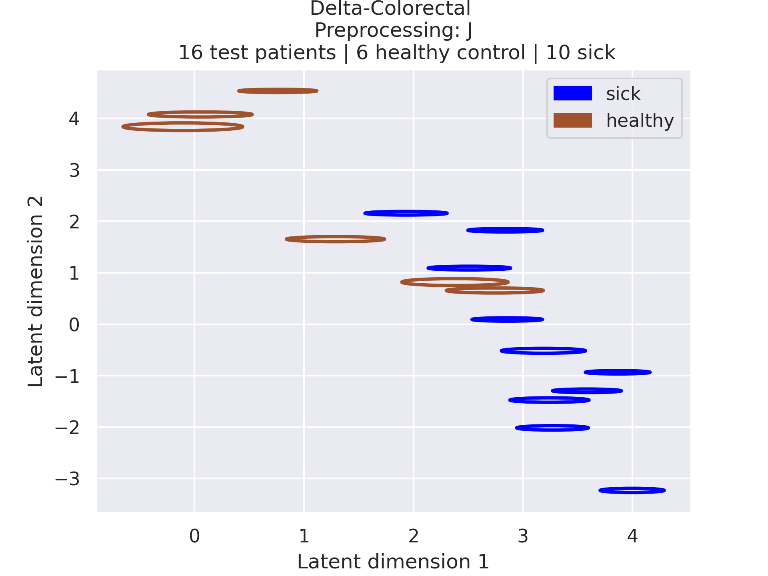

### 1_embeddings_95_confidence.png

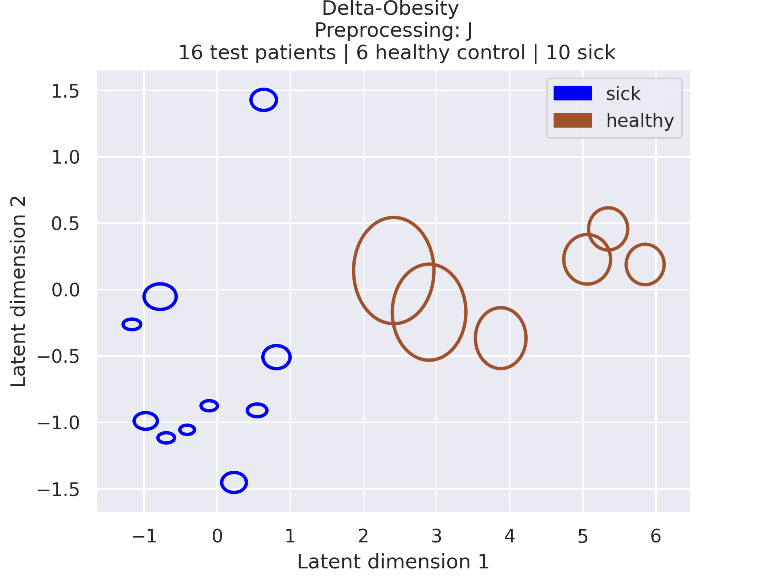

### 1_embeddings_95_confidence.png

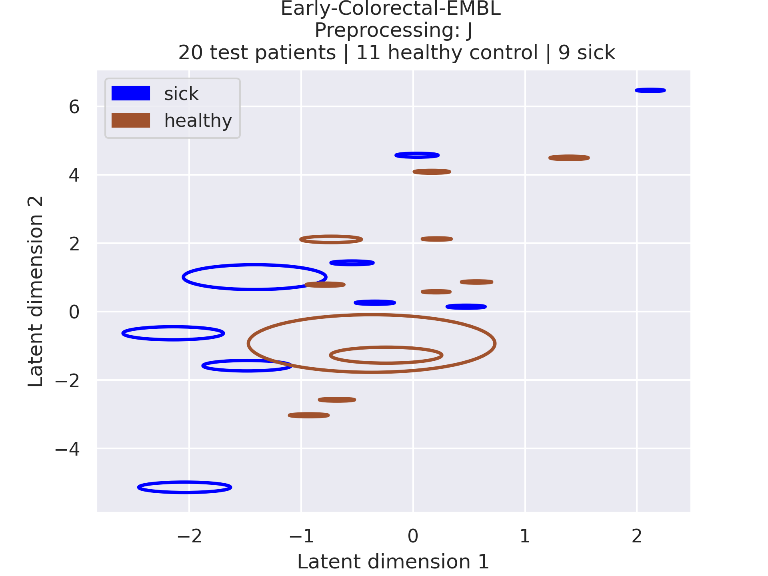

### 1_embeddings_95_confidence.png

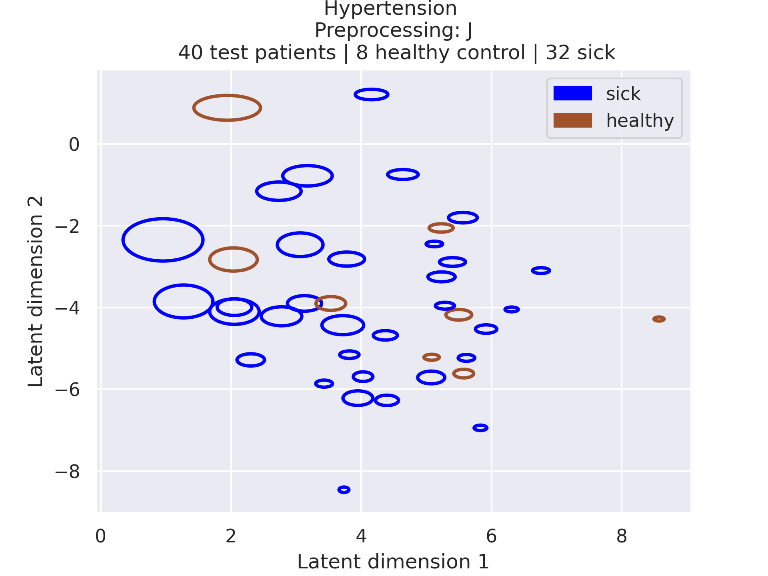

### 1_embeddings_95_confidence.png

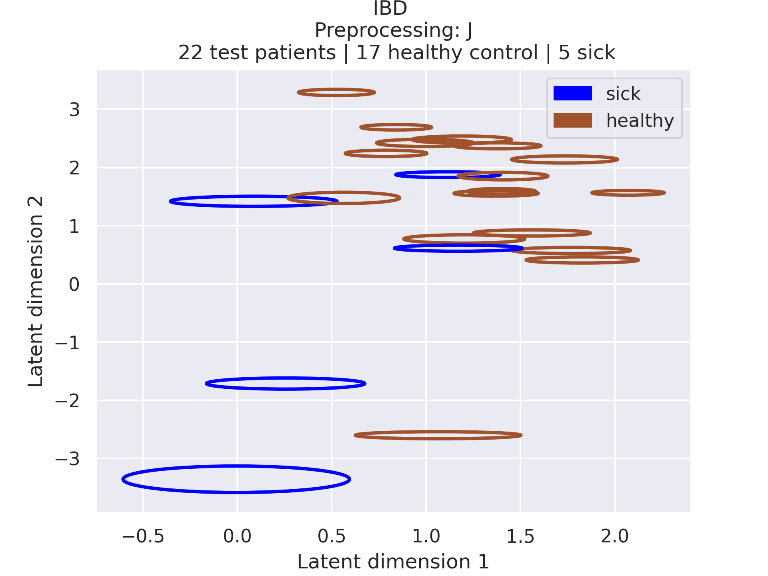

### 2_embeddings.png

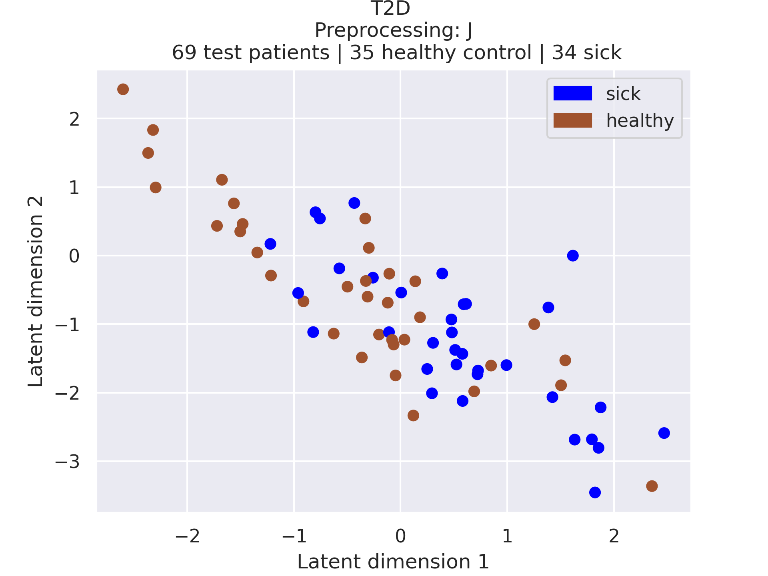

### 2_embeddings.png

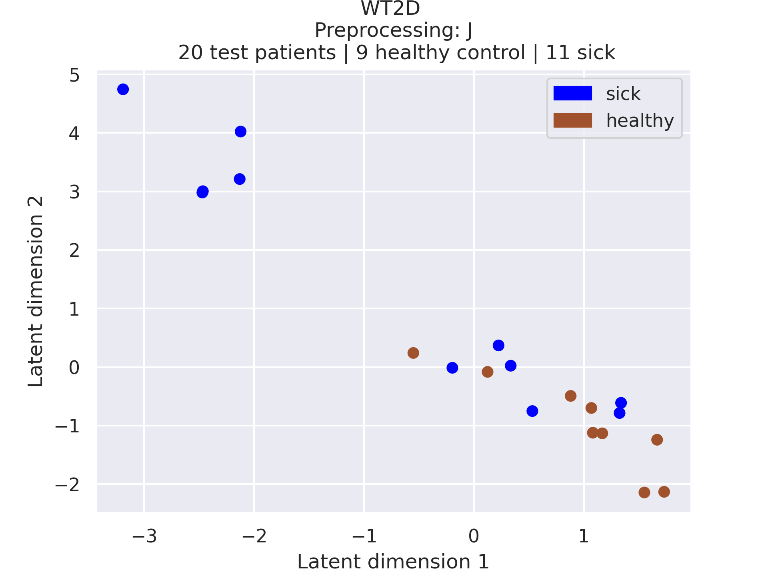

### 2_embeddings.png

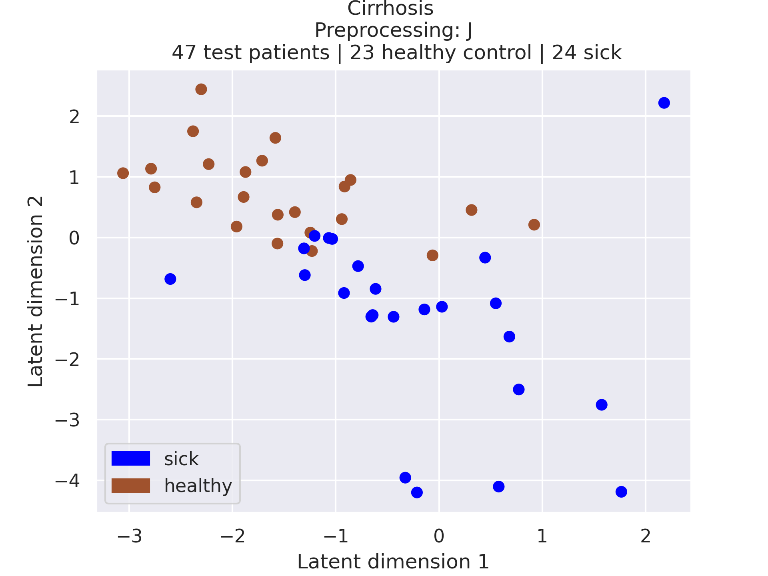

### 2_embeddings.png

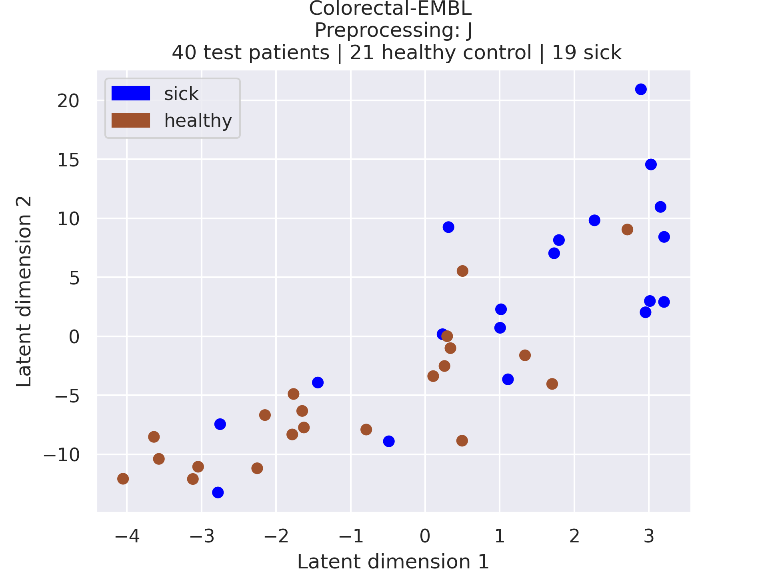

### 2_embeddings.png

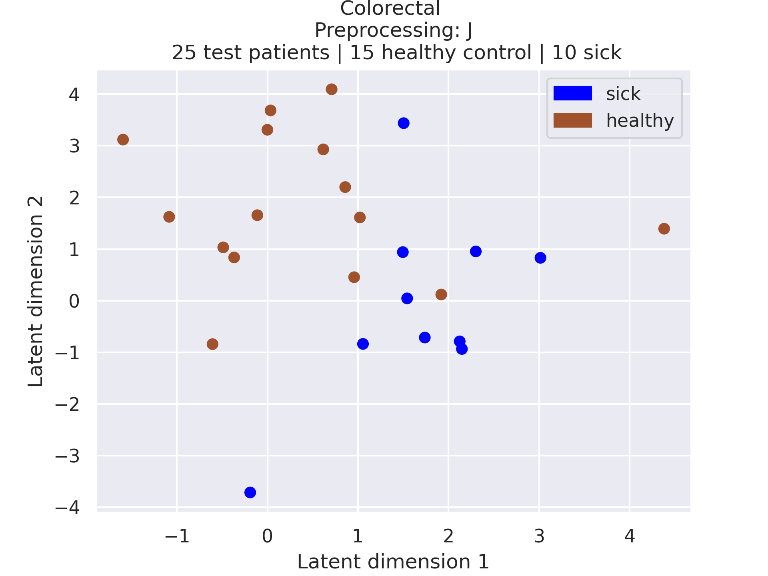

### 2_embeddings.png

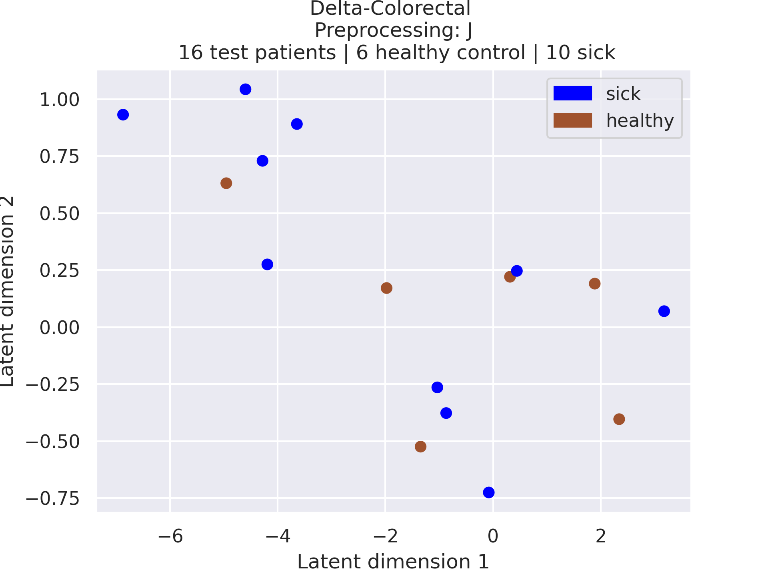

### 2_embeddings.png

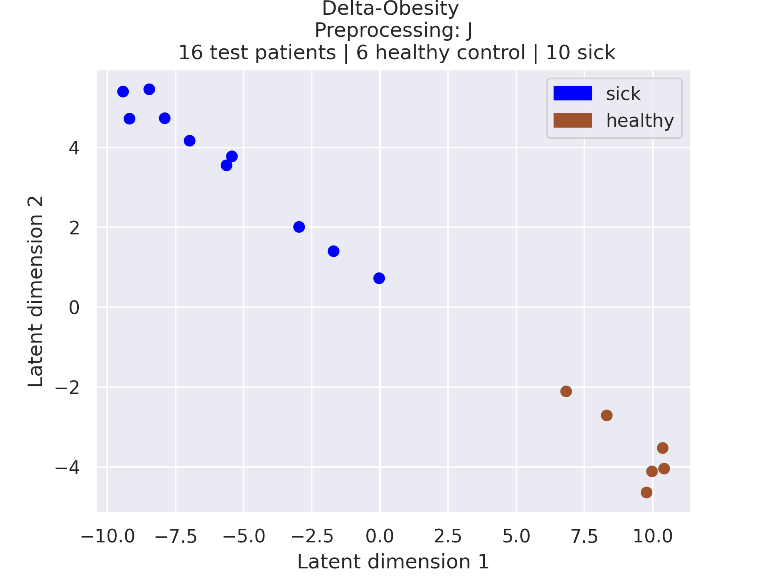

### 2_embeddings.png

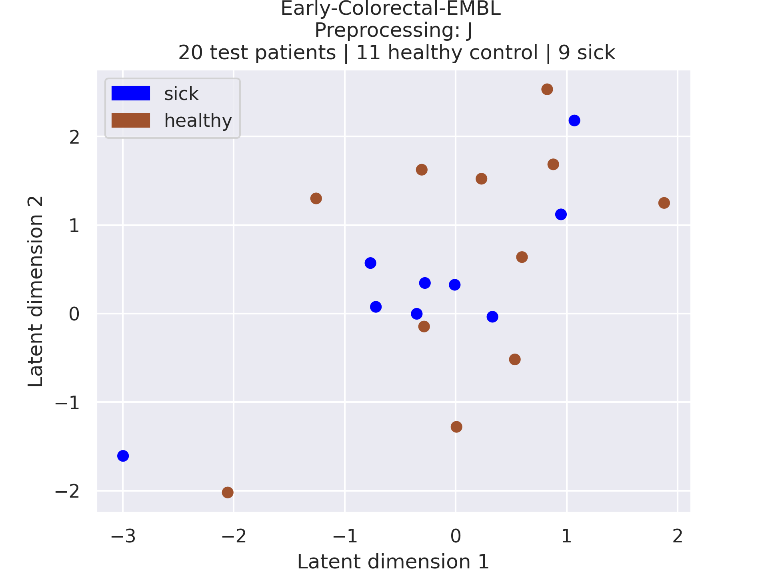

### 2_embeddings.png

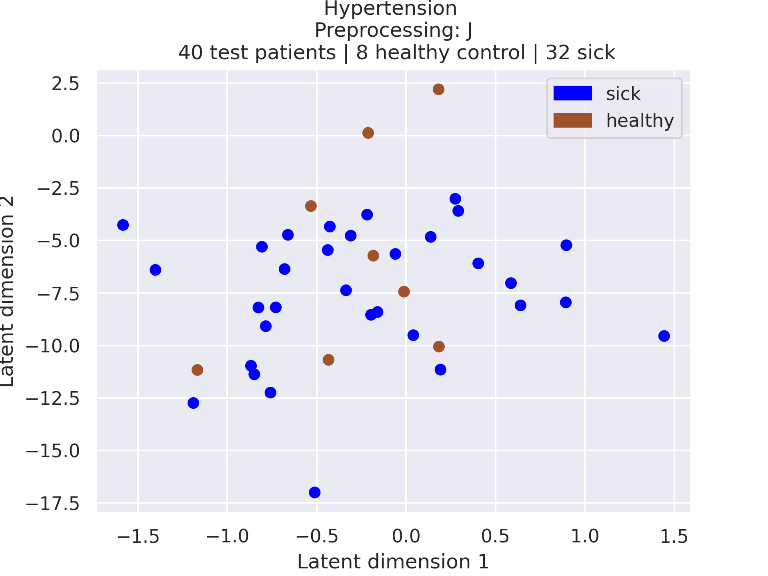

### 2_embeddings.png

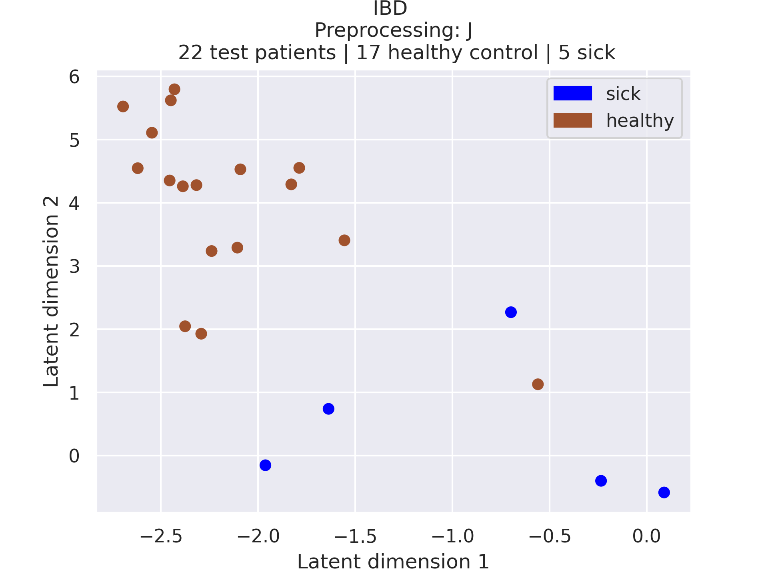

### 2_embeddings_95_confidence.png

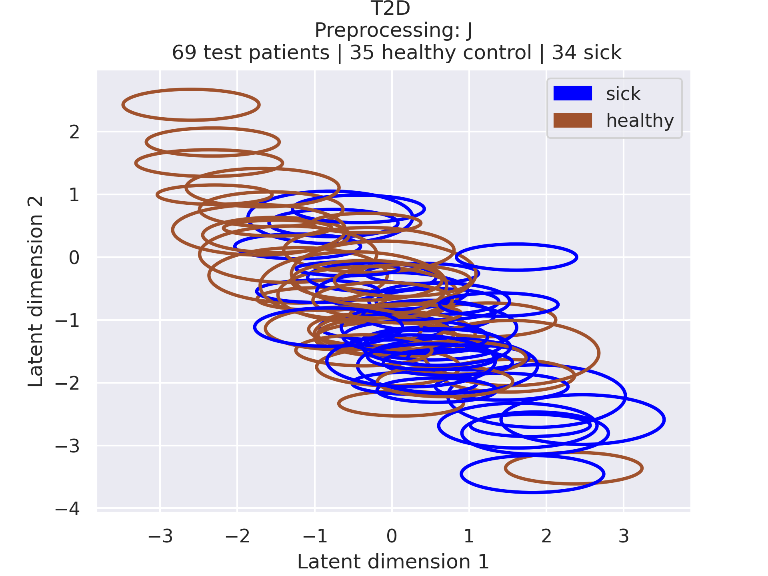

### 2_embeddings_95_confidence.png

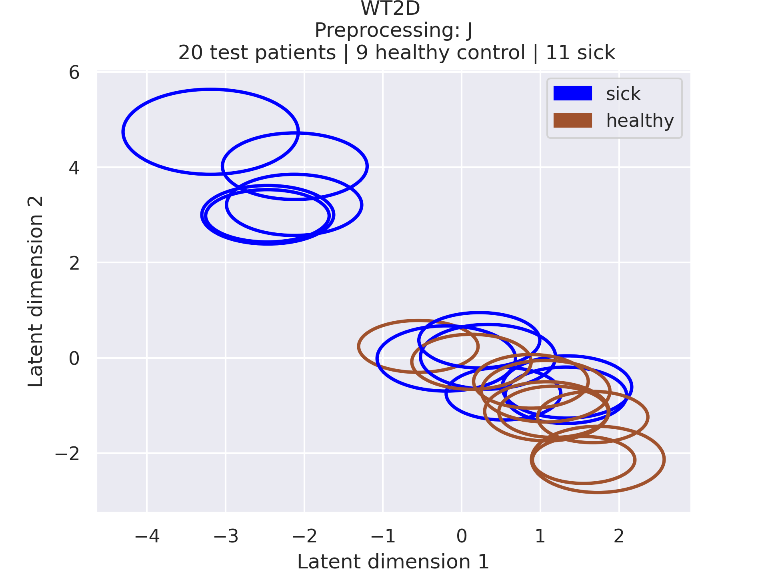

### 2_embeddings_95_confidence.png

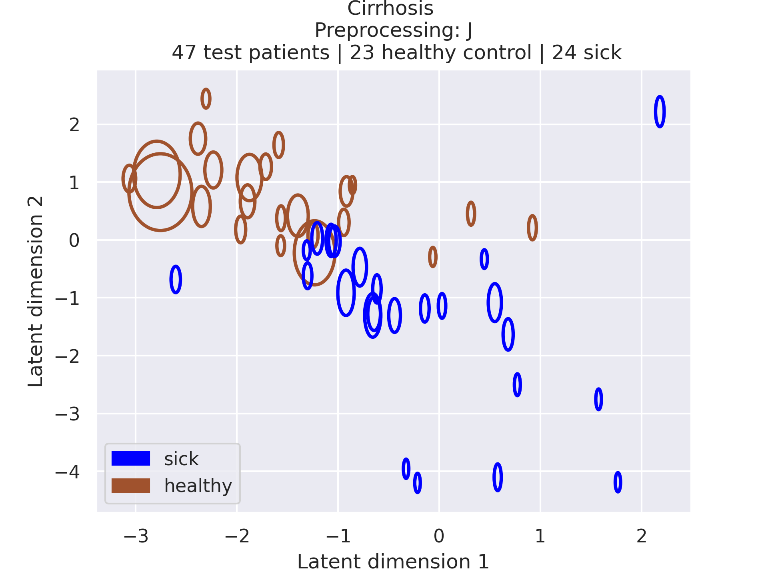

### 2_embeddings_95_confidence.png

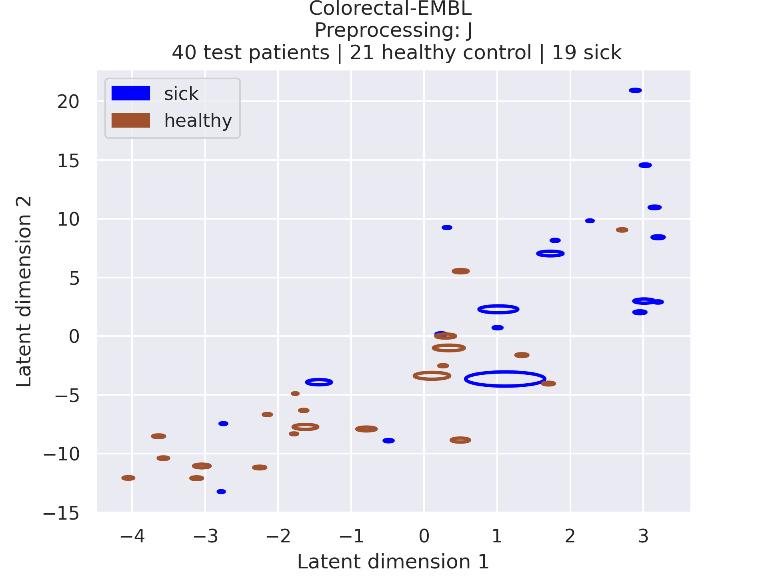

### 2_embeddings_95_confidence.png

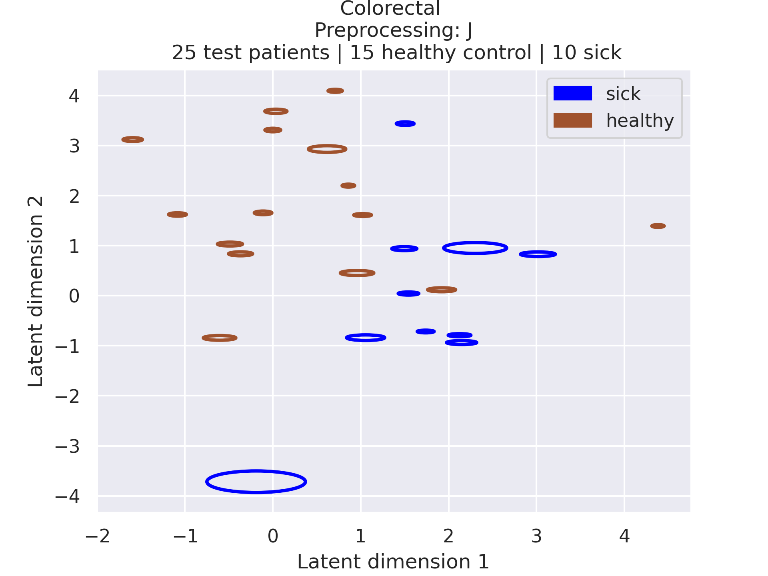

### 2_embeddings_95_confidence.png

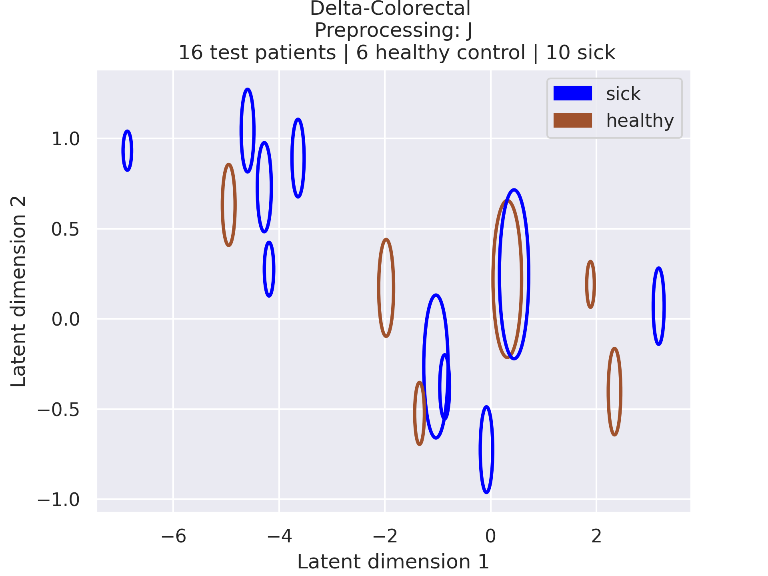

### 2_embeddings_95_confidence.png

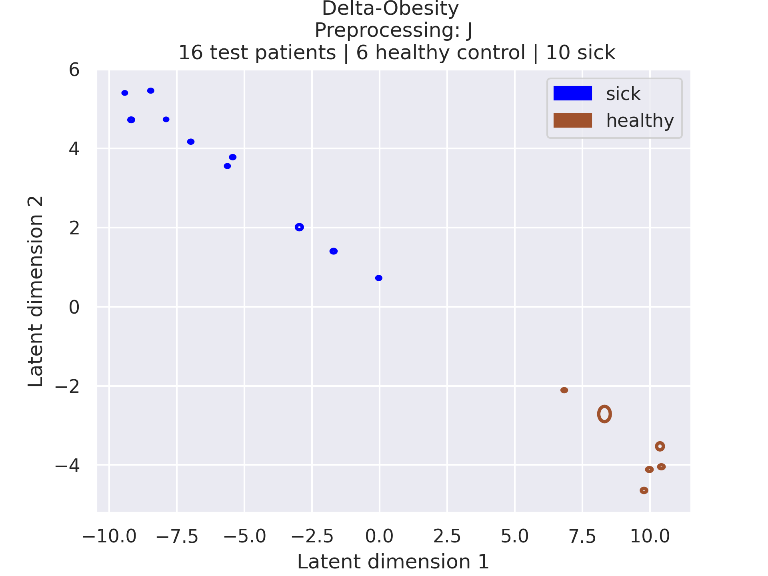

### 2_embeddings_95_confidence.png

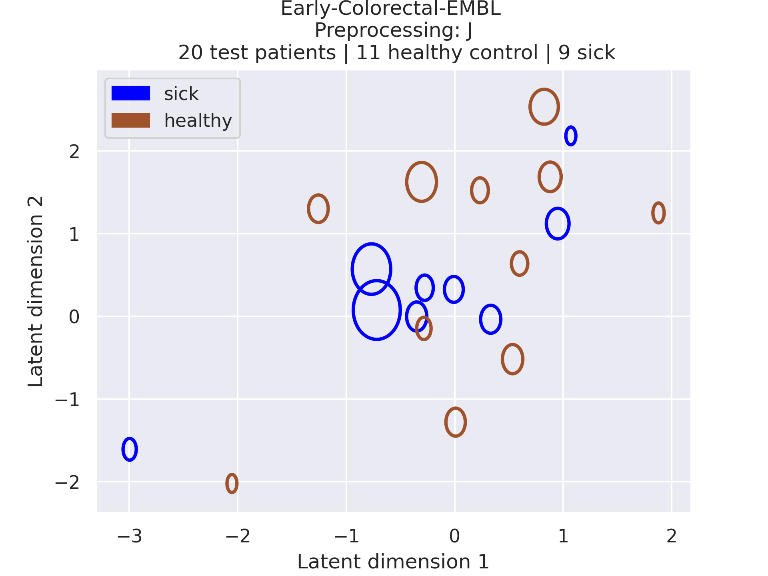

### 2_embeddings_95_confidence.png

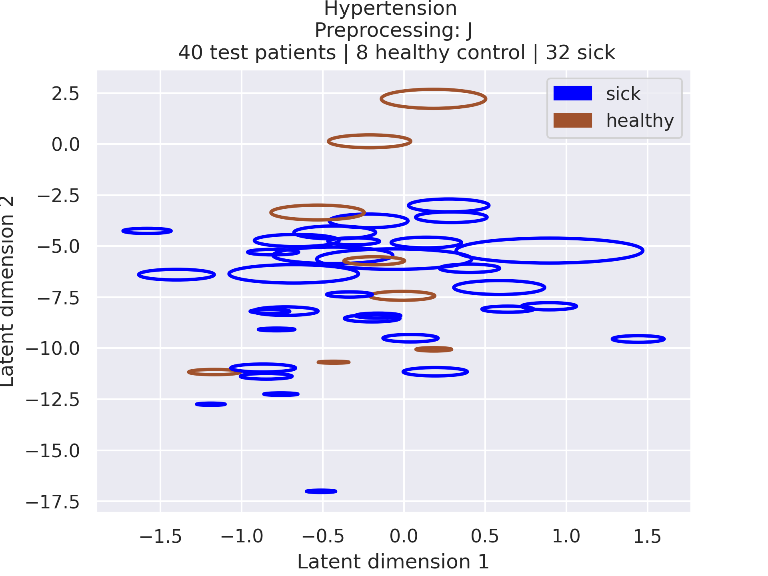

### 2_embeddings_95_confidence.png

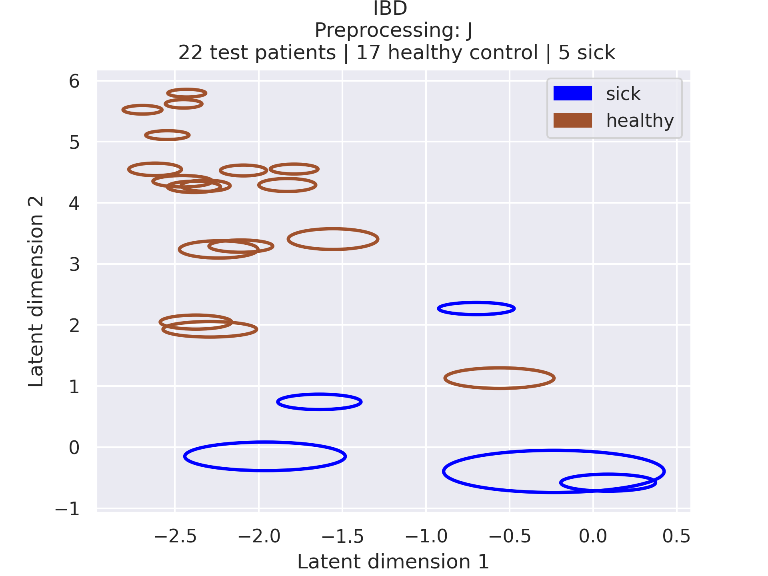

### 3_embeddings.png

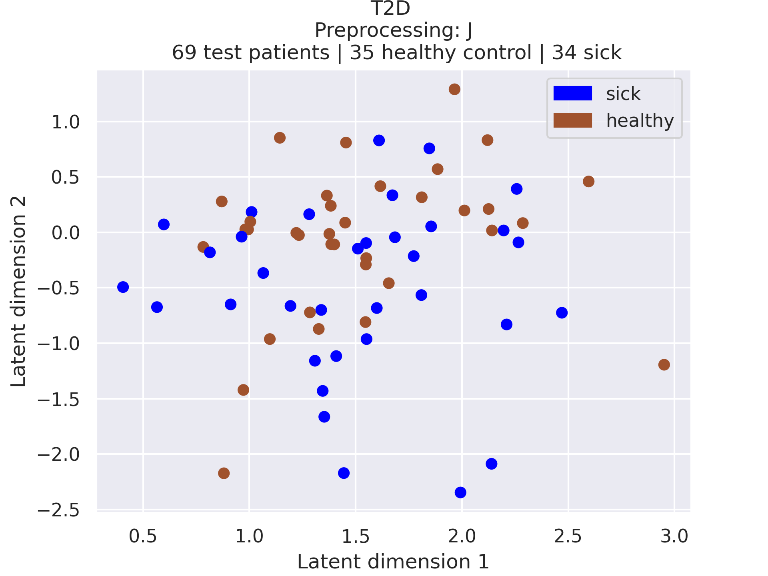

### 3_embeddings.png

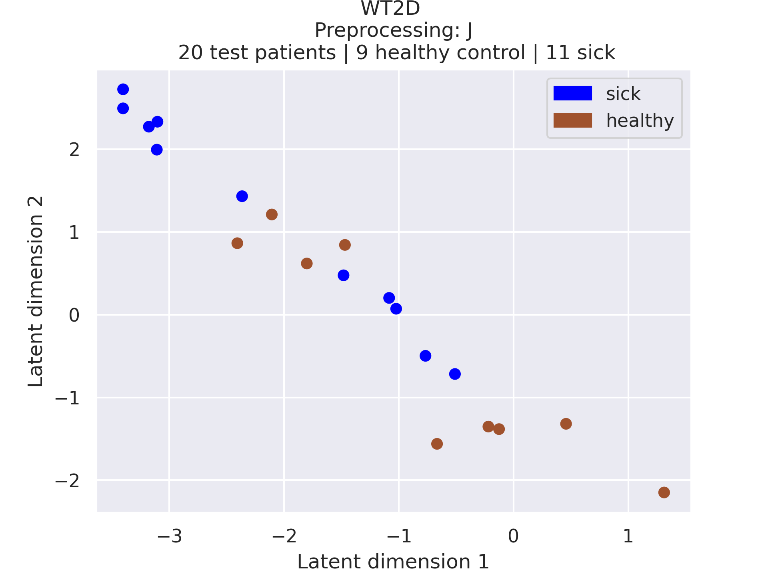

### 3_embeddings.png

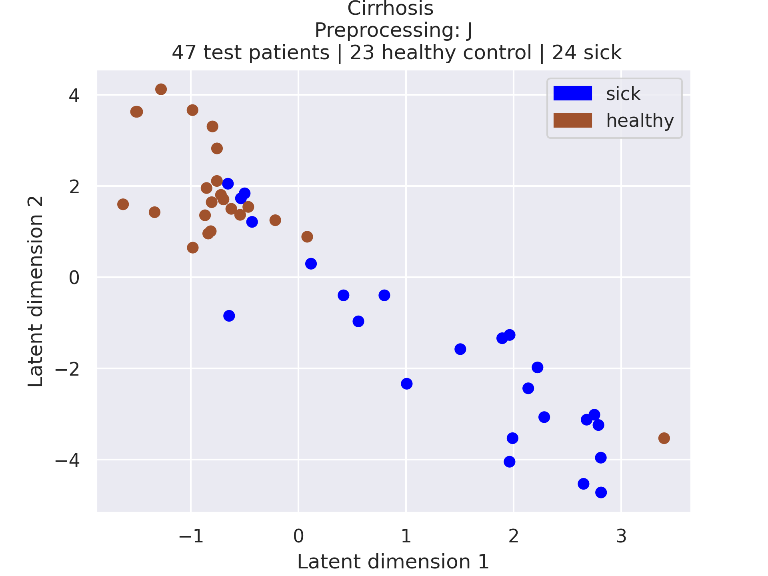

### 3_embeddings.png

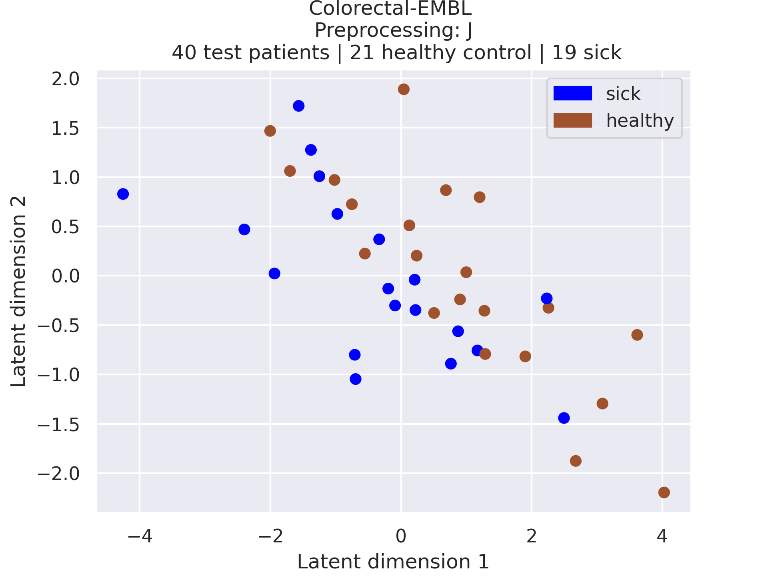

### 3_embeddings.png

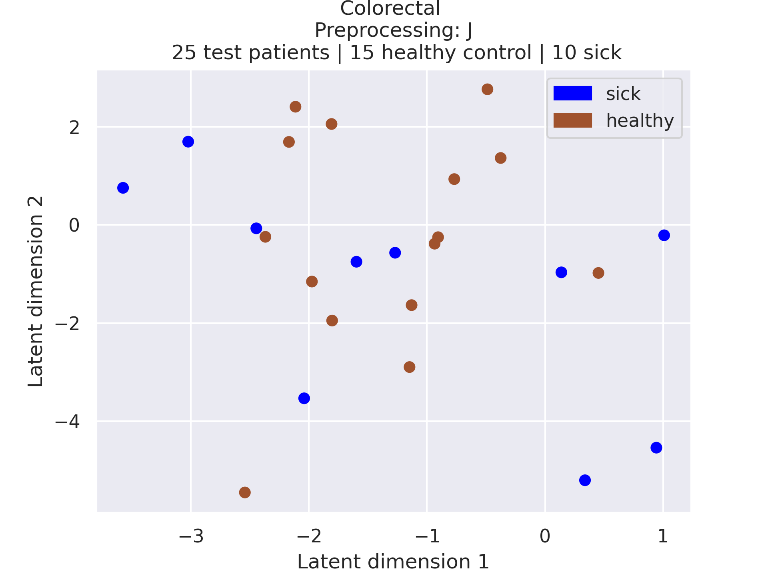

### 3_embeddings.png

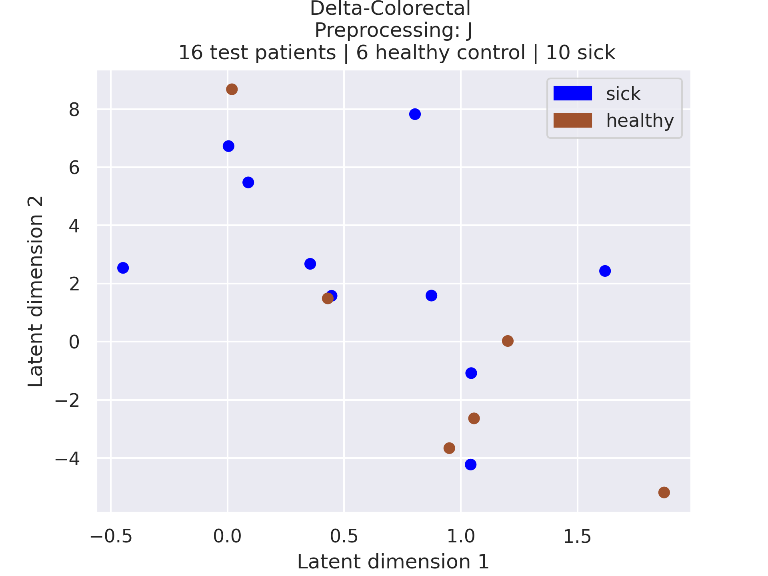

### 3_embeddings.png

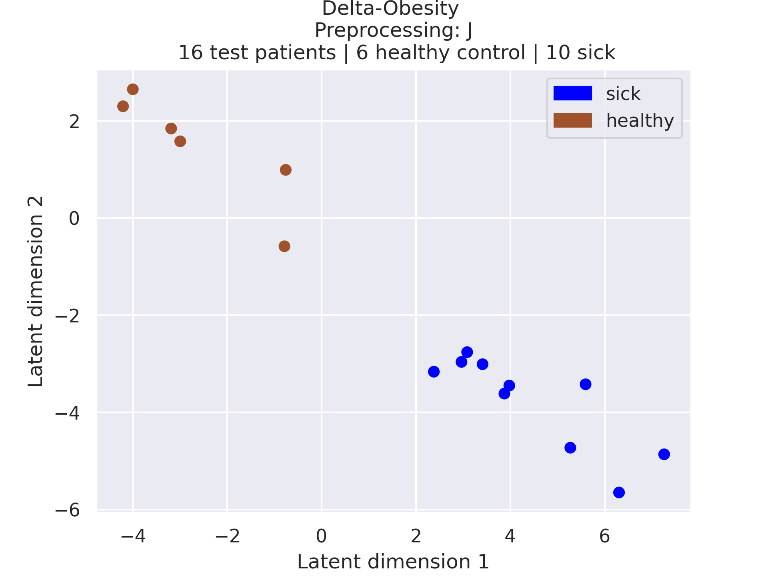

### 3_embeddings.png

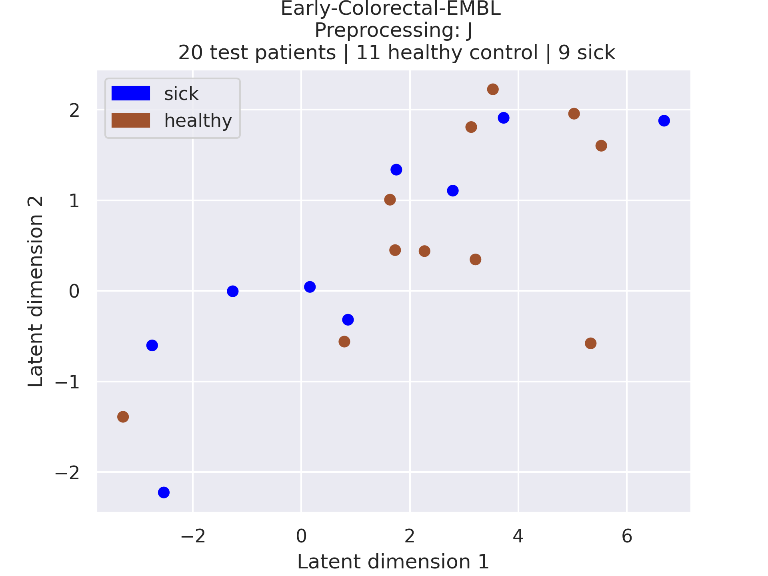

### 3_embeddings.png

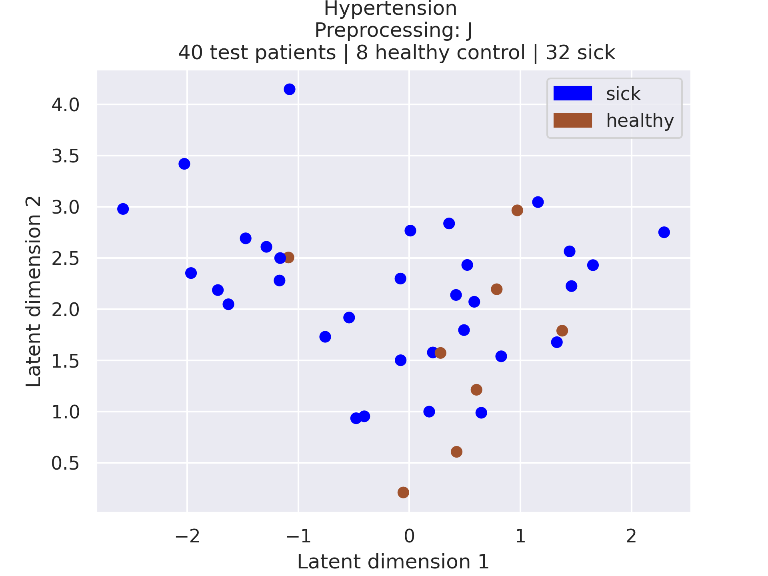

### 3_embeddings.png

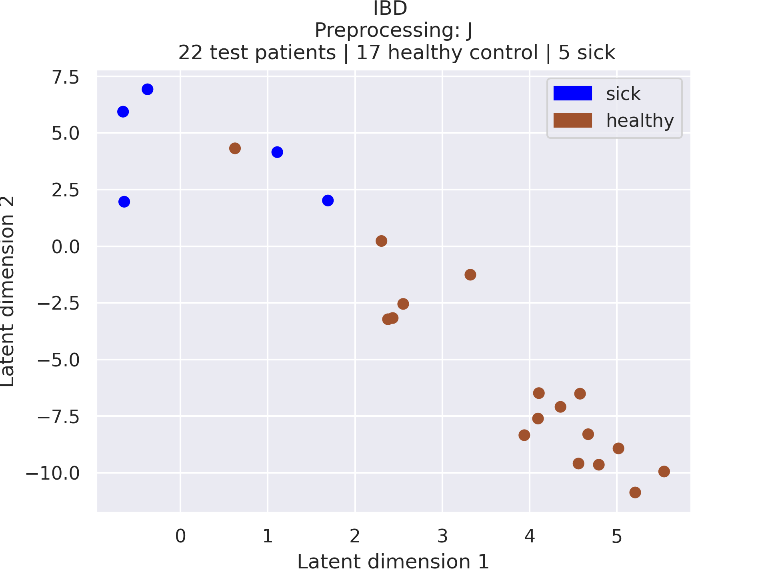

### 3_embeddings_95_confidence.png

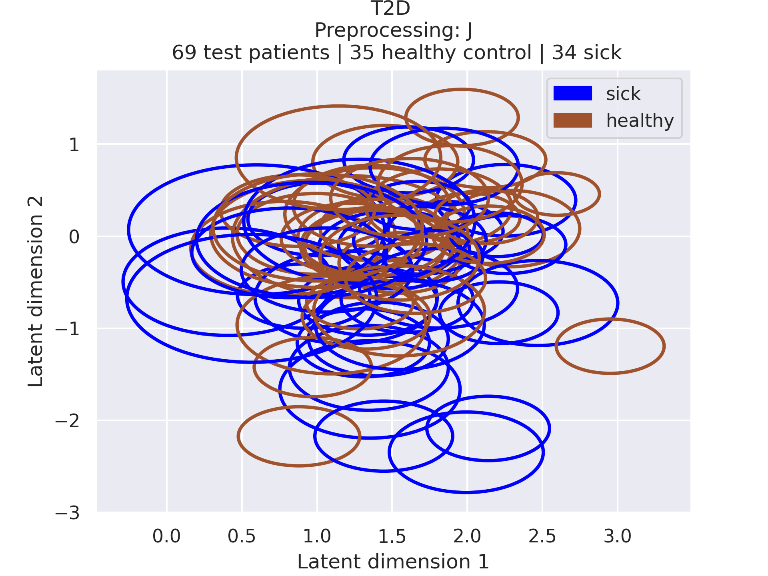

### 3_embeddings_95_confidence.png

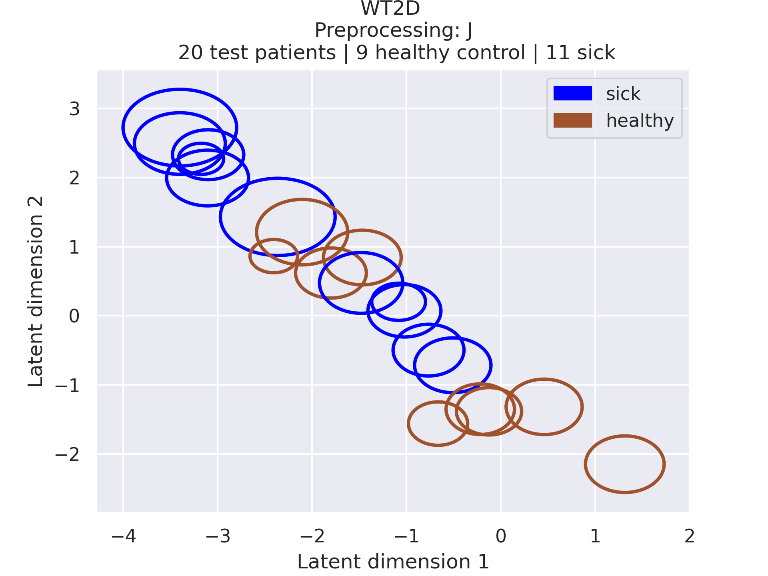

### 3_embeddings_95_confidence.png

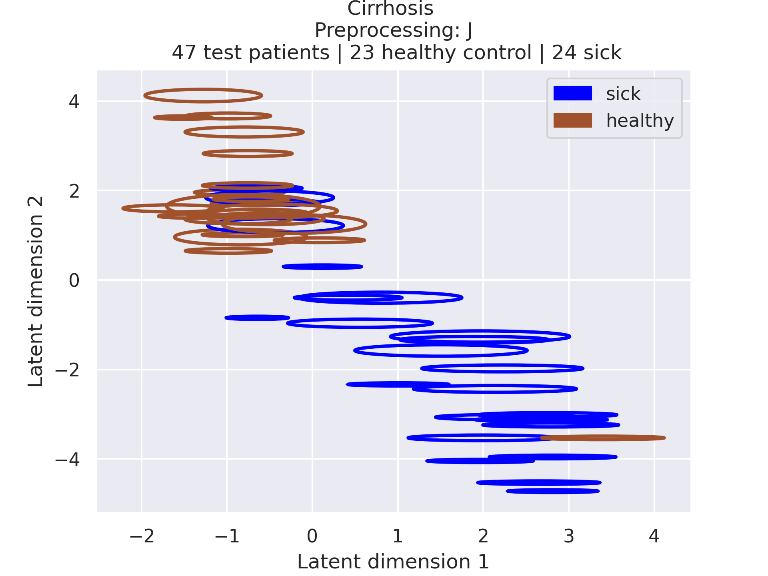

### 3_embeddings_95_confidence.png

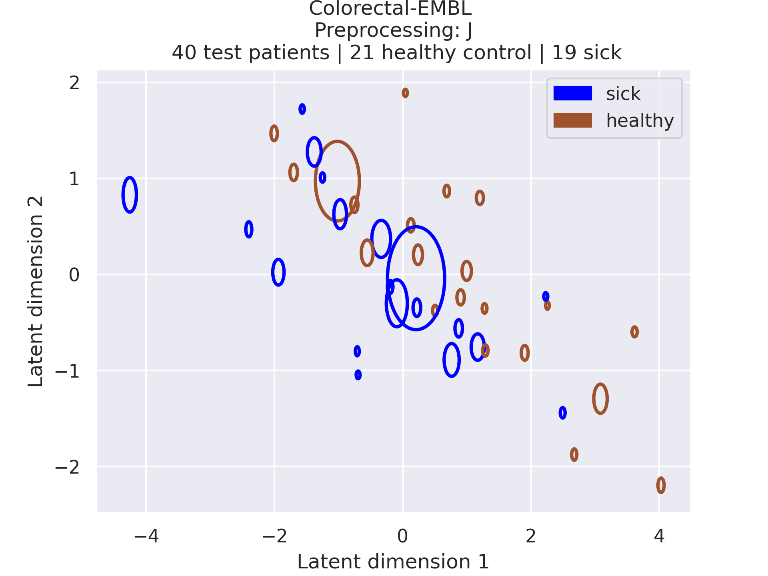

### 3_embeddings_95_confidence.png

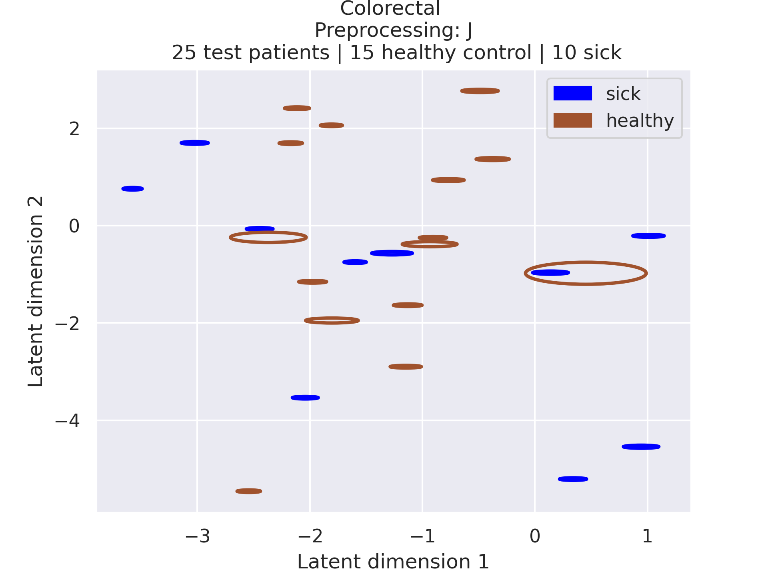

### 3_embeddings_95_confidence.png

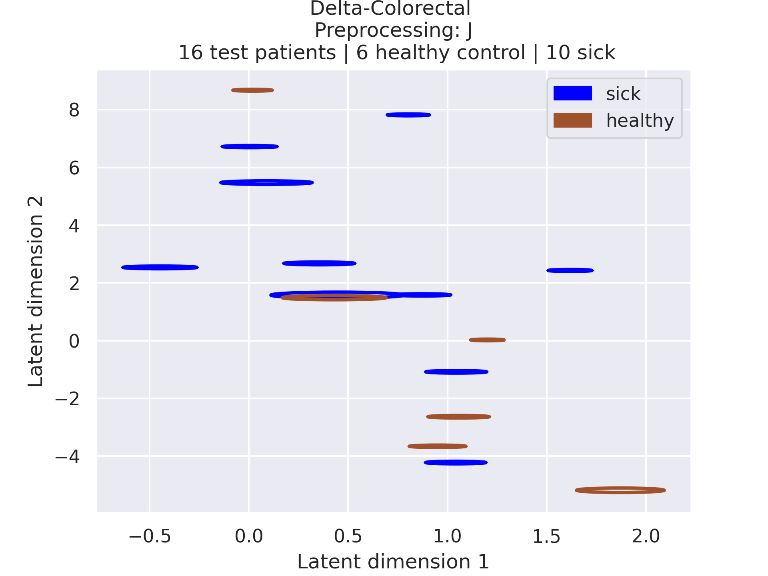

### 3_embeddings_95_confidence.png

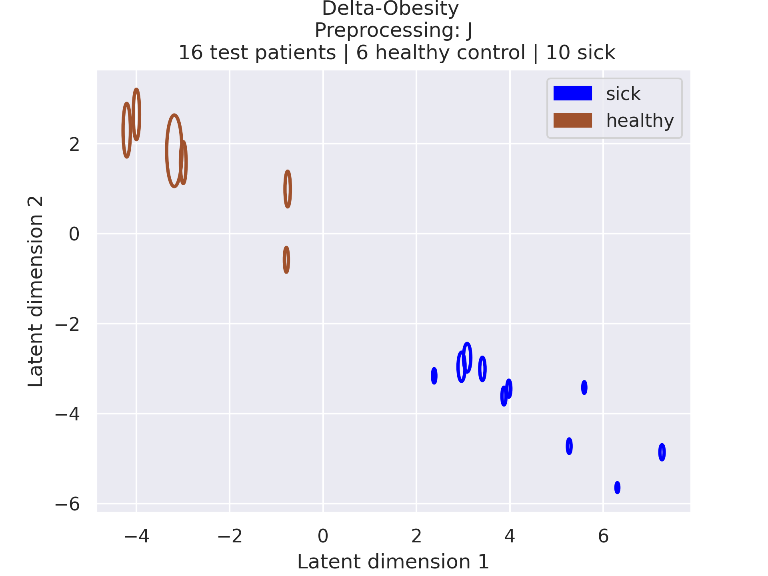

### 3_embeddings_95_confidence.png

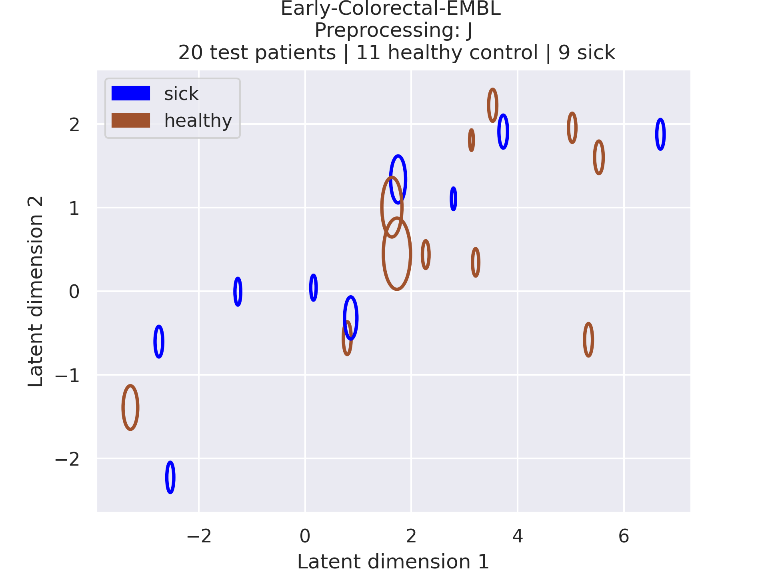

### 3_embeddings_95_confidence.png

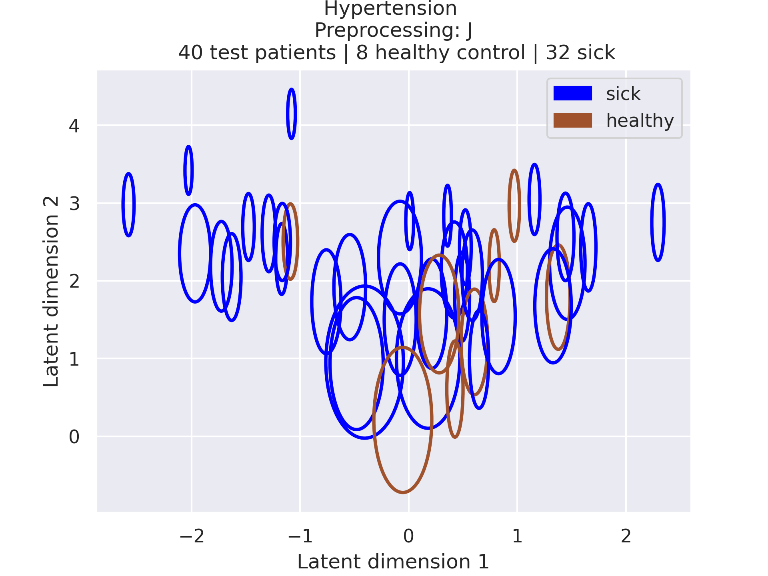

### 3_embeddings_95_confidence.png

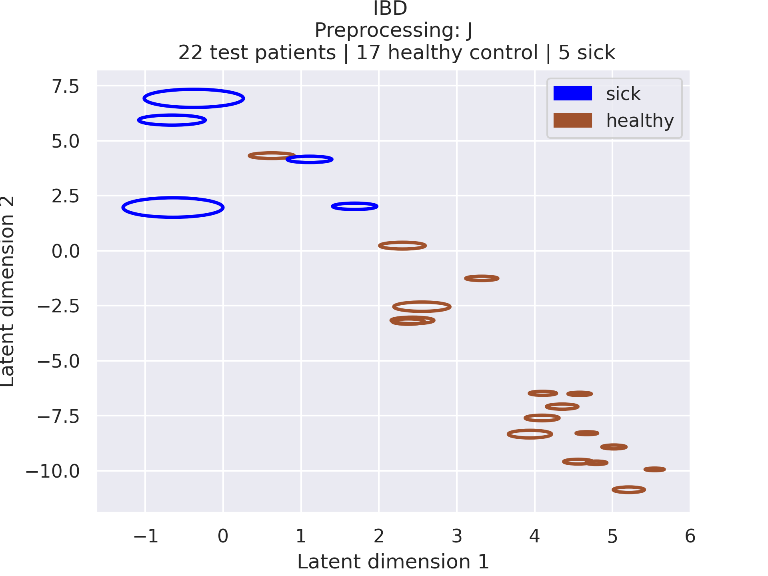

### 4_embeddings.png

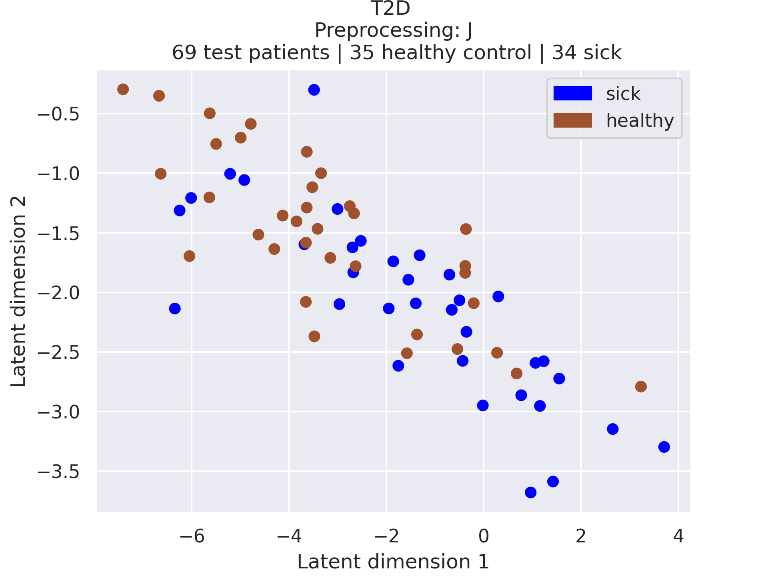

### 4_embeddings.png

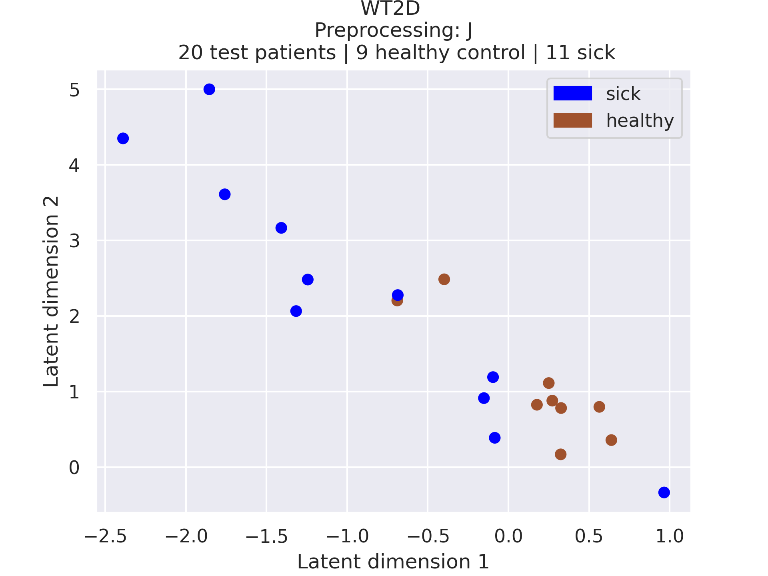

### 4_embeddings.png

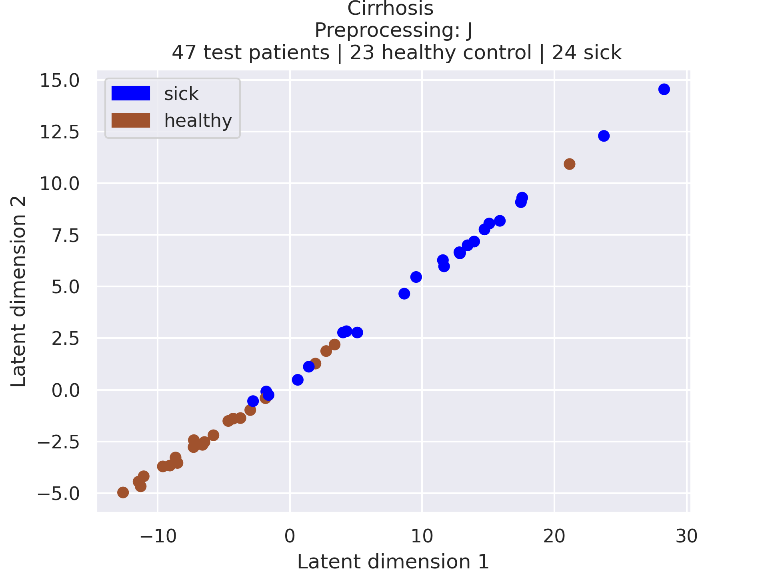

### 4_embeddings.png

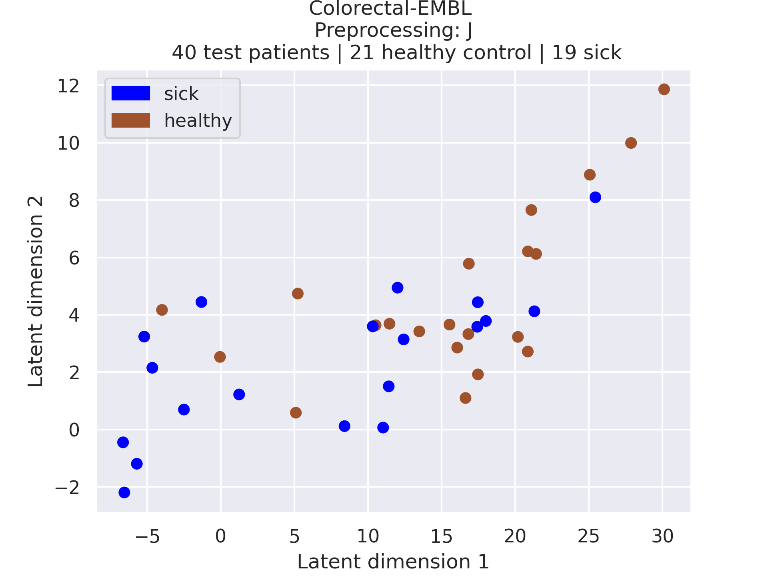

### 4_embeddings.png

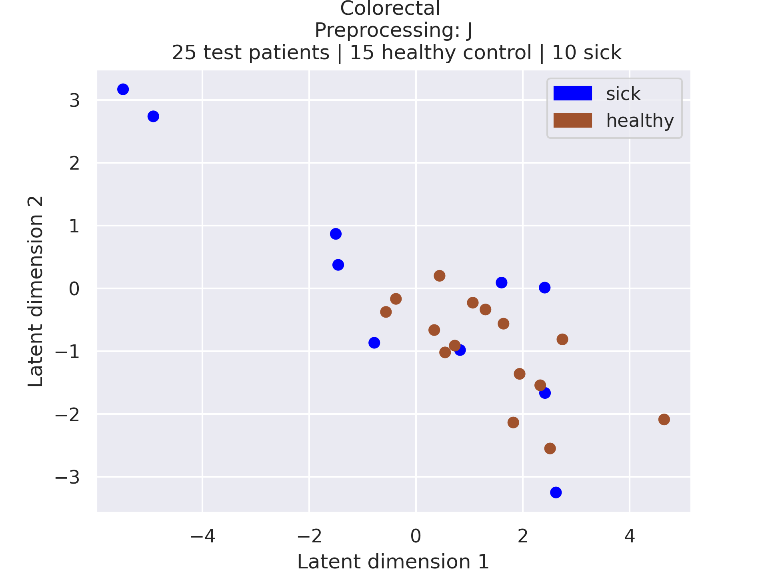

### 4_embeddings.png

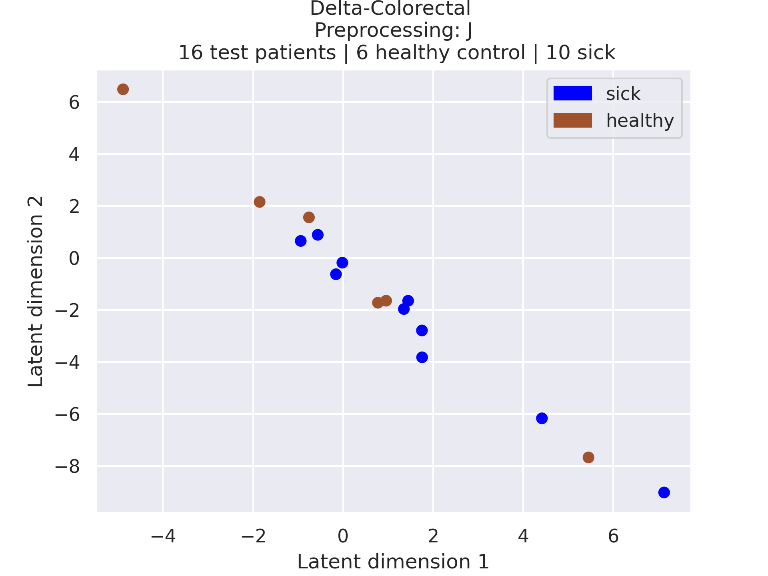

### 4_embeddings.png

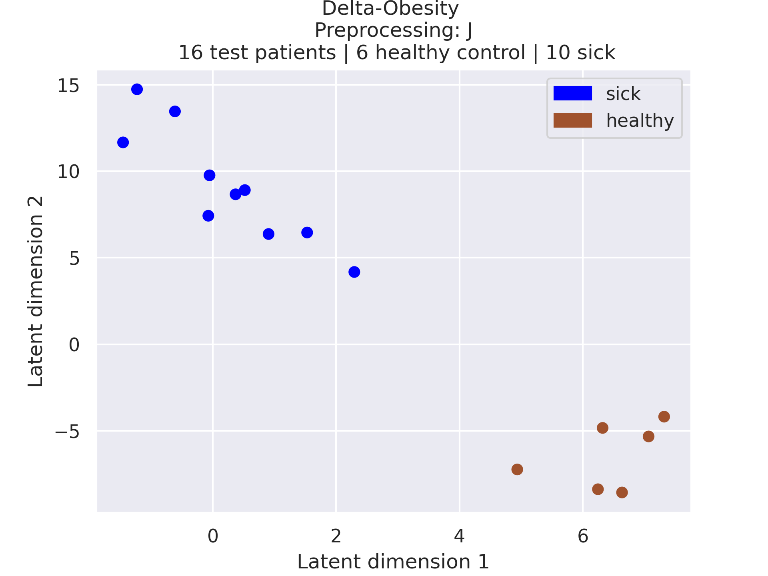

### 4_embeddings.png

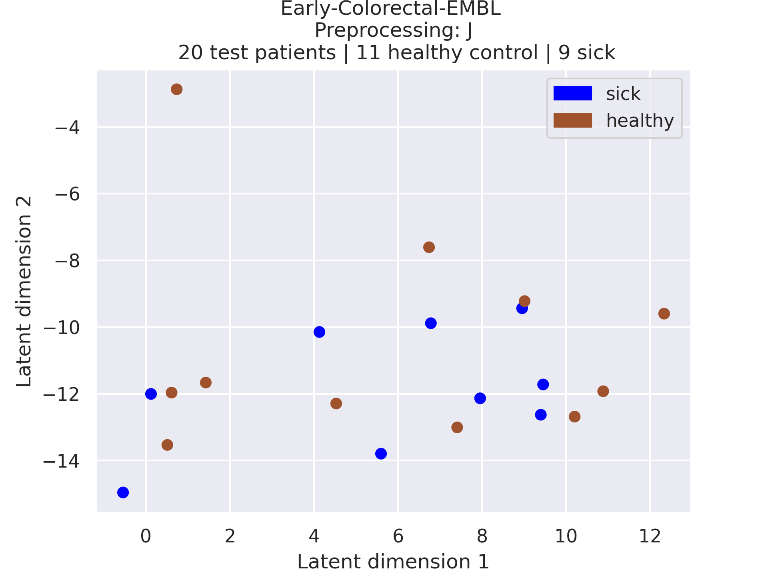

### 4_embeddings.png

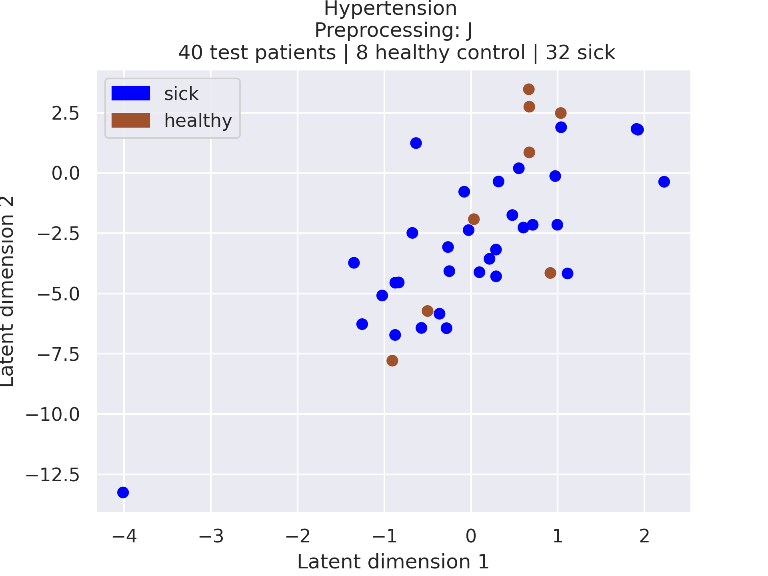

### 4_embeddings.png

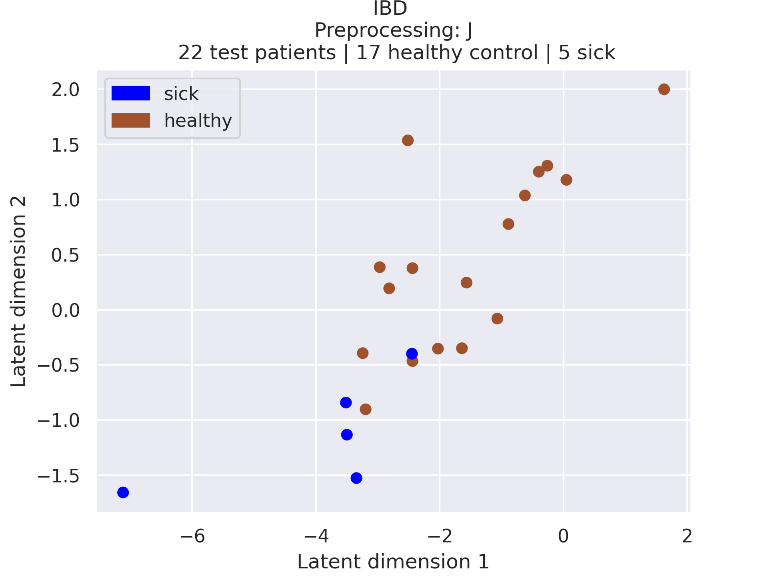

### 4_embeddings_95_confidence.png

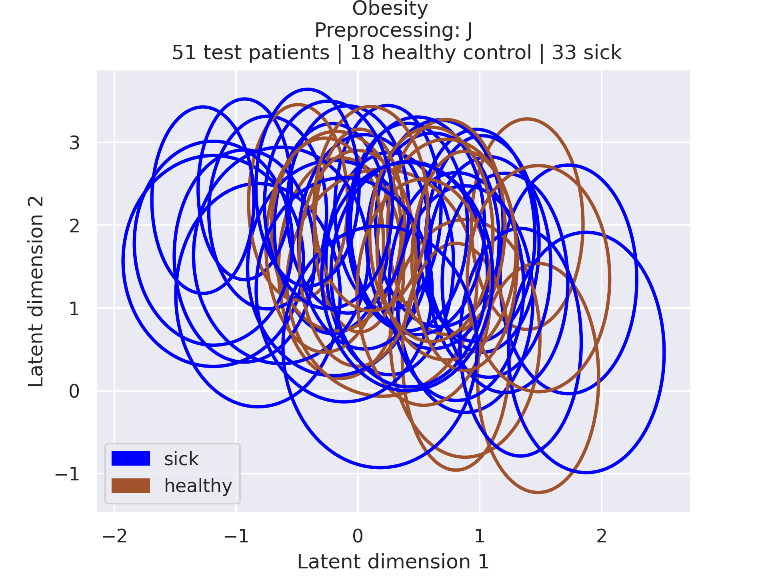

### 4_embeddings_95_confidence.png

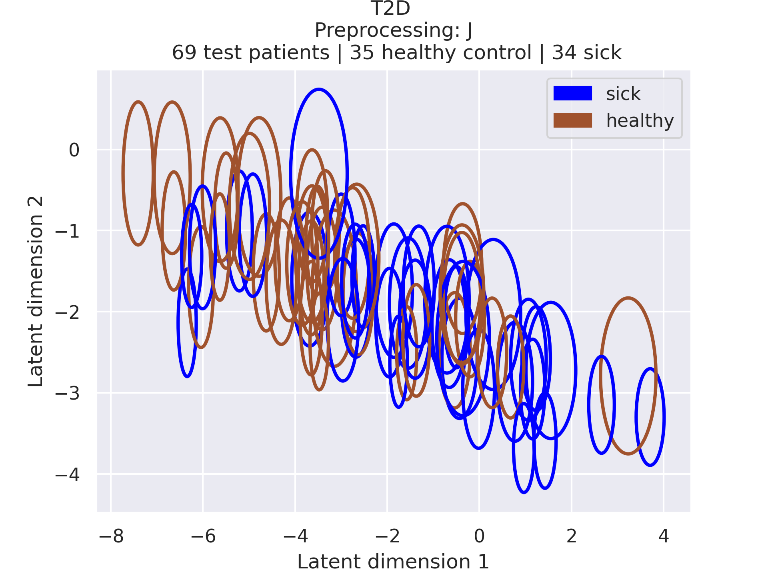

### 4_embeddings_95_confidence.png

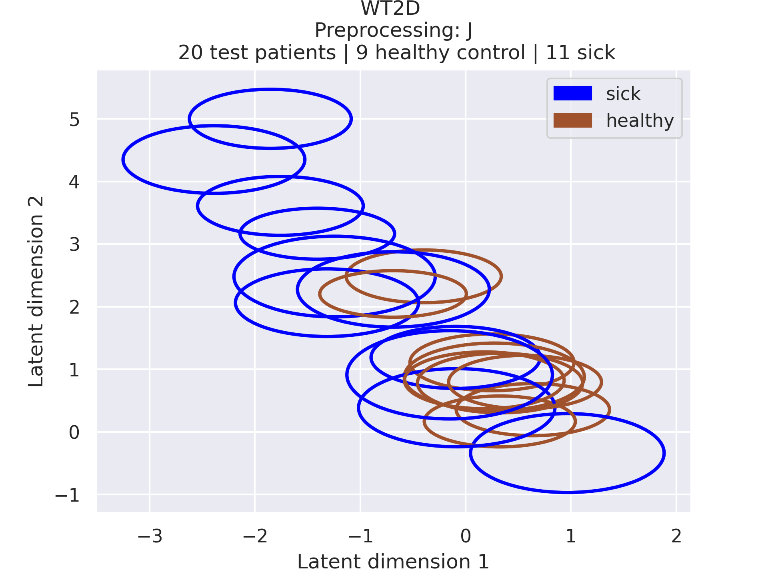

### 4_embeddings_95_confidence.png

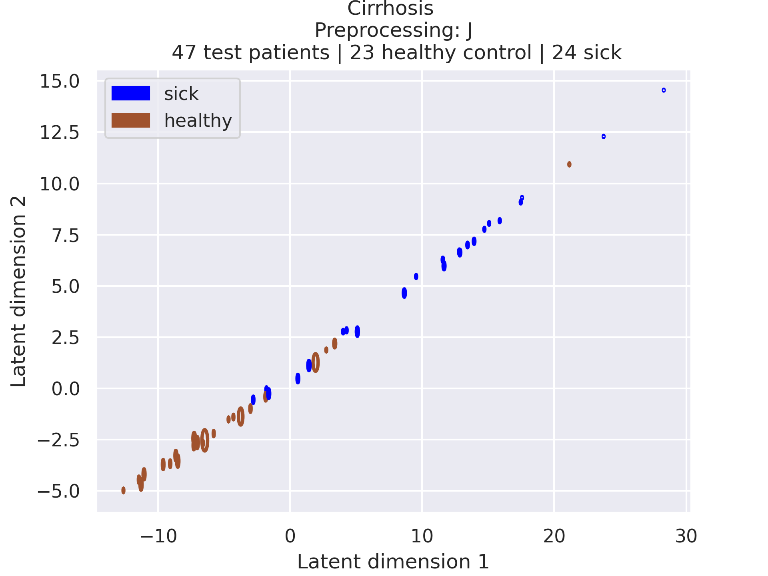

### 4_embeddings_95_confidence.png

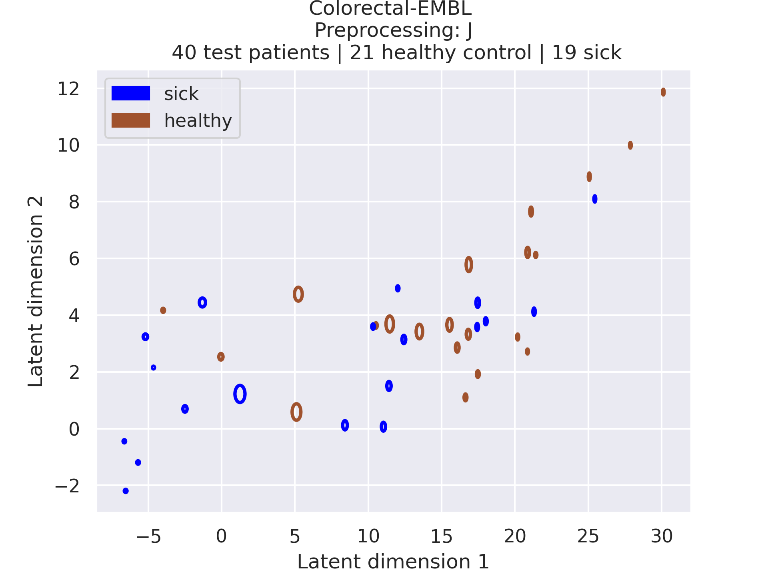

### 4_embeddings_95_confidence.png

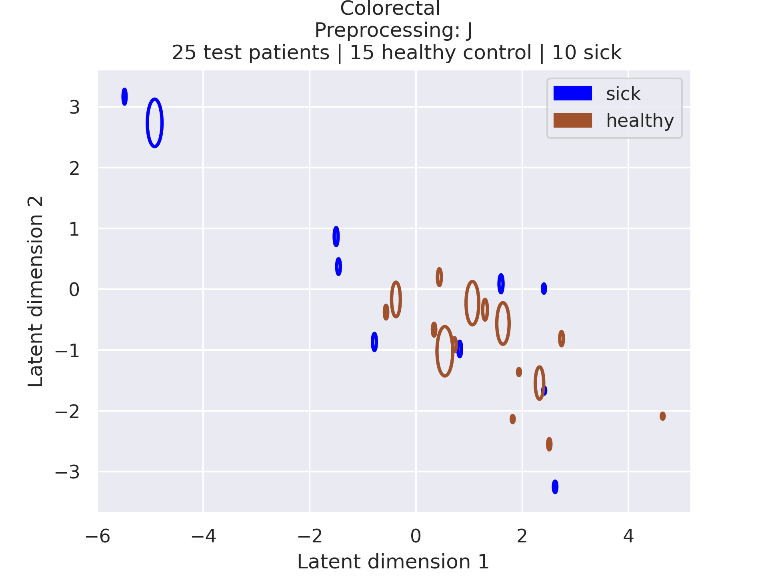

### 4_embeddings_95_confidence.png

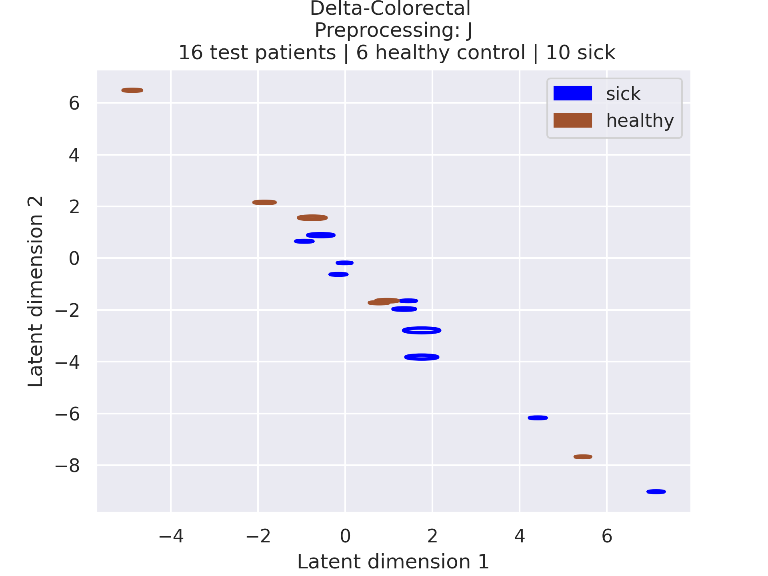

### 4_embeddings_95_confidence.png

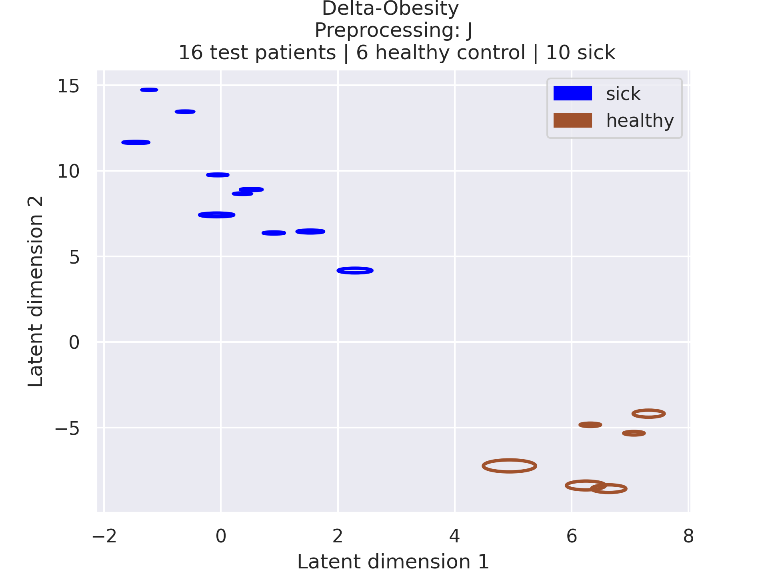

### 4_embeddings_95_confidence.png

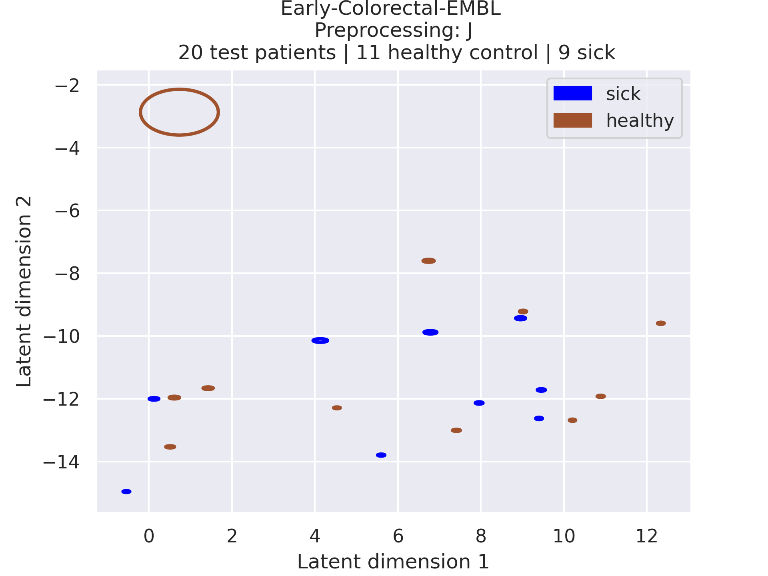

### 4_embeddings_95_confidence.png

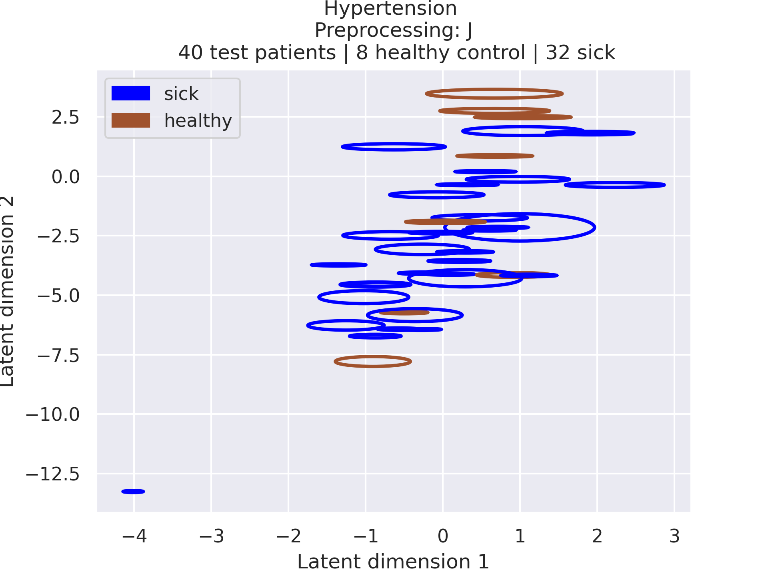
