## Supplementary material for "Microbiome-based disease prediction with multimodal variational information bottlenecks": S7 Table

**S7 Table. Comparison of pre-trained models against randomly initialised models.**

| Dataset | Random initialisation | Pre-trained model |
|---|---|---|
| | J | J |
| IBD | **0.936 (0.014)** | 0.882 (0.021) |
| EW-T2D | **0.853 (0.025)** | 0.780 (0.024) |
| C-T2D | 0.758 (0.012) | **0.774 (0.013)** |
| Obesity | 0.666 (0.027) | **0.679 (0.024)** |
| Cirrhosis | **0.924 (0.005)** | 0.918 (0.007) |
| Colorectal | **0.777 (0.069)** | 0.763 (0.068) |
| Hypertension | 0.591 (0.041) | **0.645 (0.047)** |
