## Supplementary material for "Microbiome-based disease prediction with multimodal variational information bottlenecks": S8 File: check_npy_files.ipynb

```
# %% [code]
import numpy as np

# %% [code]
data_folder = '22-Jul-2021--17-54-36'

datasets = [
#  'Delta-Obesity',
#  'Early-Colorectal-EMBL',
#  'Cirrhosis',
#  'IBD',
#  'Obesity',
#  'Obesity-joint',
# 'Delta-Colorectal',
#  'Colorectal-EMBL',
#  'Hypertension',
#  'T2D',
#  'Colorectal',
  'WT2D',
  'Colorectal-YachidaS'
]

# %% [code]
for dataset in datasets:
    fname = '/Users/nle5293/Documents/1-Projects/1-Microbiome/__paper/saliences/data/'+data_folder+'/'+dataset+'/full_abundance_saliency.npy'
    df = np.load(fname)
    df.shape
    np.save('/Users/nle5293/Documents/1-Projects/1-Microbiome/__paper/saliences/data/'+data_folder+'/'+dataset+'/full_abundance_saliency_.npy', df.astype(np.float64))
    
    fname = '/Users/nle5293/Documents/1-Projects/1-Microbiome/__paper/saliences/data/'+data_folder+'/'+dataset+'/full_labels.npy'
    df = np.load(fname)
    df.shape
    np.save('/Users/nle5293/Documents/1-Projects/1-Microbiome/__paper/saliences/data/'+data_folder+'/'+dataset+'/full_labels_.npy', df.astype(np.float64))
    
    fname = '/Users/nle5293/Documents/1-Projects/1-Microbiome/__paper/saliences/data/'+data_folder+'/'+dataset+'/full_predictions.npy'
    df = np.load(fname)
    df.shape
    np.save('/Users/nle5293/Documents/1-Projects/1-Microbiome/__paper/saliences/data/'+data_folder+'/'+dataset+'/full_predictions_.npy', df.astype(np.float64))
    
    fname = '/Users/nle5293/Documents/1-Projects/1-Microbiome/__paper/saliences/data/'+data_folder+'/'+dataset+'/abundance_saliency_maps.npy'
    df = np.load(fname)
    df.shape
    np.save('/Users/nle5293/Documents/1-Projects/1-Microbiome/__paper/saliences/data/'+data_folder+'/'+dataset+'/abundance_saliency_maps_.npy', df.astype(np.float64))

# %% [code]
for dataset in datasets:
    fname = '/Users/nle5293/Documents/1-Projects/1-Microbiome/__paper/saliences/data/'+data_folder+'/'+dataset+'/full_markers_saliency.npy'
    df = np.load(fname)
    df.shape
    np.save('/Users/nle5293/Documents/1-Projects/1-Microbiome/__paper/saliences/data/'+data_folder+'/'+dataset+'/full_markers_saliency_.npy', df.astype(np.float64))
    
    #fname = '/Users/nle5293/Documents/1-Projects/1-Microbiome/__paper/saliences/data/'+data_folder+'/'+dataset+'/markers_saliency_maps.npy'
    #df = np.load(fname)
    #df.shape
    #np.save('/Users/nle5293/Documents/1-Projects/1-Microbiome/__paper/saliences/data/'+data_folder+'/'+dataset+'/markers_saliency_maps_.npy', df.astype(np.float64))
```
