## Supplementary material for "Microbiome-based disease prediction with multimodal variational information bottlenecks": S8 File: plot_saliencies.R

```
#
# This script plots saliencies for using in PLOS publication
#

library(dplyr)

library(jsonlite)
library(stringr)
library(hash)

library(ggplot2)
library(ggpubr)

library(RcppCNPy)
library(reshape2)


# Globals  ===============================================================

setwd('/Users/nle5293/Documents/1-Projects/1-Microbiome/__paper/massSpec_Abundances')

DATA_FOLDER <- '22-Jul-2021--17-54-36'

# Global paths

ROOT_DATA <- '/Users/nle5293/Documents/1-Projects/1-Microbiome/__paper/data/data'
DIR_DATA <- file.path(ROOT_DATA, 'default')
ROOT_SALIENCY <- '/Users/nle5293/Documents/1-Projects/1-Microbiome/__paper/saliences'
DIR_SALIENCY <- file.path(ROOT_SALIENCY, 'data', DATA_FOLDER)

DIR_PLOTS <- file.path(ROOT_SALIENCY, 'plots', DATA_FOLDER)
dir.create(file.path(ROOT_SALIENCY, 'plots'))
dir.create(file.path(DIR_PLOTS))

# Select dataset

datasets = c(
  'Delta-Obesity',
  'Early-Colorectal-EMBL',
  'Cirrhosis',
  'IBD',
  'Obesity',
  'Obesity-joint',
  'Delta-Colorectal',
  'Colorectal-EMBL',
  'Hypertension',
  'T2D',
  'Colorectal',
  'WT2D',
  'Colorectal-YachidaS'
)
dataset <- datasets[13] # select dataset manually

# Other globals
start_row <- 211  # start data body in input dataframes
n_top_ <- 25  # default N top species to output
n_rep_experiments <- 25 # total number of repeated experiments
class_one_threshold <- 0.5  # current threshold to convert predictions to class 1
p_thr_ <- 0.1 # p-value threshold in Wilcoxon test
use_p_fdr_ <- TRUE # use results with FDR correction from Wilcoxon test
sort_mode_ <- 'Mean' # sort saliencies by mean values


# Create label dictionary for healthy/affected status in the dataset

label_dict = hash()
  # Controls
label_dict[['n']] = 0
  # Chirrhosis
label_dict[['cirrhosis']] = 1
  # Colorectal Cancer
label_dict[['cancer']] = 1 
label_dict[['small_adenoma']] = 0
  # IBD
label_dict[['ibd_ulcerative_colitis']] = 1 
label_dict[['ibd_crohn_disease']] = 1
# T2D and WT2D
label_dict[['t2d']] = 1
# Obesity / Obesity-joint
label_dict[['leaness']] = 0 
label_dict[['obesity']] = 1
# Early-Colorectal-EMBL (early = 0, I, II; late = III, IV)
label_dict[['crc_0']] = 0 
label_dict[['crc_I']] = 0
label_dict[['crc_II']] = 0
label_dict[['crc_III']] = 1
label_dict[['crc_IV']] = 1
# Colorectal-EMBL (healthy = n)
label_dict[['CRC_0']] = 1
label_dict[['CRC_I']] = 1
label_dict[['CRC_II']] = 1
label_dict[['CRC_III']] = 1
label_dict[['CRC_IV']] = 1
# Hypertension
label_dict[['Health']] = 0
label_dict[['Hypertension']] = 1
# Custom
label_dict[['sick']] = 1
# YachidaS
label_dict[['healthy']] = 0
label_dict[['CRC-0']] = 1
label_dict[['CRC-I']] = 1
label_dict[['CRC-II']] = 1
label_dict[['CRC-III']] = 1
label_dict[['CRC-IV']] = 1
label_dict[['adenoma']] = 1
label_dict[['carcinoma_surgery_history']] = 1


# Function definitions ======================================================

GetSpeciesNames <- function(data) {
  # Get substring with species names from whole taxa branch.
  #
  # Args:
  #   data: Column of dataframe that contains taxa paths.
  #
  # Returns:
  #   Vector of strings with species names.
  
  species <- as.vector(sapply(
    data, 
    function(in_str)
      ifelse(str_split_fixed(in_str, ".s__", 2)[2] != "", 
             str_split_fixed(in_str, ".s__", 2)[2], 
             str_split_fixed(in_str, ".g__", 2)[2])))
  
  return(species)
}


TestRepresentedSpecies <- function(df_abundance, p_thr = 0.05, use_p_fdr = TRUE) {
  # Perform Wilcoxon test to identify over- and underrepresented species.
  #
  # Args:
  #   df_abundance: Dataframe with abundances.
  #   p_thr: P-value threshold.
  #   use_p_fdr: Use adjusted p-values.
  #
  # Returns:
  #   Dataframe with statistical test results for the species.  
  
  # Split head and body
  df_header <- df_abundance[1:start_row-1, ]
  df_body <- df_abundance[start_row:nrow(df_abundance), ]
  
  # Get status of the patients
  patient_labels <- df_header[df_header[, 1]=='disease', 2:ncol(df_header)]
  patient_labels <- unlist(lapply(patient_labels, function(data) label_dict[[data]]))
  groups <- factor(patient_labels, levels = c(0, 1))
  
  # Create dataframe with summary statistics on patient groups
  #df_stats <- data.frame(row.names = str_split_fixed(df_body[, 1], ".s__", 2)[,2])
  df_stats <- data.frame(row.names = GetSpeciesNames(df_body[, 1]))
  df_stats$check_names <- df_body[, 1]
  
  # Perform Wilcoxon test
  print("Performing Wilcoxon test to find overrepresented species...")
  
  # Over-represented in affected group
  df_stats$W_greater_pval <- apply(
    df_body[, 2:ncol(df_body)], 
    MARGIN = 1, 
    FUN = function(x_line)
      wilcox.test(
        as.numeric(unlist(x_line[groups==1])),
        as.numeric(unlist(x_line[groups==0])),
        alternative = "greater"
      )$p.value
    )
  
  # Over-represented in health group
  df_stats$W_less_pval <- apply(
    df_body[, 2:ncol(df_body)],
    MARGIN = 1,
    FUN = function(x_line)
      wilcox.test(
        as.numeric(unlist(x_line[groups==1])),
        as.numeric(unlist(x_line[groups==0])),
        alternative = "less"
      )$p.value
    )                    
  
  # Two-side test
  df_stats$W_twoside_pval <- apply(
    df_body[, 2:ncol(df_body)],
    MARGIN = 1,
    FUN = function(x_line)
      wilcox.test(
        as.numeric(unlist(x_line[groups==0])),
        as.numeric(unlist(x_line[groups==1])),
        alternative = "two.sided"
      )$p.value
    )                    
  
  # Adjust W-test p-values for multiple comparisons
  df_stats$W_gt_pval_fdr <- p.adjust(df_stats$W_greater_pval, method = "fdr")
  df_stats$W_ls_pval_fdr <- p.adjust(df_stats$W_less_pval, method = "fdr")
  df_stats$W_twoside_pval_fdr <- p.adjust(df_stats$W_twoside_pval, method = "fdr")
  
  # Columns with colors for over- and under-represented species
  if (use_p_fdr == TRUE) {
    df_stats$representation <- 
      if_else(
        df_stats$W_gt_pval_fdr < p_thr, "prevail in affected",
                  if_else(df_stats$W_ls_pval_fdr < p_thr, "prevail in healthy", "nd")
      )
  } else {
    df_stats$representation <- 
      if_else(
        df_stats$W_greater_pval < p_thr, "prevail in affected",
                  if_else(df_stats$W_less_pval < p_thr, "prevail in healthy", "nd")
      )
  }
  
  return(df_stats)
}


SymLogRescale <- function(base = 10, thr = 1e-10, scale = 1, zoom = NULL) {
  # Rescale plot output into pseudo-logarithmic scale.
  #
  # Args:
  #   base: Base of the logarithms.
  #   thr: For x < thr plot without transforms, for x > thr plot in log scale.
  #
  # Returns:
  #   Function for scale_x/y_continuous()
  #
  #thr = 1e-8
  
  trans <- function(x)
    ifelse(abs(x) < thr, x, sign(x) * 
             (scale * suppressWarnings(log(sign(x) * x / thr, base))))
  
  inv <- function(x)
    ifelse(abs(x) < thr, x, sign(x) * 
             base^((sign(x) * x) / scale) * thr)
  
  breaks <- function(x) {
    sgn <- sign(x[which.max(abs(x))])
    sgn * unique(c(scales::log_breaks(base)(c(min(abs(x)), max(abs(x))))))
  }
    
  scales::trans_new(paste("symlog", thr, base, scale, sep = "-"), trans, inv, breaks)
}


PlotSaliencyMap <- function(df_saliencies, df_stats, sort_mode = 'Mean', n_top = 25, 
                            rescale_plot = TRUE, plot_title = "", plot_lims = NULL) {
  # Builda plots from saliency maps.
  #
  # Args:
  #   df_saliencies: Dataframe with saliency map of size n_experiments x n_features.
  #   df_stats: Dataframe with marks of over- and under-represented species.
  #   sort_mode: How to select and sort top saliencies, one of {'Mean', 'Median'}.
  #
  # Returns:
  #   Plot objects.
  
  # Calculate summary statistics
  df_saliencies_summary <- as.data.frame(t(do.call(cbind, lapply(df_saliencies, summary))))
  df_saliencies_summary <- merge(df_saliencies_summary, df_stats['representation'], by.x = 0, by.y = 0, all.x = TRUE, all.y = FALSE)
  rownames(df_saliencies_summary) <- df_saliencies_summary[,'Row.names']
  df_saliencies_summary <- df_saliencies_summary[order(df_saliencies_summary[,sort_mode], decreasing = TRUE),]

  # Select top species with highest mean saliencies
  top_species <- row.names(df_saliencies_summary[1:n_top,])
  
  # Vector of means
  mean_vec <- df_saliencies_summary[1:n_top, 'Mean']
  zoom_range <- c(min(mean_vec)-sd(mean_vec), max(mean_vec)+sd(mean_vec))
  
  # Draw manual boxplot from summary statistics
  p_summaries <- ggplot() +
    geom_boxplot(
      aes(
        x = reorder(top_species, df_saliencies_summary[1:n_top, sort_mode]),
        ymin = df_saliencies_summary[1:n_top,'Min.'], 
        lower = df_saliencies_summary[1:n_top,'1st Qu.'],                    
        middle = df_saliencies_summary[1:n_top, 'Median'], 
        upper = df_saliencies_summary[1:n_top,'3rd Qu.'], 
        ymax = df_saliencies_summary[1:n_top,'Max.'],
        fill = df_saliencies_summary[1:n_top,'representation']), 
      stat = "identity", 
      width = .5
    ) +
    geom_point(
      aes(x = row.names(df_saliencies_summary[1:n_top,]), y = df_saliencies_summary[1:n_top,'Mean']),
      shape = 5, 
      colour = "black"
    ) +
    scale_fill_manual("", values = c("nd" = "grey", "prevail in affected" = "red", "prevail in healthy" = "blue")) +
    labs(
      title = plot_title, 
      x = paste("Top", n_top, "species by", tolower(sort_mode), "saliency"), 
      y = "Saliency"
    )
  if (rescale_plot == TRUE)
    p_summaries <- p_summaries + scale_y_continuous(trans=SymLogRescale(zoom = zoom_range))

  
  # Melt dataframe for other plots
  df_melt <- melt(df_saliencies[,top_species], id.vars = NULL)
  df_melt <- merge(df_melt, df_stats['representation'], by.x = 1, by.y = 0, all.x = TRUE, all.y = FALSE)
  
  
  # Basic ggplot for boxplot and violine plot
  if (sort_mode == 'Mean') {
    p_ggplot <- ggplot(
      df_melt, 
      aes(x = reorder(variable, value, mean), y = value, fill = representation)
    )
  } else {
    p_ggplot <- ggplot(
      df_melt, 
      aes(x = reorder(variable, value, median), y = value, fill = representation)
    )
  }
  
  # Draw boxplot
  p_boxplot <- p_ggplot + 
    geom_boxplot(outlier.shape = NA) + 
    stat_summary(
      fun = function(x) 
              ifelse(rescale_plot == TRUE, log10(mean(10^x)), mean(x)), 
      geom = "point", shape = 5, colour = "black"
    ) +
    scale_fill_manual("", values = c("nd" = "grey", "prevail in affected" = "red", "prevail in healthy" = "blue")) +
    labs(
      title = plot_title, 
      x = paste("Top", n_top, "species by", tolower(sort_mode), "saliency"), 
      y = "Saliency"
    )
  if (rescale_plot == TRUE)
    p_boxplot <- p_boxplot + scale_y_continuous(trans=SymLogRescale())

  
  # Draw violine plot
  p_violine <- p_ggplot +
    geom_violin(draw_quantiles = c(0.25, 0.5, 0.75),) + 
    stat_summary(
      fun = function(x) 
        ifelse(rescale_plot == TRUE, log10(mean(10^x)), mean(x)), 
      geom = "point", shape = 5, colour = "black"
    ) +
    scale_fill_manual("", values = c("nd" = "grey", "prevail in affected" = "red", "prevail in healthy" = "blue")) +
    labs(
      title = plot_title, 
      x = paste("Top", n_top, "species by", tolower(sort_mode), "saliency"), 
      y = "Saliency"
    )
  if (rescale_plot == TRUE)
    p_violine <- p_violine + scale_y_continuous(trans=SymLogRescale())
  
  if (is.null(plot_lims))
    p_list <- list("summary_boxplot" = p_summaries + coord_flip(), 
                   "boxplot" = p_boxplot + coord_flip(), 
                   "violine_plot" = p_violine + coord_flip())
  else
    p_list <- list("summary_boxplot" = p_summaries + coord_flip(ylim = plot_lims), 
                   "boxplot" = p_boxplot + coord_flip(ylim = plot_lims), 
                   "violine_plot" = p_violine + coord_flip(ylim = plot_lims))
}


# Start pipeline ======================================================

# Calculate under- and over-represented species
in_filename <- file.path(DIR_DATA, "abundance", paste("abundance_", dataset, ".txt", sep = ""))

df_abundance <- read.csv(in_filename, sep="\t")
df_stats <- TestRepresentedSpecies(df_abundance, p_thr = p_thr_, use_p_fdr = use_p_fdr_)

# Plot saliences.
# Saliencies in (1) were plotted from maps without re-scaling.
# Saliencies in (2)-(5) were plotted from scales values.

print("Generate plots...")

dir.create(file.path(DIR_PLOTS, dataset))


# (0) Plot saliency maps for true positives (full maps, BATCH RESCALING).

fname <- file.path(DIR_SALIENCY, dataset, "full_abundance_saliency_.npy")
if (file.exists(fname)) {
  print("Generate plots from full saliency maps.")
  
  df_full_saliencies <- as.data.frame(npyLoad(fname))
  #colnames(df_full_saliencies)<- str_split_fixed(df_abundance[start_row:nrow(df_abundance),1], ".s__", 2)[,2]
  colnames(df_full_saliencies) <- GetSpeciesNames(df_abundance[start_row:nrow(df_abundance),1])
  dim_saliency_map <- dim(df_full_saliencies)
  print(paste("Dimension of saliency map:", dim_saliency_map[[1]], "x", dim_saliency_map[[2]]))
  
  # Rescale saliency maps by batch (in total there are n_experiment batches)
  batch_size <- dim_saliency_map[[1]] / n_rep_experiments

  # assert(n_batch_size == ncol(df_abundance)-1)
  for (batch_idx in 1:n_rep_experiments) {
    start_idx <- as.integer((batch_idx-1)*batch_size+1)
    end_idx <- as.integer(batch_idx*batch_size)
    df_full_saliencies[start_idx:end_idx,] <- 
      (df_full_saliencies[start_idx:end_idx,]-min(df_full_saliencies[start_idx:end_idx,]))/(max(df_full_saliencies[start_idx:end_idx,])-min(df_full_saliencies[start_idx:end_idx,]))
  }
  
  fname_labels <- file.path(DIR_SALIENCY, dataset, "full_labels_.npy")
  df_full_labels <- as.data.frame(npyLoad(fname_labels))
  
  fname_predictions <- file.path(DIR_SALIENCY, dataset, "full_predictions_.npy")
  df_full_predictions <- as.data.frame(npyLoad(fname_predictions))
  
  # Convert predictions to 0/1 labels
  df_full_predictions_lb_ <- df_full_predictions
  df_full_predictions_lb_[] <- lapply(
    df_full_predictions_lb_, function(x) ifelse(x>class_one_threshold, 1, 0))
  
  # Get all true positive experiments
  tp_experiments_ind <- which(c(df_full_predictions_lb_)$V1 == c(df_full_labels)$V1, arr.ind = TRUE)
  df_tp_saliences <- df_full_saliencies[tp_experiments_ind,]
  
  # Just for check
  arr_pos_predictions <- df_full_predictions[tp_experiments_ind,]
  arr_pos_labels <- df_full_labels[tp_experiments_ind,]
  
  # Plot saliencies
  plots_tp_saliency <- PlotSaliencyMap(
    df_tp_saliences, df_stats, sort_mode = 'Mean', n_top = n_top_, rescale_plot = FALSE, plot_lims = c(0, 0.02))
  
  # Combine all plots and save
  figure0 <- annotate_figure(
    ggarrange(
      plots_tp_saliency[[1]] + theme(
        plot.title = element_text(hjust = 0, face= "bold"), 
        plot.title.position = "plot",
        legend.position = "bottom", 
        plot.margin = margin(2, 1, 0, 1, unit = 'line')
      ), 
      ggarrange(
        plots_tp_saliency[[2]] + theme(
          plot.title = element_text(hjust = 0, face= "bold"), 
          plot.title.position = "plot",
          legend.position = "bottom", 
          plot.margin = margin(2, 1, 0, 1, unit = 'line')
        ), 
        plots_tp_saliency[[3]] + theme(
          plot.title = element_text(hjust = 0, face= "bold"), 
          plot.title.position = "plot",
          legend.position = "bottom", 
          plot.margin = margin(2, 1, 0, 1, unit = 'line')
        ),
        ncol = 2,
        legend = "none",
        labels = c("B.", "C.")
      ),
      nrow = 2,
      common.legend = TRUE, 
      legend = "bottom",
      labels = "A."
    ),
    top = text_grob(
      paste("Saliencies from TP maps for", dataset), 
      color = "black", face = "bold", size = 16)
  )
  #figure0
  fname <- file.path(DIR_PLOTS, dataset, paste("abundance_plots_TP_saliencies_rescaled_all_pval-", p_thr_, ".png", sep = ""))
  ggsave(figure0, width = 20, height = 14, dpi = 300, bg = 'white', filename = fname)
  print(paste("Plots from TP saliency map saved into", fname))
  
  # Save plots separately
  
  figure0A <- annotate_figure(
    plots_tp_saliency[[1]],
    top = text_grob(
      paste("Saliencies from TP maps for", dataset), 
      color = "black", face = "bold", size = 16)
  )
  fname <- file.path(DIR_PLOTS, dataset, paste("abundance_plots_TP_saliencies_rescaled_summaries_pval-", p_thr_, ".png", sep = ""))
  ggsave(figure0A, width = 12, height = 8, dpi = 300, bg = 'white', filename = fname)
  print(paste("Plot from summary statistics from TP saliency map saved into", fname))
  
  figure0B <- annotate_figure(
    plots_tp_saliency[[2]],
    top = text_grob(
      paste("Saliencies from TP maps for", dataset), 
      color = "black", face = "bold", size = 16)
  )
  fname <- file.path(DIR_PLOTS, dataset, paste("abundance_plots_TP_saliencies_rescaled_boxplot_pval-", p_thr_, ".png", sep = ""))
  ggsave(figure0B, width = 12, height = 8, dpi = 300, bg = 'white', filename = fname)
  print(paste("Boxplot from TP saliency map saved into", fname))
  
  figure0C <- annotate_figure(
    plots_tp_saliency[[3]],
    top = text_grob(
      paste("Saliencies from TP maps for", dataset), 
      color = "black", face = "bold", size = 16)
  )
  fname <- file.path(DIR_PLOTS, dataset, paste("abundance_plots_TP_saliencies_rescaled_violine_pval-", p_thr_, ".png", sep = ""))
  ggsave(figure0C, width = 12, height = 8, dpi = 300, bg = 'white', filename = fname)
  print(paste("Violine plot from TP saliency map saved into", fname))
}

# -------------------------------------------------------------------------
# (1) Plot from mean/std saliencies

fname1 <- file.path(DIR_SALIENCY, dataset, "abundance_saliency.json")
fname2 <- file.path(DIR_SALIENCY, dataset, "abundance_std.json")
if (file.exists(fname1) & file.exists(fname2)) {
  # Read values from JSONs
  in_json <- readChar(fname1, file.info(fname1)$size)
  df_saliencies_mean <- t(as.data.frame(fromJSON(in_json)))
  in_json <- readChar(fname2, file.info(fname2)$size)
  df_saliencies_std <- t(as.data.frame(fromJSON(in_json)))
  
  df_saliencies <- merge(df_saliencies_mean, df_saliencies_std, by.x = 0, by.y = 0, all.x = TRUE, all.y = TRUE)
  colnames(df_saliencies) <- c('species', 'mean', 'std')
  df_saliencies$se <- df_saliencies$std / sqrt(dim_saliency_map[1])
  df_saliencies$species <- GetSpeciesNames(df_saliencies$species)
  
  # Add microbe representation values and sort by mean
  df_saliencies <- merge(df_saliencies, df_stats, by.x = 1, by.y = 0, all.x = TRUE, all.y = TRUE)
  df_saliencies <- df_saliencies[order(df_saliencies$mean, decreasing = TRUE), ][1:n_top_,]
  
  # Draw barplot
  p_barplot_err <- ggplot(
    df_saliencies,
    aes(x = reorder(species, mean), y = mean, fill = representation)
  ) +
  geom_col(aes(), width = 0.7) + 
  geom_errorbar(
    aes(ymin = mean - se, ymax = mean + se), width = .2, position = position_dodge(.9)
  ) +
  scale_fill_manual("", values = c("nd" = "grey", "prevail in affected" = "red", "prevail in healthy" = "blue")) +
  labs(
    title = paste("Mean saliencies for", dataset, "(positive predictions)"), 
    x = paste("Top", n_top_, "species by mean saliency"), 
    y = "Saliency"
  ) +
  theme(
    plot.title = element_text(hjust = 0.5, face= "bold"), 
    plot.title.position = "plot",
    legend.position = "bottom", 
    plot.margin = margin(2, 1, 0, 1, unit = 'line')
  ) + 
  coord_flip()
  p_barplot_err  
}

fname <- file.path(DIR_PLOTS, dataset, "abundance_plots_POS_saliencies_barplot.png")
ggsave(p_barplot_err, width = 12, height = 8, dpi = 300, bg = 'white', filename = fname)
print(paste("Error barplot from mean saliencies saved into", fname))


# -------------------------------------------------------------------------
# (2) Plot saliency maps for true positives (full maps, no rescaling).

fname <- file.path(DIR_SALIENCY, dataset, "full_abundance_saliency_.npy")
if (file.exists(fname)) {
  print("Generate plots from full saliency maps.")
  
  df_full_saliencies <- as.data.frame(npyLoad(fname))
  #colnames(df_full_saliencies)<- str_split_fixed(df_abundance[start_row:nrow(df_abundance),1], ".s__", 2)[,2]
  colnames(df_full_saliencies) <- GetSpeciesNames(df_abundance[start_row:nrow(df_abundance),1])
  dim_saliency_map <- dim(df_full_saliencies)
  print(paste("Dimension of saliency map:", dim_saliency_map[[1]], "x", dim_saliency_map[[2]]))
  
  fname_labels <- file.path(DIR_SALIENCY, dataset, "full_labels_.npy")
  df_full_labels <- as.data.frame(npyLoad(fname_labels))
  
  fname_predictions <- file.path(DIR_SALIENCY, dataset, "full_predictions_.npy")
  df_full_predictions <- as.data.frame(npyLoad(fname_predictions))

  # Convert predictions to 0/1 labels
  df_full_predictions_lb_ <- df_full_predictions
  df_full_predictions_lb_[] <- lapply(
    df_full_predictions_lb_, function(x) ifelse(x>class_one_threshold, 1, 0))
  
  # Get all true positive experiments

  tp_experiments_ind <- which(c(df_full_predictions_lb_)$V1 == c(df_full_labels)$V1, arr.ind = TRUE)
  df_tp_saliences <- df_full_saliencies[tp_experiments_ind,]
  
  # Just for check
  arr_pos_predictions <- df_full_predictions[tp_experiments_ind,]
  arr_pos_labels <- df_full_labels[tp_experiments_ind,]
  
  # Plot saliencies
  plots_tp_saliency <- PlotSaliencyMap(
    df_tp_saliences, df_stats, sort_mode = 'Mean', n_top = n_top_)
  
  # Combine all plots and save
  figure2 <- annotate_figure(
    ggarrange(
      plots_tp_saliency[[1]] + theme(
        plot.title = element_text(hjust = 0, face= "bold"), 
        plot.title.position = "plot",
        legend.position = "bottom", 
        plot.margin = margin(2, 1, 0, 1, unit = 'line')
      ), 
      ggarrange(
        plots_tp_saliency[[2]] + theme(
          plot.title = element_text(hjust = 0, face= "bold"), 
          plot.title.position = "plot",
          legend.position = "bottom", 
          plot.margin = margin(2, 1, 0, 1, unit = 'line')
        ), 
        plots_tp_saliency[[3]] + theme(
          plot.title = element_text(hjust = 0, face= "bold"), 
          plot.title.position = "plot",
          legend.position = "bottom", 
          plot.margin = margin(2, 1, 0, 1, unit = 'line')
        ),
        ncol = 2,
        legend = "none",
        labels = c("B.", "C.")
      ),
      nrow = 2,
      common.legend = TRUE, 
      legend = "bottom",
      labels = "A."
    ),
    top = text_grob(
      paste("Saliencies from TP maps for", dataset), 
      color = "black", face = "bold", size = 16)
  )
  #figure5
  fname <- file.path(DIR_PLOTS, dataset, paste("abundance_plots_TP_saliencies_no_rescale_all_pval-", p_thr_, ".png", sep = ""))
  ggsave(figure2, width = 20, height = 14, dpi = 300, bg = 'white', filename = fname)
  print(paste("Plots from TP saliency map saved into", fname))
  
  # Save plots separately
  
  figure2A <- annotate_figure(
    plots_tp_saliency[[1]],
    top = text_grob(
      paste("Saliencies from TP maps for", dataset), 
      color = "black", face = "bold", size = 16)
  )
  fname <- file.path(DIR_PLOTS, dataset, paste("abundance_plots_TP_saliencies_no_rescale_summaries_pval-", p_thr_, ".png", sep = ""))
  ggsave(figure2A, width = 12, height = 8, dpi = 300, bg = 'white', filename = fname)
  print(paste("Plot from summary statistics from TP saliency map saved into", fname))
  
  figure2B <- annotate_figure(
    plots_tp_saliency[[2]],
    top = text_grob(
      paste("Saliencies from TP maps for", dataset), 
      color = "black", face = "bold", size = 16)
  )
  fname <- file.path(DIR_PLOTS, dataset, paste("abundance_plots_TP_saliencies_no_rescale_boxplot_pval-", p_thr_, ".png", sep = ""))
  ggsave(figure2B, width = 12, height = 8, dpi = 300, bg = 'white', filename = fname)
  print(paste("Boxplot from TP saliency map saved into", fname))
  
  figure2C <- annotate_figure(
    plots_tp_saliency[[3]],
    top = text_grob(
      paste("Saliencies from TP maps for", dataset), 
      color = "black", face = "bold", size = 16)
  )
  fname <- file.path(DIR_PLOTS, dataset, paste("abundance_plots_TP_saliencies_no_rescale_violine_pval-", p_thr_, ".png", sep = ""))
  ggsave(figure2C, width = 12, height = 8, dpi = 300, bg = 'white', filename = fname)
  print(paste("Violine plot from TP saliency map saved into", fname))
}


# -------------------------------------------------------------------------
# (3) Barplots of mean saliences from true positives (full maps, no rescaling)
#
# Abundances

fname <- file.path(DIR_SALIENCY, dataset, "full_abundance_saliency_.npy")
if (file.exists(fname)) {
  print("Generate plots from full saliency maps.")
  
  df_full_saliencies <- as.data.frame(npyLoad(fname))
  #colnames(df_full_saliencies)<- str_split_fixed(df_abundance[start_row:nrow(df_abundance),1], ".s__", 2)[,2]
  colnames(df_full_saliencies) <- GetSpeciesNames(df_abundance[start_row:nrow(df_abundance),1])
  dim_saliency_map <- dim(df_full_saliencies)
  print(paste("Dimension of saliency map:", dim_saliency_map[[1]], "x", dim_saliency_map[[2]]))
  
  fname_labels <- file.path(DIR_SALIENCY, dataset, "full_labels_.npy")
  df_full_labels <- as.data.frame(npyLoad(fname_labels))
  
  fname_predictions <- file.path(DIR_SALIENCY, dataset, "full_predictions_.npy")
  df_full_predictions <- as.data.frame(npyLoad(fname_predictions))
  
  # Convert predictions to 0/1 labels
  df_full_predictions_lb_ <- df_full_predictions
  df_full_predictions_lb_[] <- lapply(
    df_full_predictions_lb_, function(x) ifelse(x>class_one_threshold, 1, 0))
  
  # Get all true positive experiments
  tp_experiments_ind <- which(c(df_full_predictions_lb_)$V1 == c(df_full_labels)$V1, arr.ind = TRUE)
  df_tp_saliences <- df_full_saliencies[tp_experiments_ind,]

  # Calculate summary statistics
  df_melt <- melt(df_tp_saliences, id.vars = NULL)

  df_mean_summr <- aggregate(df_melt[, 2], list(df_melt$variable), mean)
  df_se_summr <- aggregate(df_melt[, 2], list(df_melt$variable), function(x) sqrt(var(x)/length(x)))
  
  df_mean_summr <- merge(df_mean_summr, df_se_summr, by.x = 1, by.y = 1, all.x = TRUE, all.y = FALSE) 
  df_summr <- merge(df_mean_summr, df_stats['representation'], by.x = 1, by.y = 0, all.x = TRUE, all.y = FALSE)
  
  colnames(df_summr) <- c('species', 'mean', 'se', 'representation')
  
  # Save to CSV ordered list of species
  write.table(df_summr[order(df_summr$mean, decreasing = TRUE), ], 
            file.path(DIR_PLOTS, dataset, paste("abundance_TP_saliencies_no_rescale_pval-", p_thr_, ".csv", sep = "")), 
            row.names=FALSE,
            sep = "\t",
            quote = FALSE)
  
  df_summr <- df_summr[order(df_summr$mean, decreasing = TRUE), ][1:n_top_,]
  
  # Draw barplot
  p_barplot_err <- ggplot(
    df_summr,
    aes(x = reorder(species, mean), y = mean, fill = representation)
  ) +
    geom_col(aes(), width = 0.7) + 
    geom_errorbar(
      aes(ymin = mean - se, ymax = mean + se), width = .2, position = position_dodge(.9)
    ) +
    scale_fill_manual("", values = c("nd" = "grey", "prevail in affected" = "red", "prevail in healthy" = "blue")) +
    labs(
      title = paste("Mean saliencies of abundances for", dataset, "(true positives)"), 
      x = paste("Top", n_top_, "species by mean saliency"), 
      y = "Saliency"
    ) +
    theme(
      plot.title = element_text(hjust = 0.5, face= "bold"), 
      plot.title.position = "plot",
      legend.position = "bottom", 
      plot.margin = margin(2, 1, 0, 1, unit = 'line')
    ) + 
    coord_flip()
  p_barplot_err
    
  fname <- file.path(DIR_PLOTS, dataset, paste("abundance_errbarplot_TP_saliencies_no_rescale_pval-", p_thr_, ".png", sep = ""))
  ggsave(p_barplot_err, width = 12, height = 8, dpi = 300, bg = 'white', filename = fname)
  print(paste("Barplots from TP saliency map saved into", fname))
  
}


# Markers plots ----------------------------------------------------------------
# (4) Error barplots, boxplot and violine plot for markers 
# from full saliency maps (true positives, no rescale)

fname_markers <- file.path(DIR_DATA, "marker", paste("marker_", dataset, ".txt", sep = ""))
fname <- file.path(DIR_SALIENCY, dataset, "full_markers_saliency_.npy")
if (file.exists(fname_markers) & file.exists(fname)) {
  print("Generate marker plots from full saliency maps.")
  
  df_markers <- read.csv(fname_markers, sep="\t")
  df_full_saliencies <- as.data.frame(npyLoad(fname))
  colnames(df_full_saliencies) <- df_markers[start_row:nrow(df_markers),1]
  dim_saliency_map <- dim(df_full_saliencies)
  print(paste("Dimension of saliency map:", dim_saliency_map[[1]], "x", dim_saliency_map[[2]]))
  
  fname_labels <- file.path(DIR_SALIENCY, dataset, "full_labels_.npy")
  df_full_labels <- as.data.frame(npyLoad(fname_labels))
  
  fname_predictions <- file.path(DIR_SALIENCY, dataset, "full_predictions_.npy")
  df_full_predictions <- as.data.frame(npyLoad(fname_predictions))
  
  # Convert predictions to 0/1 labels
  df_full_predictions_lb_ <- df_full_predictions
  df_full_predictions_lb_[] <- lapply(
    df_full_predictions_lb_, function(x) ifelse(x>class_one_threshold, 1, 0))
  
  # Get all true positive experiments
  tp_experiments_ind <- which(c(df_full_predictions_lb_)$V1 == c(df_full_labels)$V1, arr.ind = TRUE)
  df_tp_saliences <- df_full_saliencies[tp_experiments_ind,]
  
  # Select top markers
  df_saliencies_summary <- as.data.frame(t(do.call(cbind, lapply(df_tp_saliences, summary))))
  df_saliencies_summary <- df_saliencies_summary[order(df_saliencies_summary[,'Mean'], decreasing = TRUE),]
  
  # Save into CSV ordered list of markers
  write.table(df_saliencies_summary['Mean'], 
              file.path(DIR_PLOTS, dataset, paste("marker_TP_saliencies_no_rescale.csv", sep = "")), 
              row.names=TRUE,
              sep = "\t",
              quote = FALSE, 
              col.names = NA)
  
  # Select top species with highest mean saliencies
  top_species <- row.names(df_saliencies_summary[1:n_top_,])
  
  # Calculate summary statistics
  df_melt <- melt(df_tp_saliences[,top_species], id.vars = NULL)
  
  df_mean_summr <- aggregate(df_melt[, 2], list(df_melt$variable), mean)
  df_se_summr <- aggregate(df_melt[, 2], list(df_melt$variable), function(x) sqrt(var(x)/length(x)))
  
  df_summr <- merge(df_mean_summr, df_se_summr, by.x = 1, by.y = 1, all.x = TRUE, all.y = FALSE)
  colnames(df_summr) <- c('species', 'mean', 'se')
  
  # Draw barplot
  p_barplot_err <- ggplot(
    df_summr,
    aes(x = reorder(species, mean), y = mean, fill = mean)
  ) +
    geom_col(aes(), width = 0.7) + 
    geom_errorbar(
      aes(ymin = mean - se, ymax = mean + se), width = .2, position = position_dodge(.9)
    ) +
    labs(
      title = paste("Mean saliencies of markers for", dataset, "(true positives)"), 
      x = paste("Top", n_top_, "markers by mean saliency"), 
      y = "Saliency"
    ) +
    theme(
      plot.title = element_text(hjust = 0.5, face= "bold"), 
      plot.title.position = "plot",
      legend.position = "none", 
      plot.margin = margin(2, 1, 0, 1, unit = 'line')
    ) + 
    coord_flip()
  p_barplot_err  
  
  fname <- file.path(DIR_PLOTS, dataset, paste("markers_errbarplot_TP_saliencies_no_rescale.png", sep = ""))
  ggsave(p_barplot_err, width = 12, height = 8, dpi = 300, bg = 'white', filename = fname)
  print(paste("Barplots plot from TP markers saliency map saved into", fname))
  
  # Other plots
  rescale_plot <- TRUE
  
  # Draw boxplot
  p_boxplot <- ggplot(
    df_melt, 
    aes(x = reorder(variable, value, mean), y = value, fill = value)
  ) + 
    geom_boxplot(outlier.shape = NA) + 
    stat_summary(
      fun = function(x) 
        ifelse(rescale_plot == TRUE, log10(mean(10^x)), mean(x)), 
      geom = "point", shape = 5, colour = "black"
    ) +
    labs(
      title = "", 
      x = paste("Top", n_top_, "markers by", tolower(sort_mode_), "saliency"), 
      y = "Saliency"
    ) +
    scale_y_continuous(trans=SymLogRescale()) + 
    coord_flip()
  #p_boxplot
  
  figure4A <- annotate_figure(
    p_boxplot,
    top = text_grob(
      paste("Marker saliencies from TP maps for", dataset), 
      color = "black", face = "bold", size = 16)
  )
  fname <- file.path(DIR_PLOTS, dataset, paste("marker_plots_TP_saliencies_no_rescale_boxplot.png", sep = ""))
  ggsave(figure4A, width = 12, height = 8, dpi = 300, bg = 'white', filename = fname)
  print(paste("Boxplot from TP saliency map saved into", fname))
  
  # Draw violine plot
  p_violine <- ggplot(
    df_melt, 
    aes(x = reorder(variable, value, mean), y = value, fill = value)
  ) + 
    geom_violin(draw_quantiles = c(0.25, 0.5, 0.75),) + 
    stat_summary(
      fun = function(x) 
        ifelse(rescale_plot == TRUE, log10(mean(10^x)), mean(x)), 
      geom = "point", shape = 5, colour = "black"
    ) +
    labs(
      title = "", 
      x = paste("Top", n_top_, "markers by", tolower(sort_mode_), "saliency"), 
      y = "Saliency"
    ) +
    scale_y_continuous(trans=SymLogRescale()) + 
    coord_flip()
  #p_violine
  
  figure4B <- annotate_figure(
    p_violine,
    top = text_grob(
      paste("Marker saliencies from TP maps for", dataset), 
      color = "black", face = "bold", size = 16)
  )
  fname <- file.path(DIR_PLOTS, dataset, paste("marker_plots_TP_saliencies_no_rescale_violine.png", sep = ""))
  ggsave(figure4B, width = 12, height = 8, dpi = 300, bg = 'white', filename = fname)
  print(paste("Violine from TP saliency map saved into", fname))
  
}


  
```
