## Supplementary material for "Microbiome-based disease prediction with multimodal variational information bottlenecks": S8 File: README.txt

This folder contains for each dataset:

1) plots of saliency maps for top 25 microbial species
2) plots of saliency maps for top 25 strain markers

Additionally, you can find in this folder:

- the .R script used to generate the plots, in case you will need to make plots again with other parameters.

- the .ipynb notebook to convert the .npy array saved by the microbiome_mvib.saliency.Saliency class in a format which works well with .R
Remark: microbiome_mvib.saliency.Saliency saves on disk `full_prediction.npy`, `full_markers_saliency.npy`, `full_labels.npy`, `full_abundance_saliency.npy`
