## Supplementary figures and images for "Microbiome-based disease prediction with multimodal variational information bottlenecks"

### abundance_barplot_TP_saliencies_no_rescale_pval-0.1.png

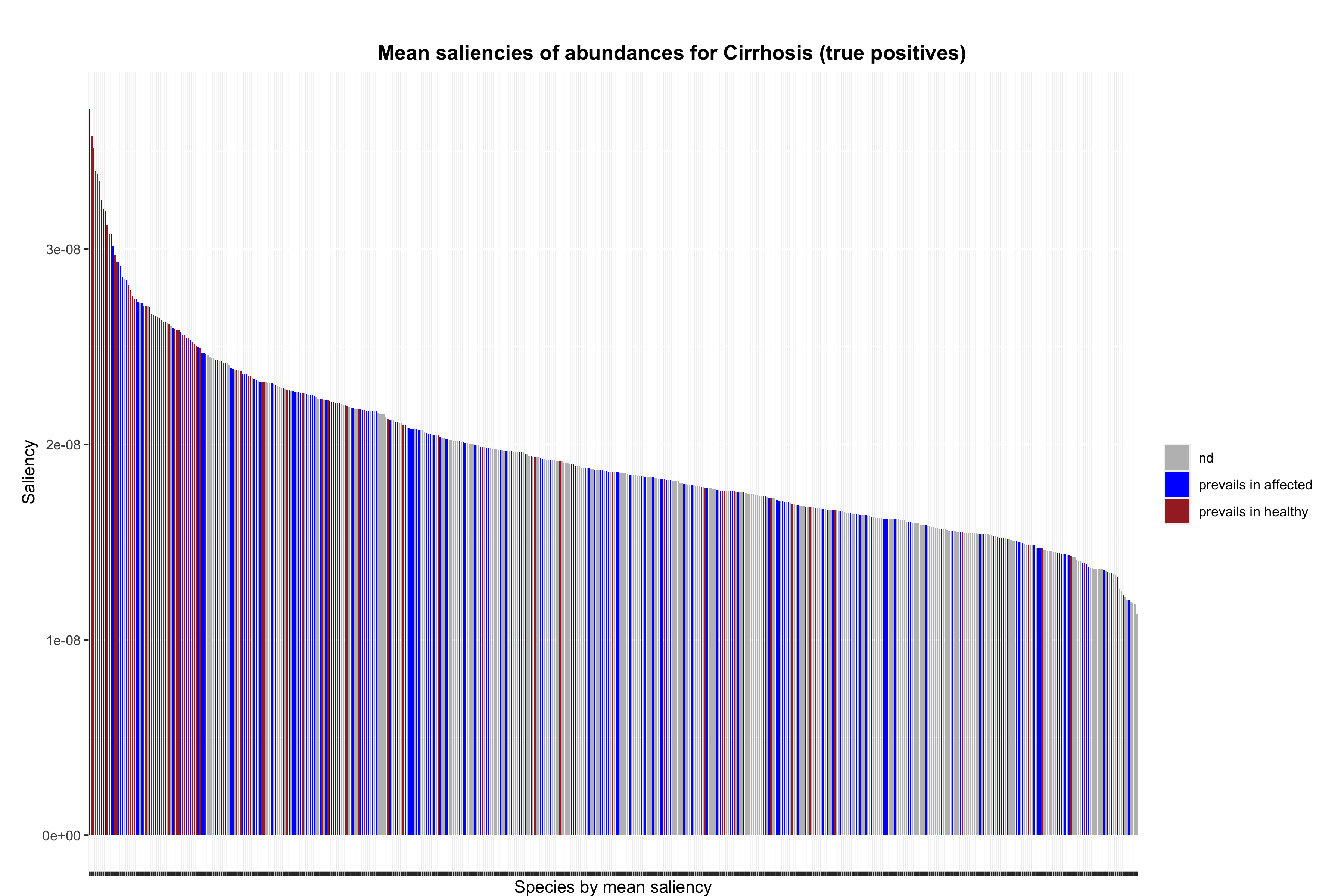

### abundance_barplot_TP_saliencies_no_rescale_pval-0.1.png

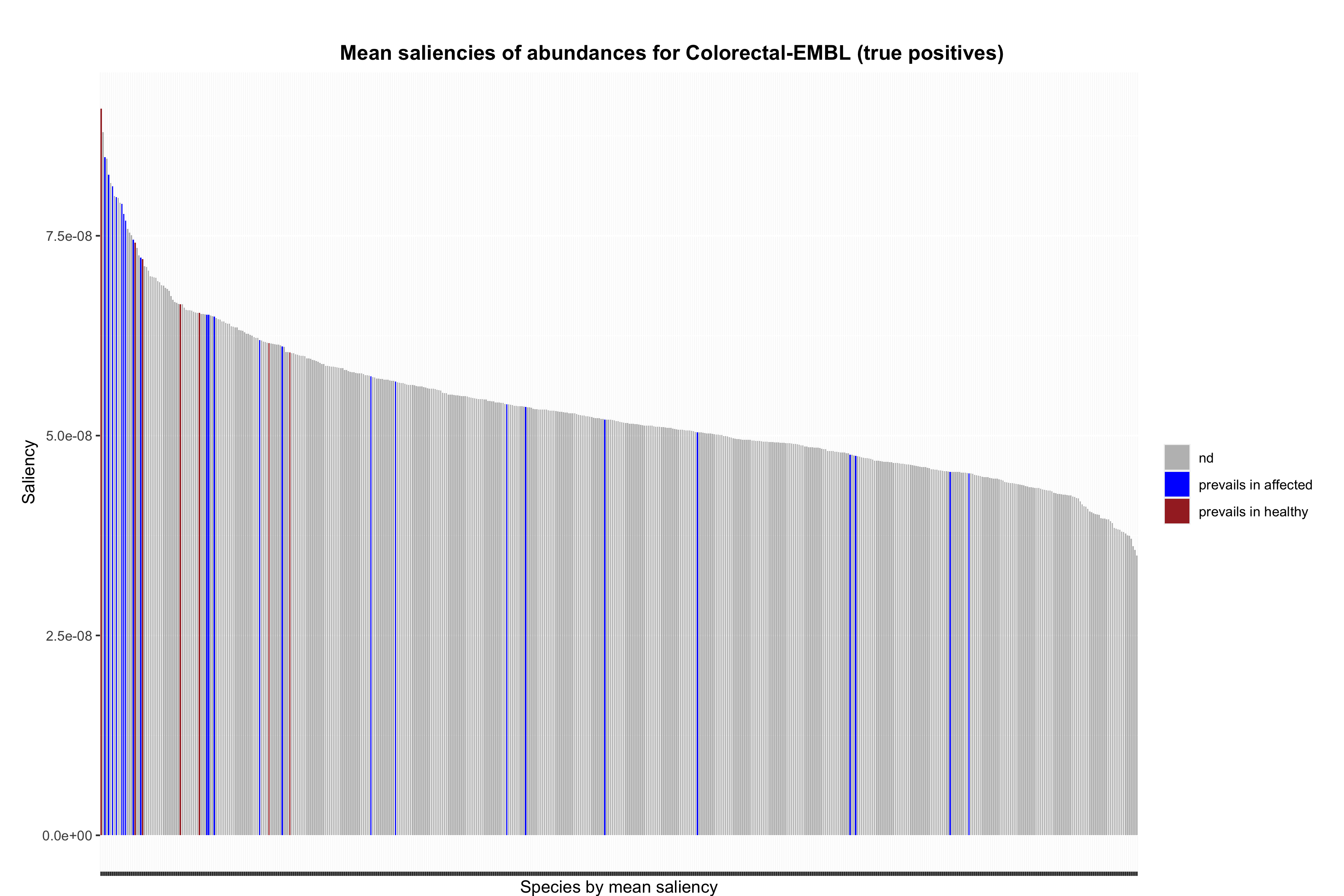

### abundance_barplot_TP_saliencies_no_rescale_pval-0.1.png

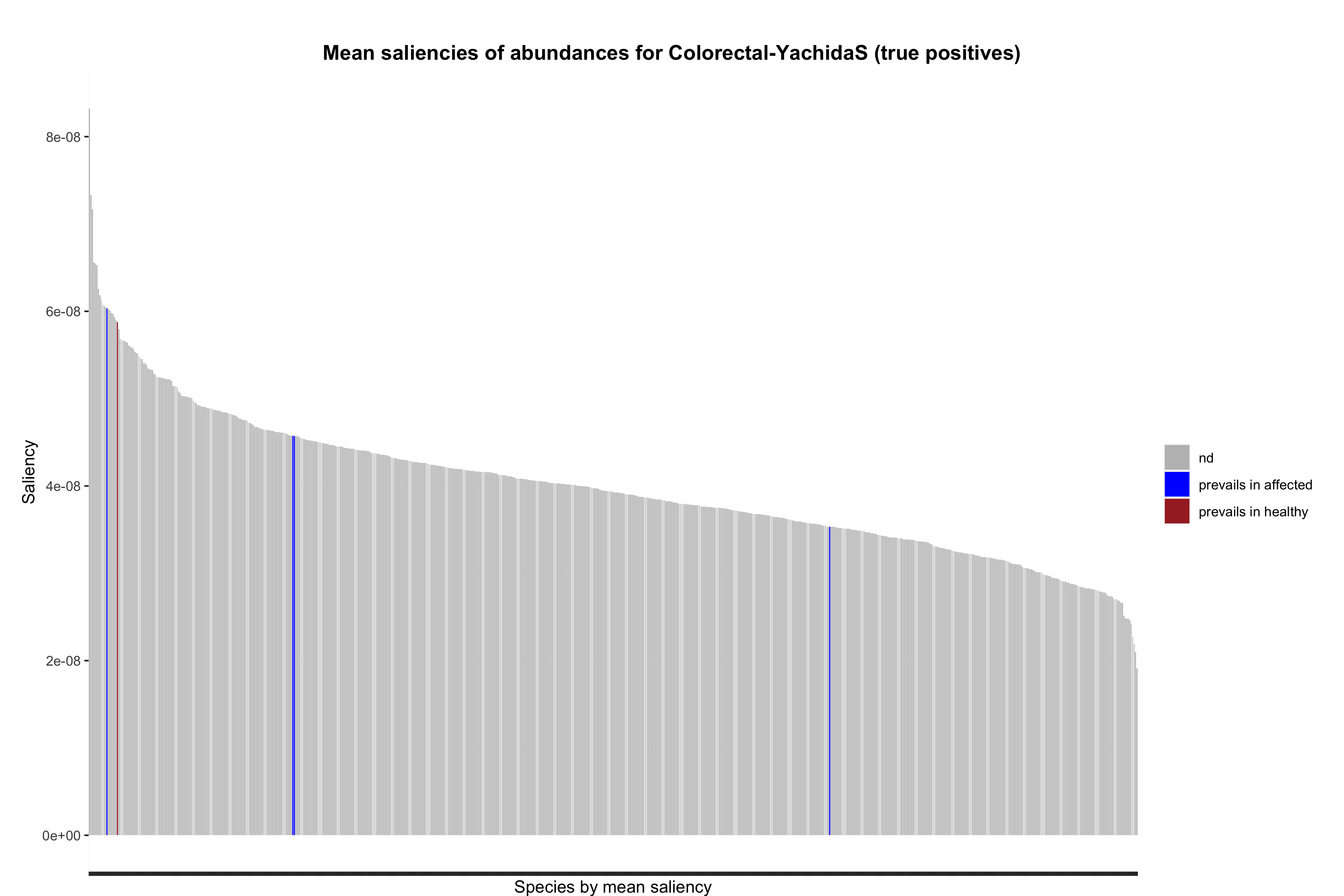

### abundance_barplot_TP_saliencies_no_rescale_pval-0.1.png

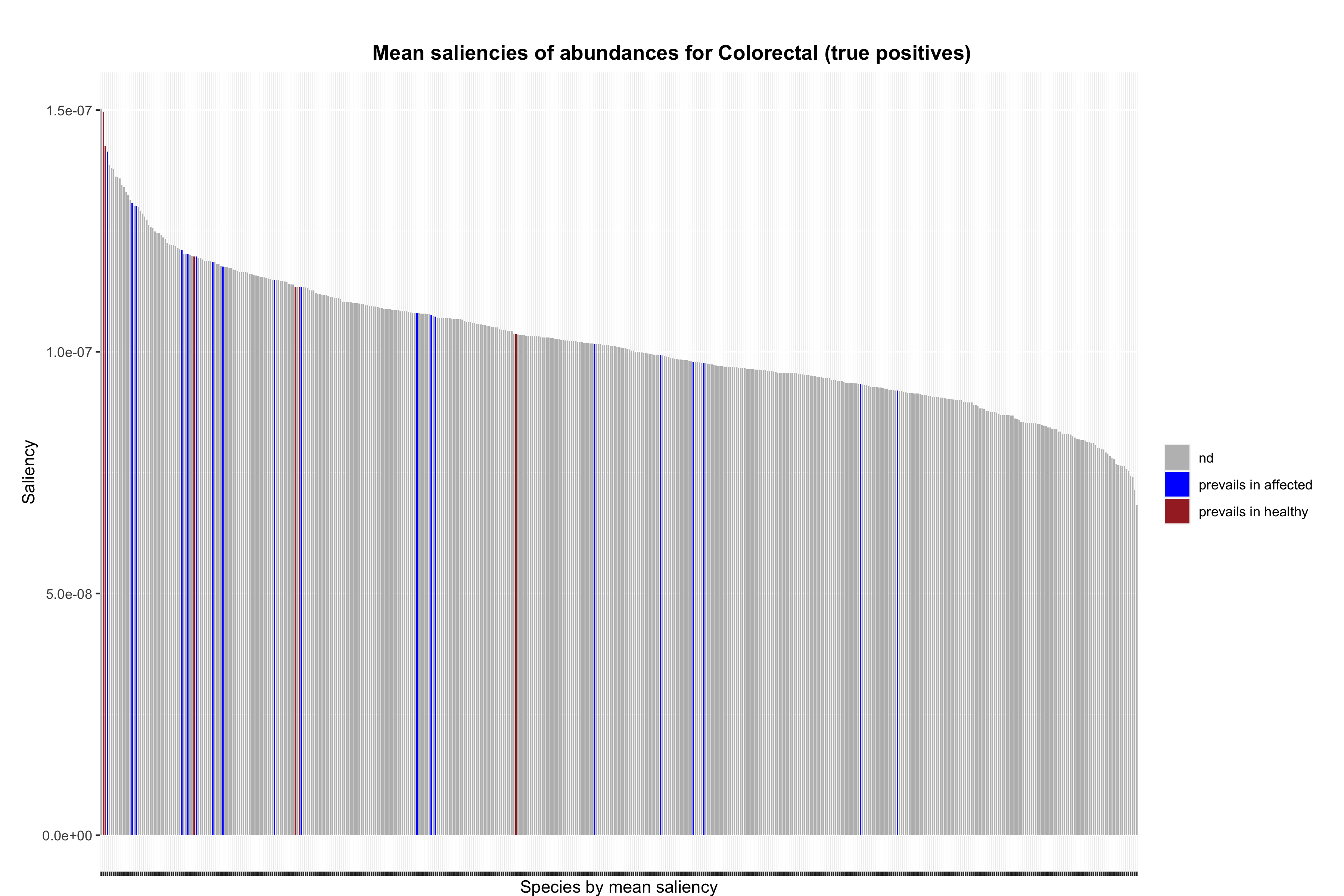

### abundance_barplot_TP_saliencies_no_rescale_pval-0.1.png

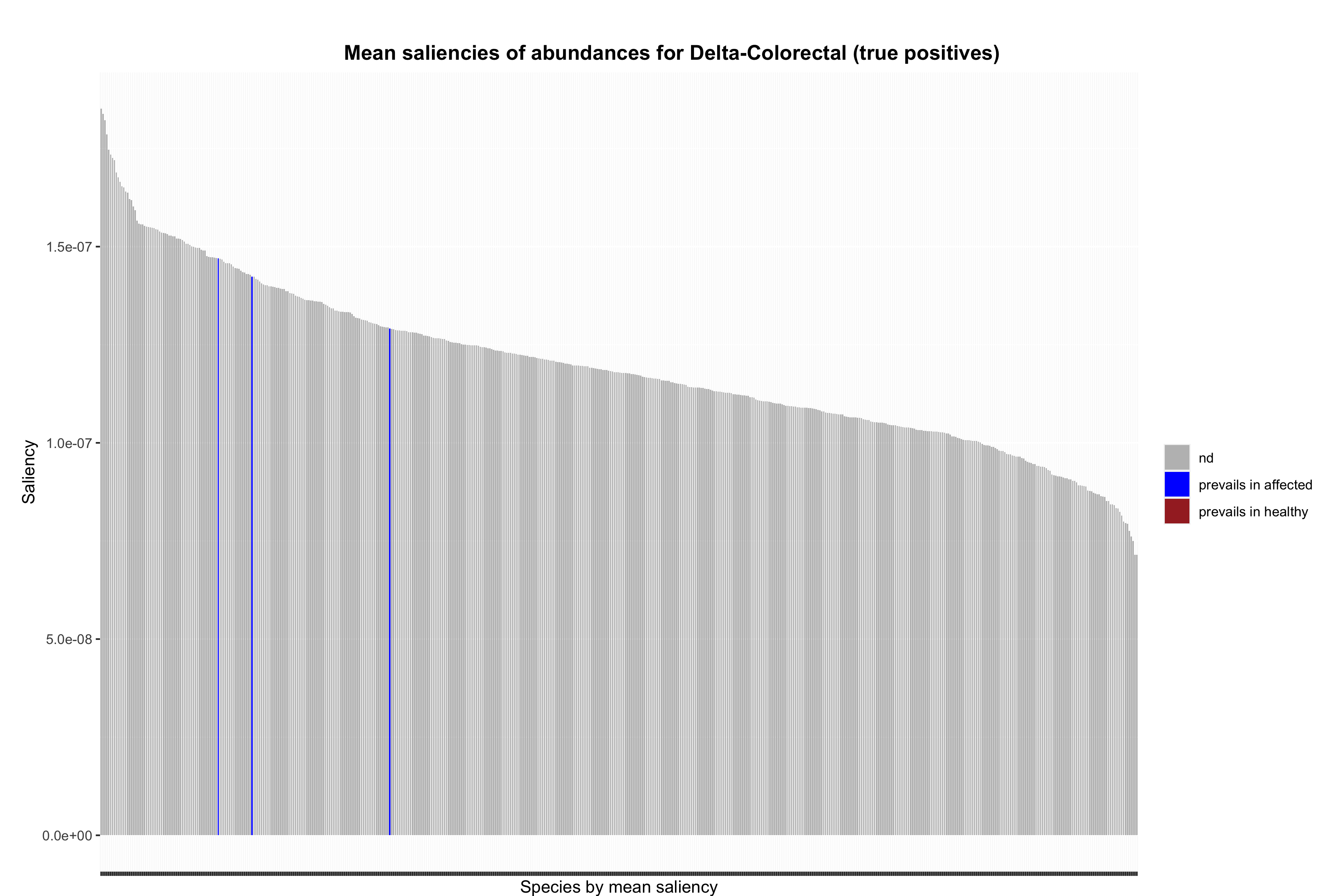

### abundance_barplot_TP_saliencies_no_rescale_pval-0.1.png

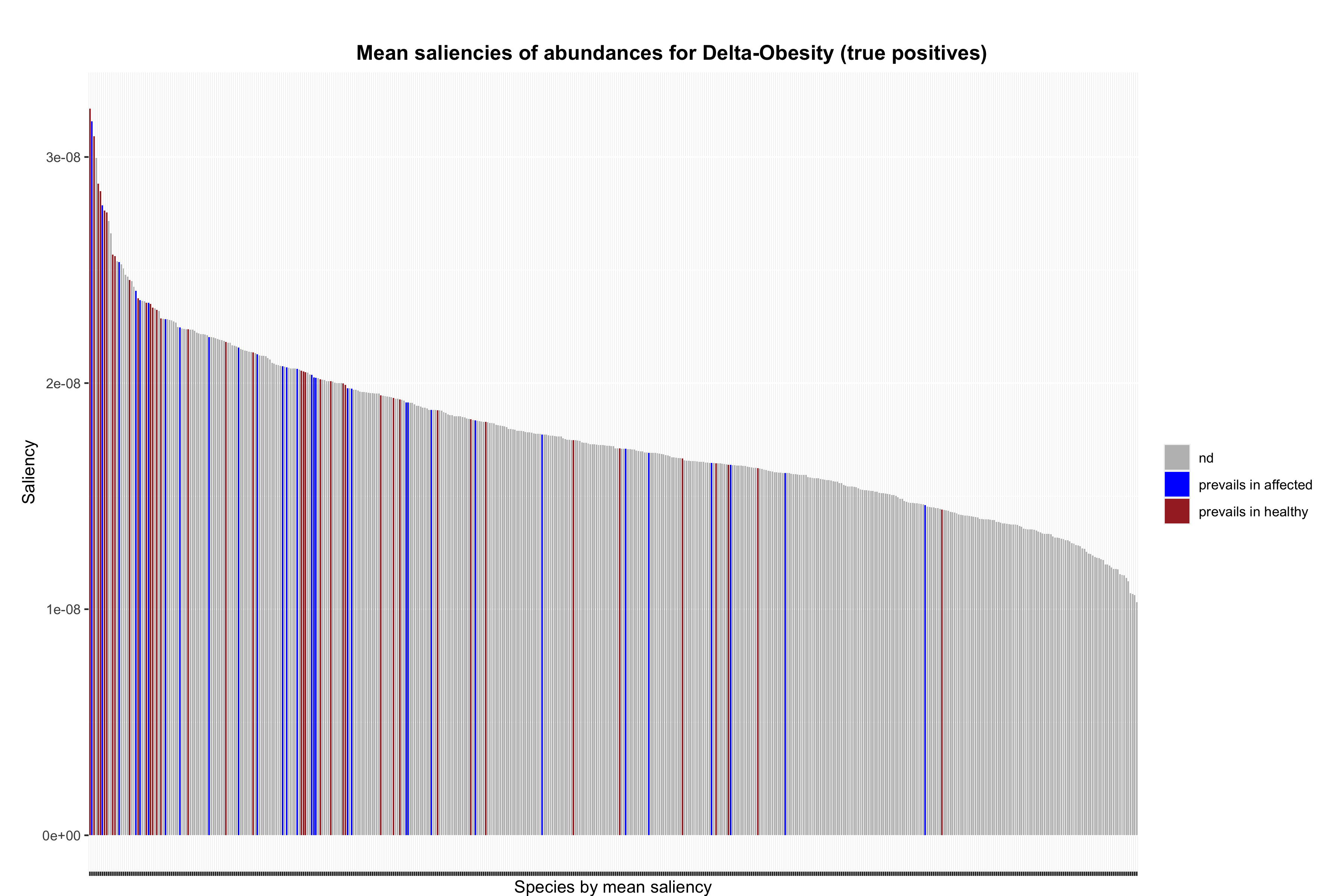

### abundance_barplot_TP_saliencies_no_rescale_pval-0.1.png

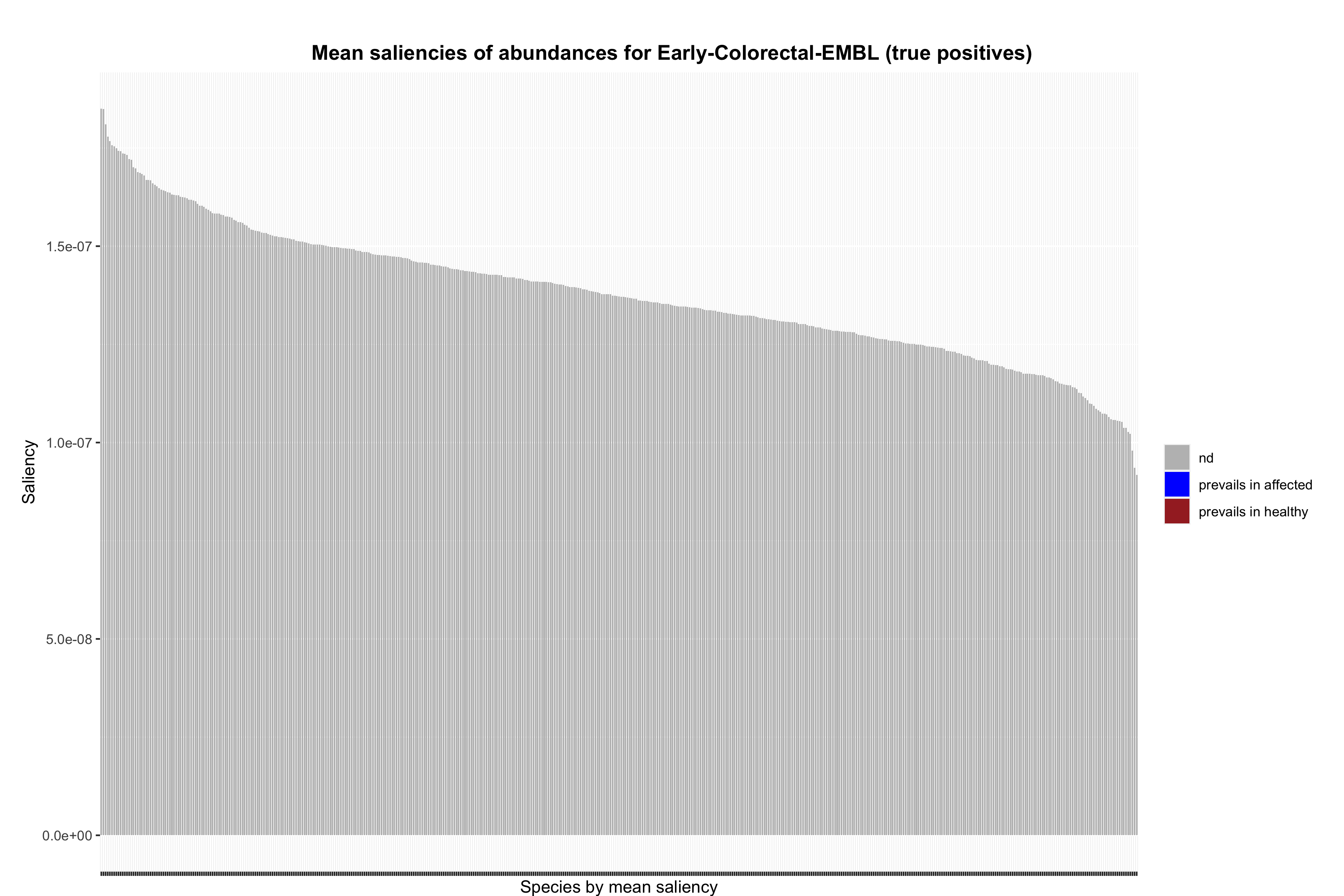

### abundance_barplot_TP_saliencies_no_rescale_pval-0.1.png

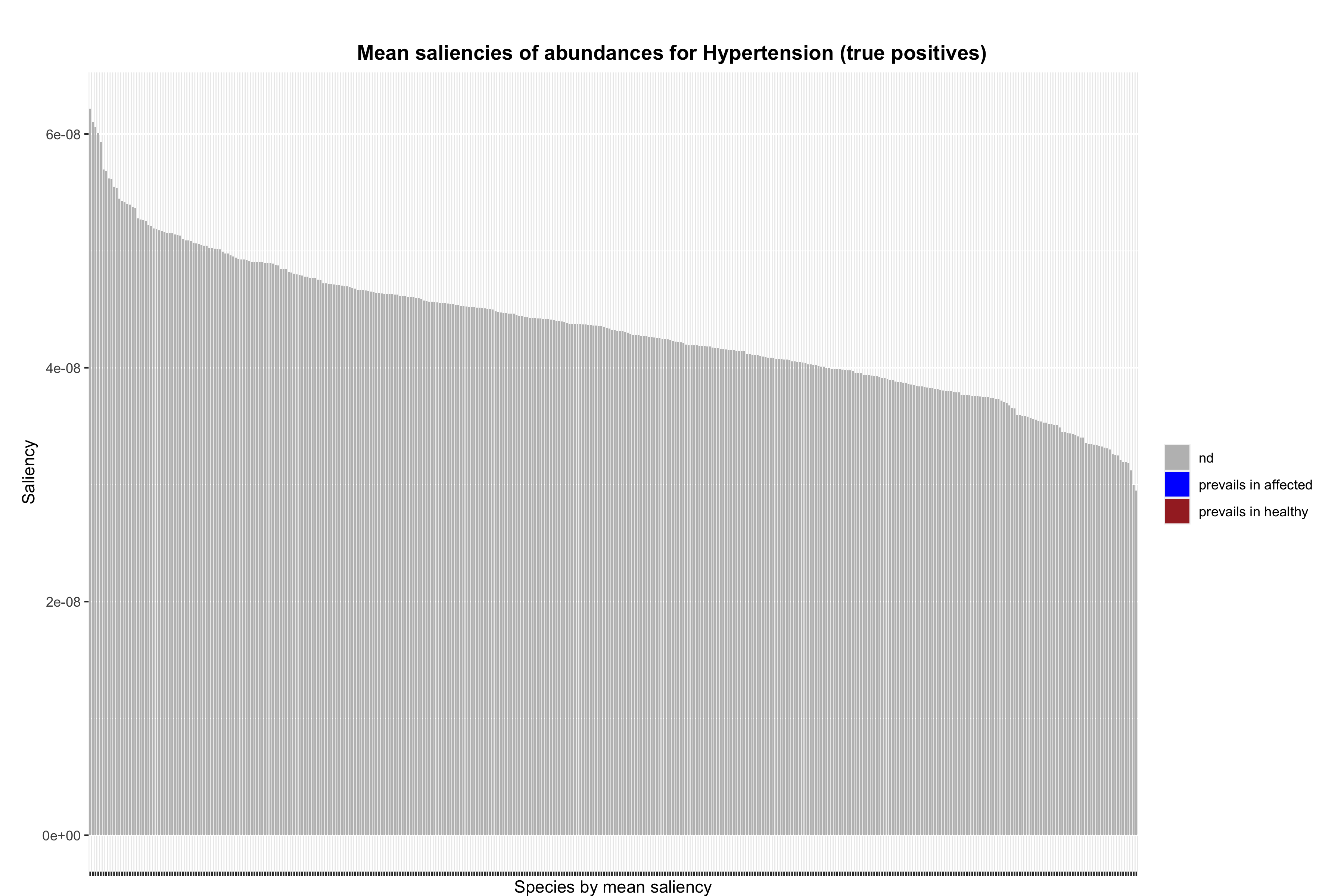

### abundance_barplot_TP_saliencies_no_rescale_pval-0.1.png

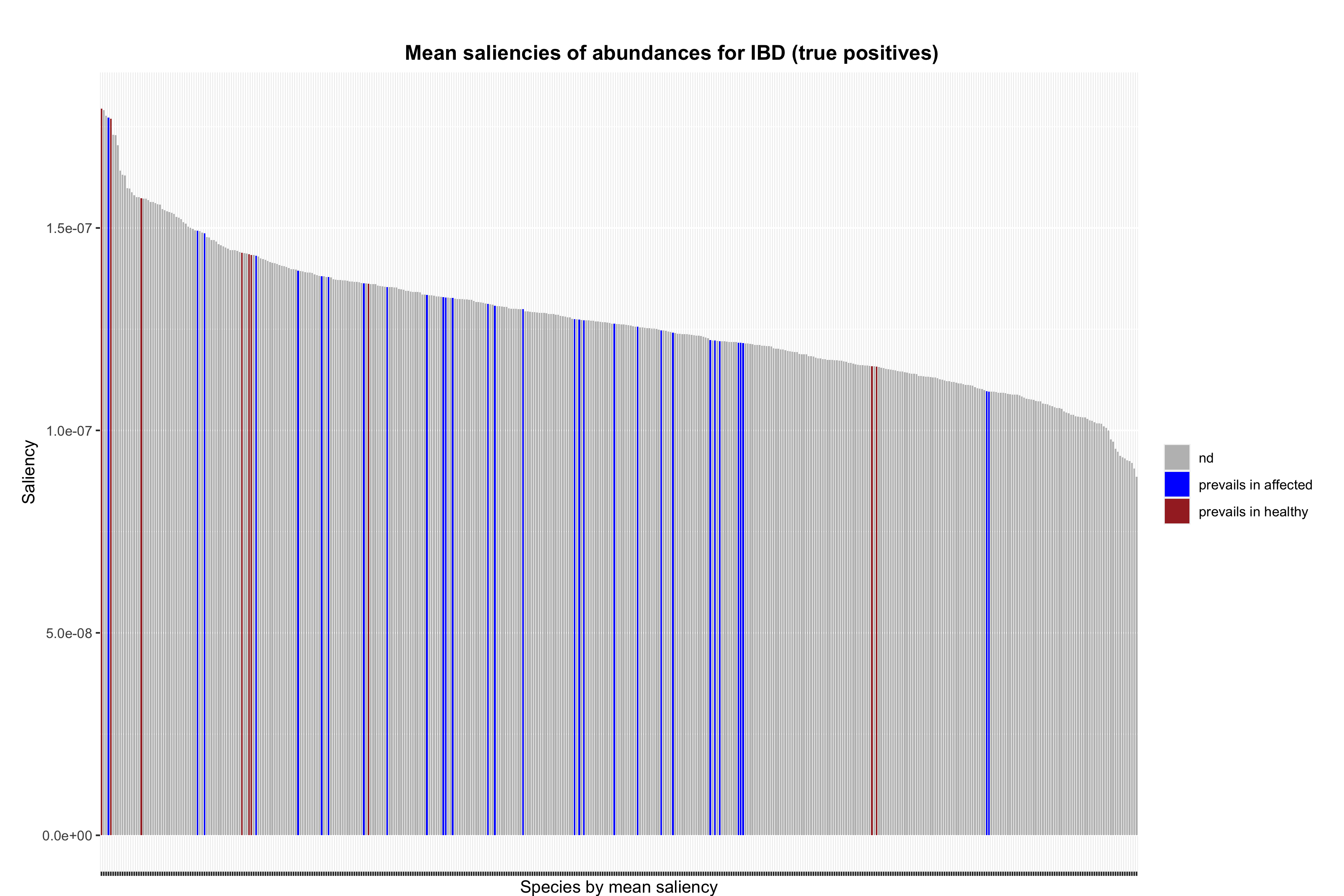

### abundance_barplot_TP_saliencies_no_rescale_pval-0.1.png

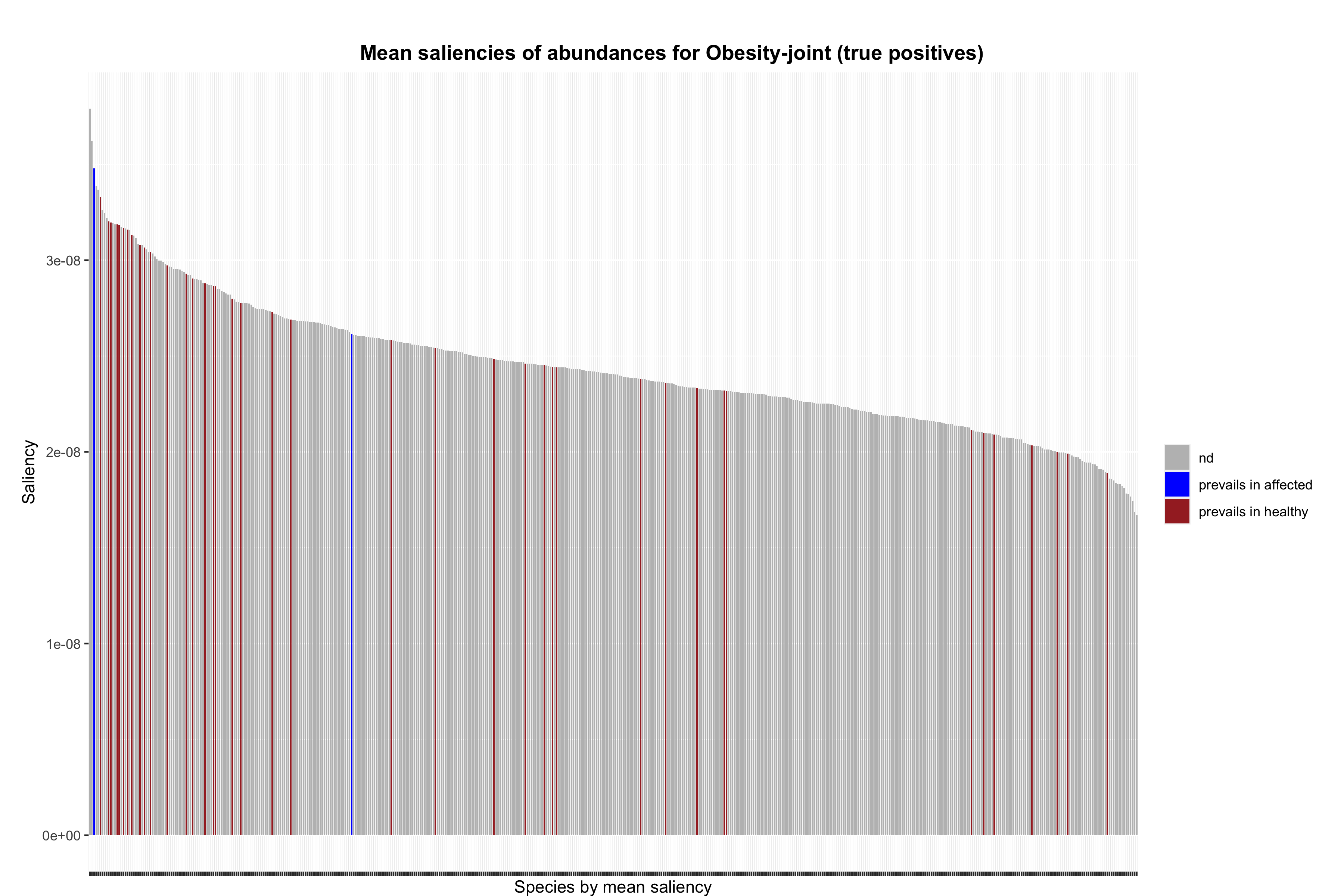

### abundance_barplot_TP_saliencies_no_rescale_pval-0.1.png

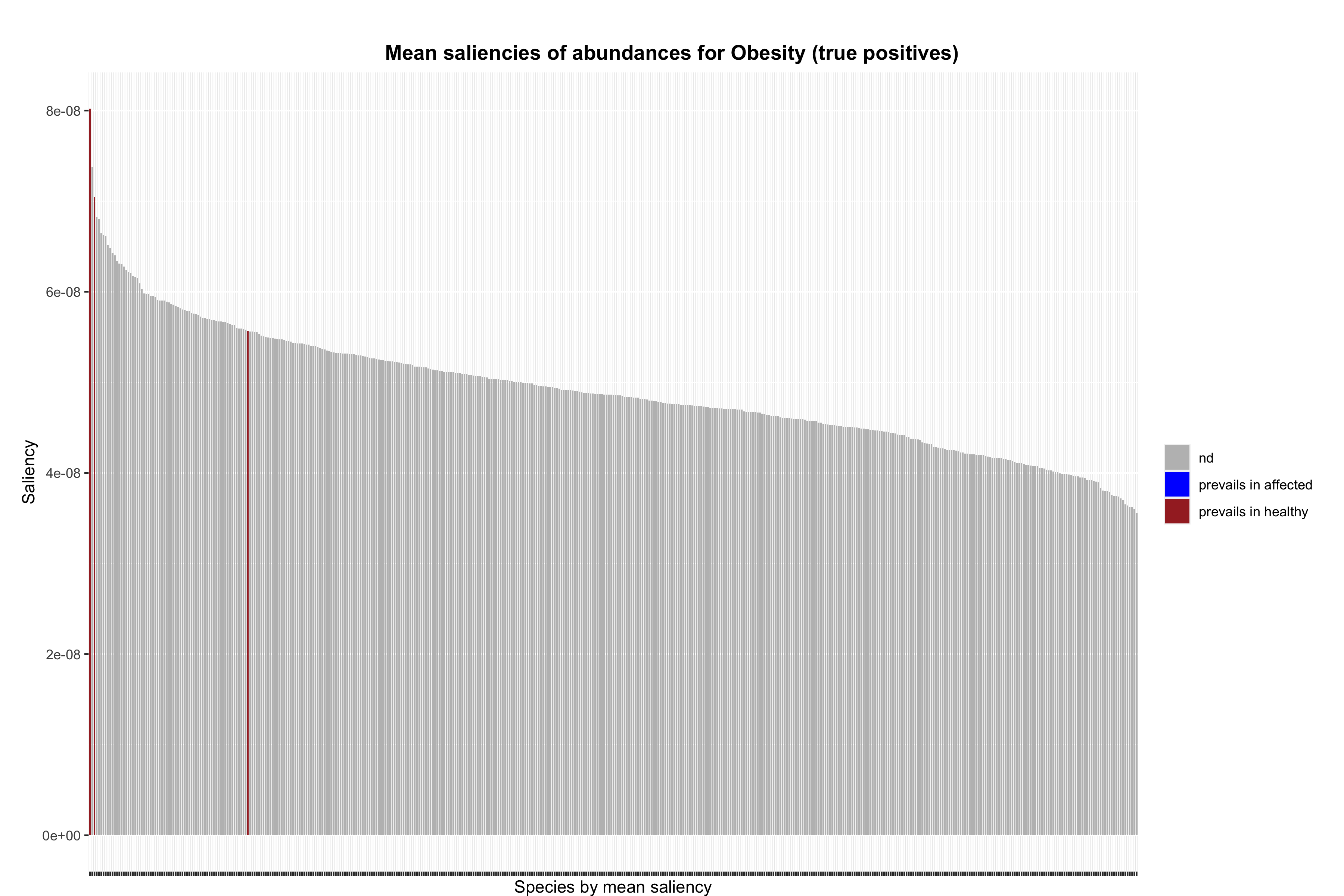

### abundance_barplot_TP_saliencies_no_rescale_pval-0.1.png

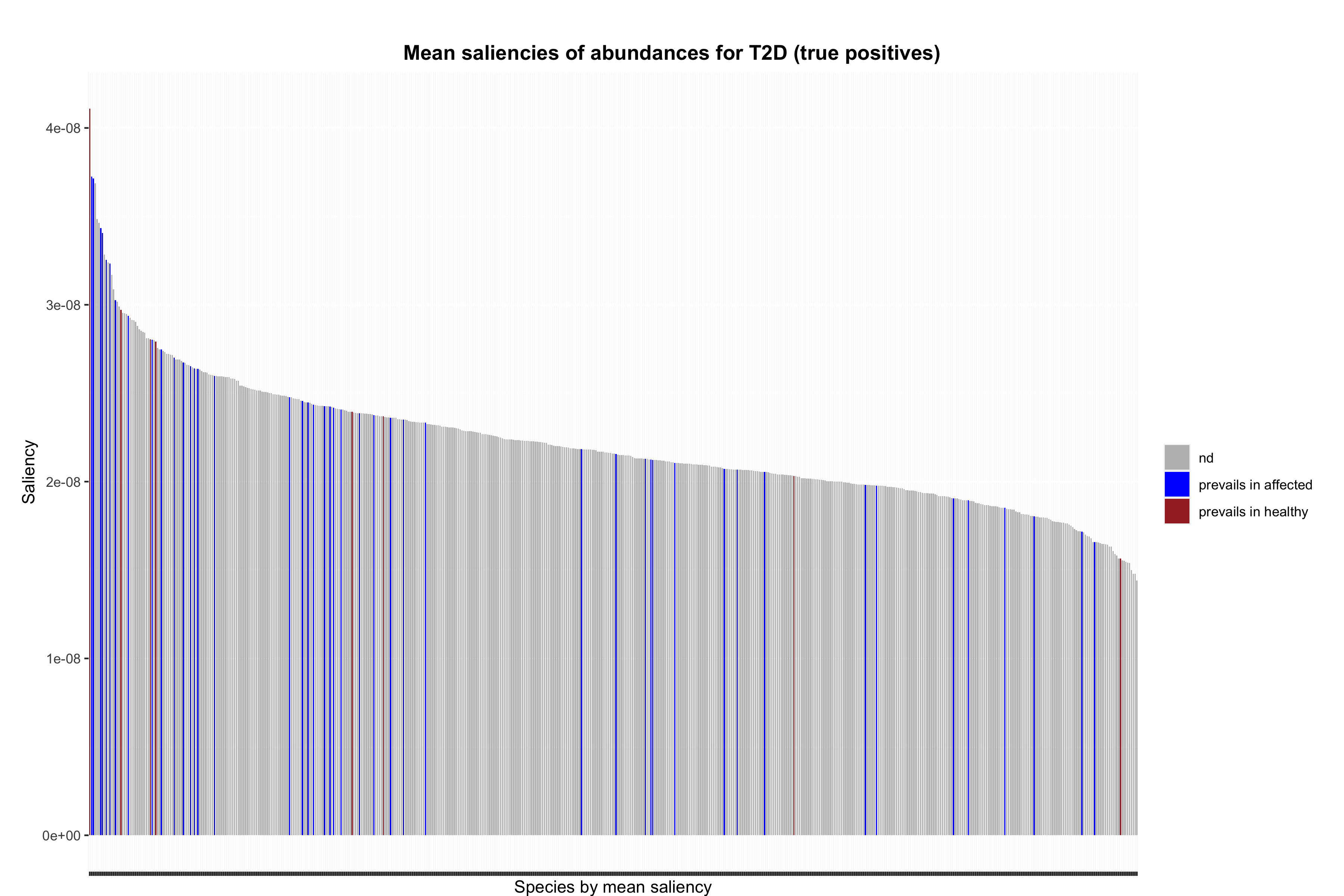

### abundance_barplot_TP_saliencies_no_rescale_pval-0.1.png

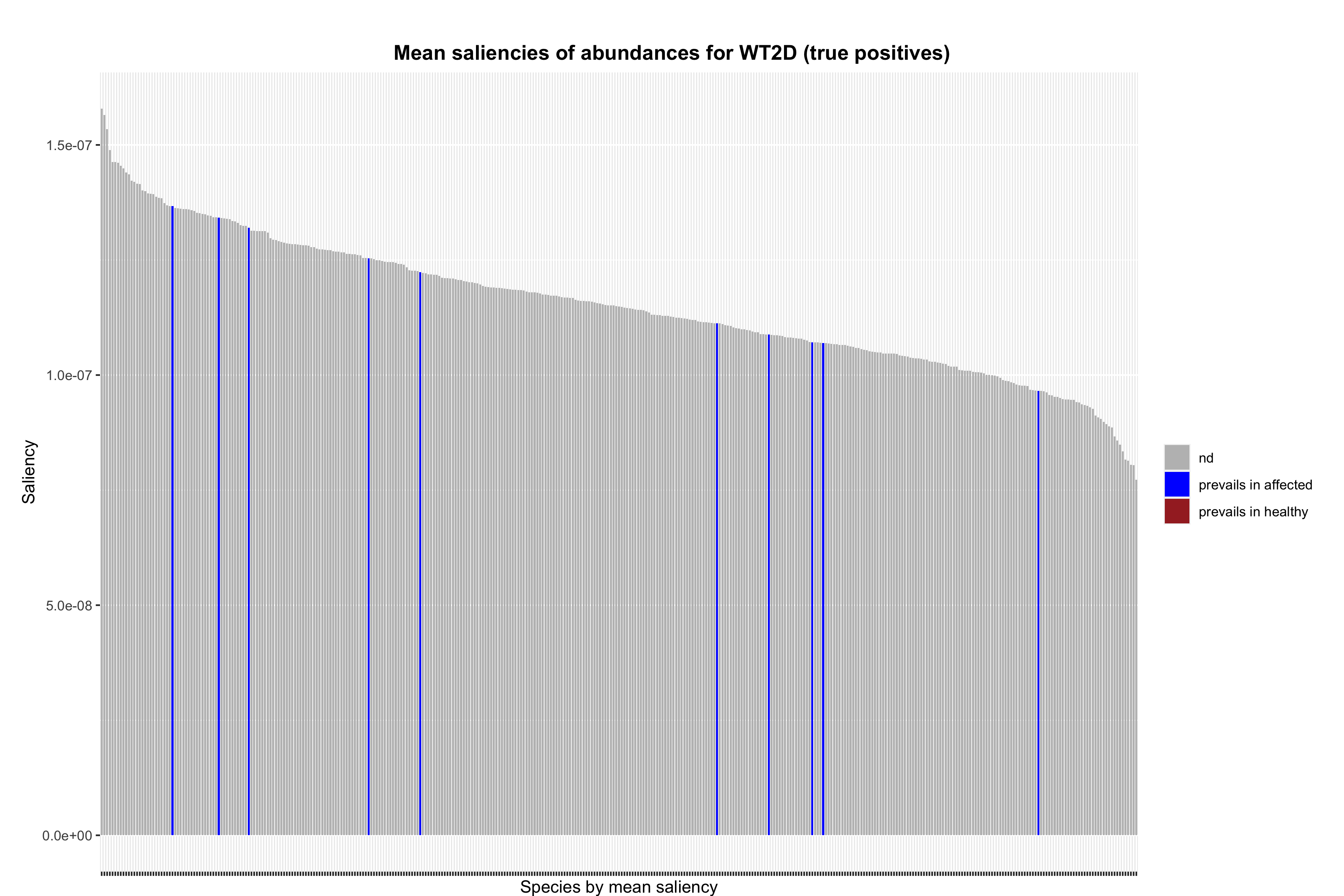

### abundance_errbarplot_TP_saliencies_no_rescale_pval-0.1.png

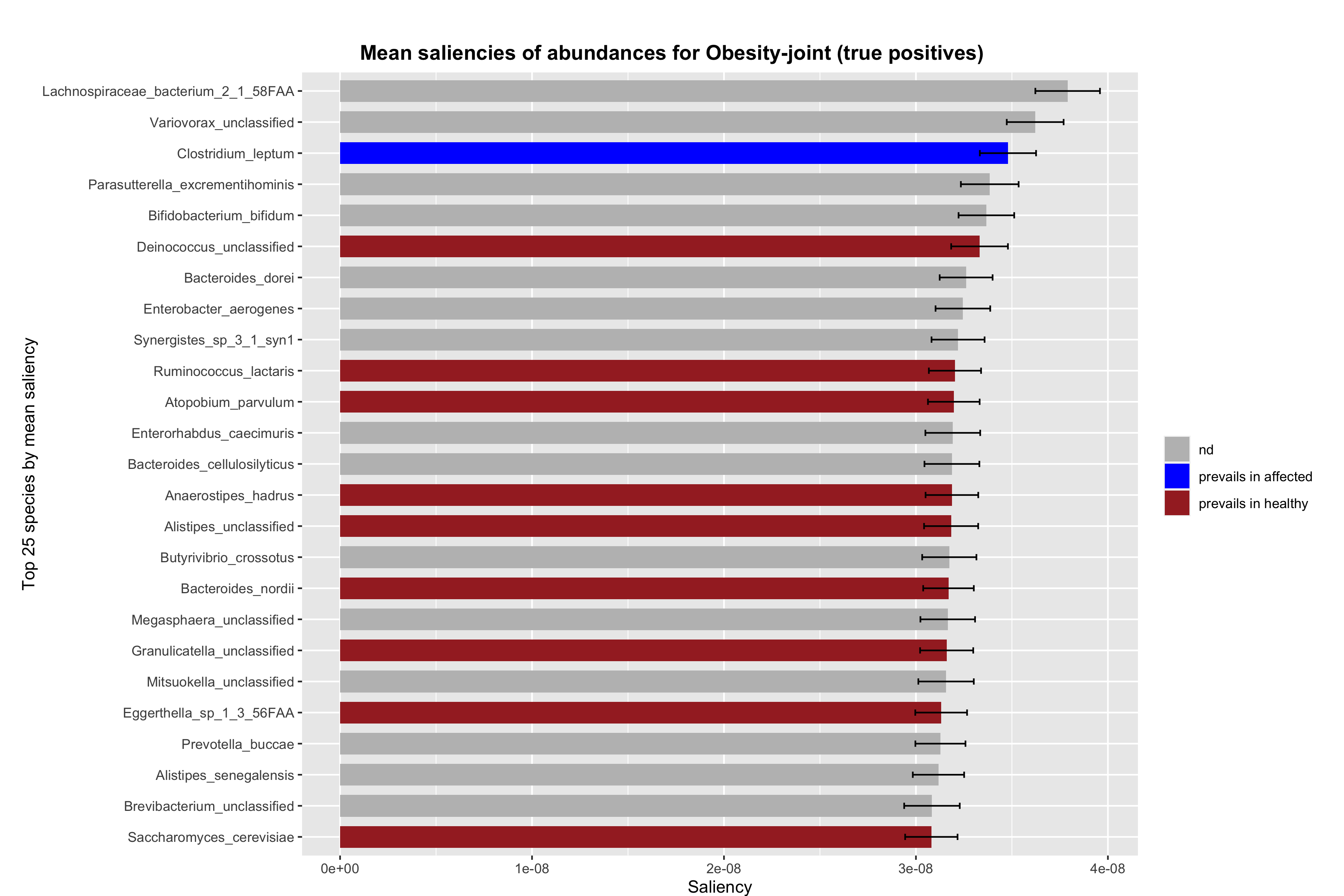

### abundance_errbarplot_TP_saliencies_no_rescale_pval-0.1.png

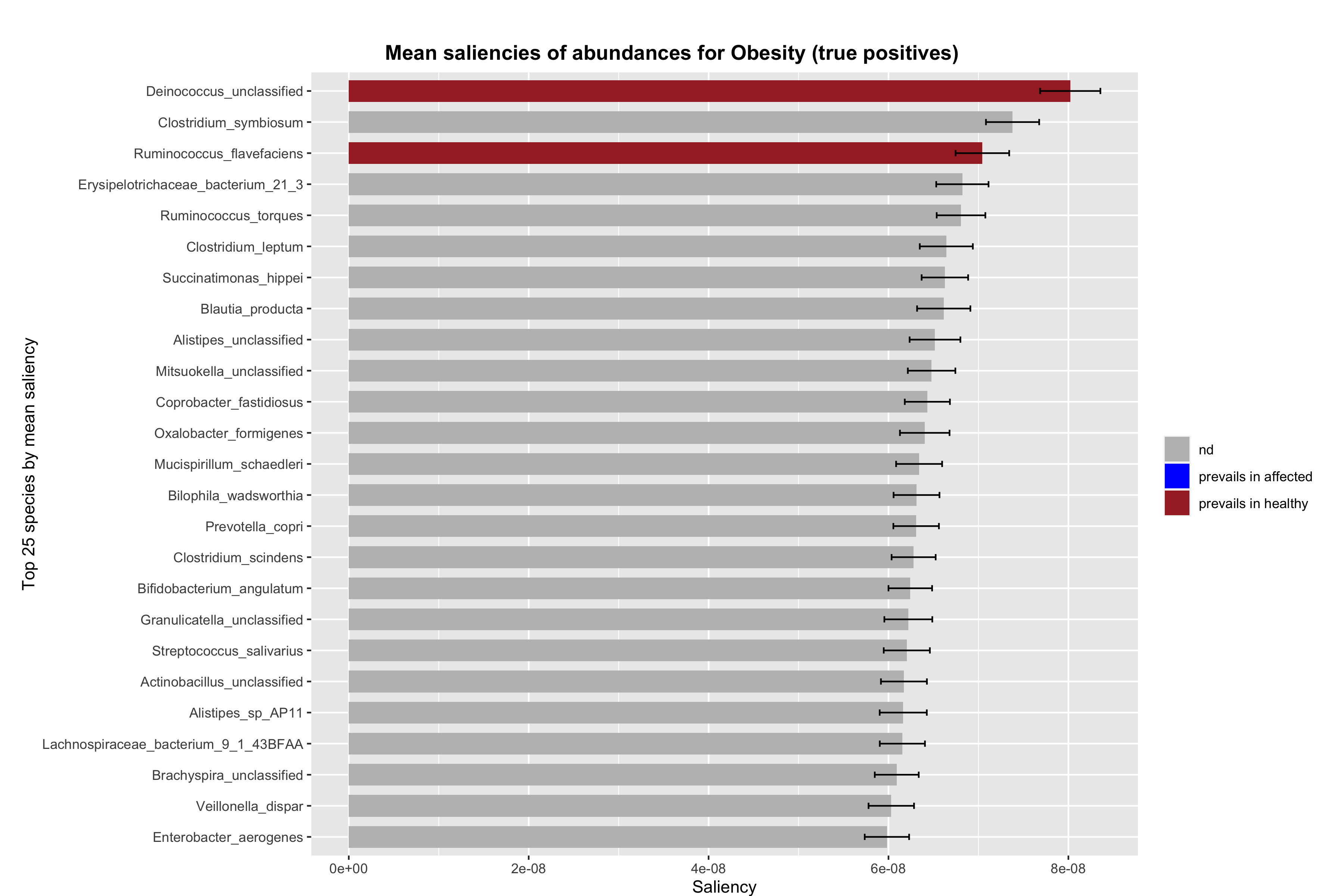

### abundance_errbarplot_TP_saliencies_no_rescale_pval-0.1.png

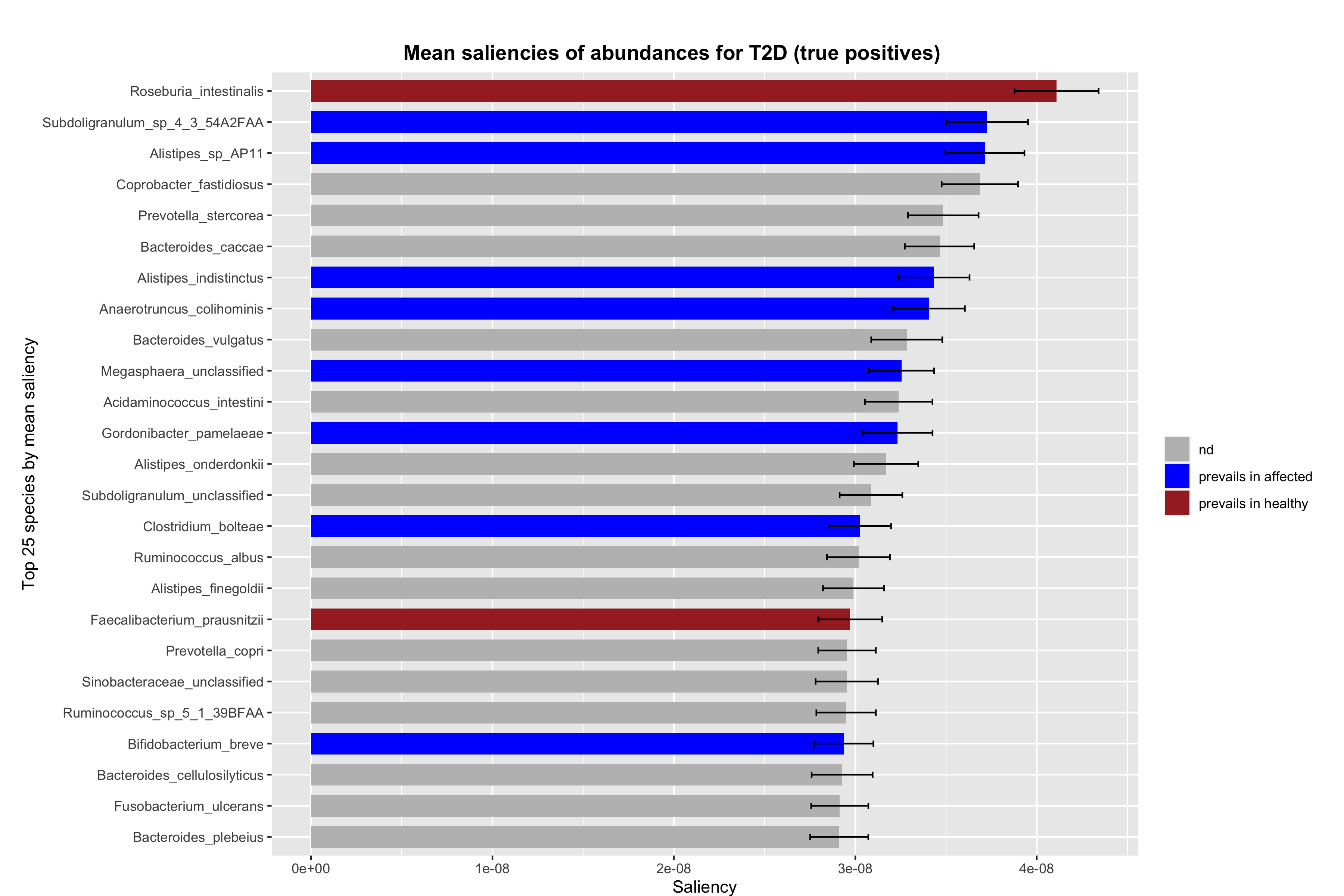

### abundance_errbarplot_TP_saliencies_no_rescale_pval-0.1.png

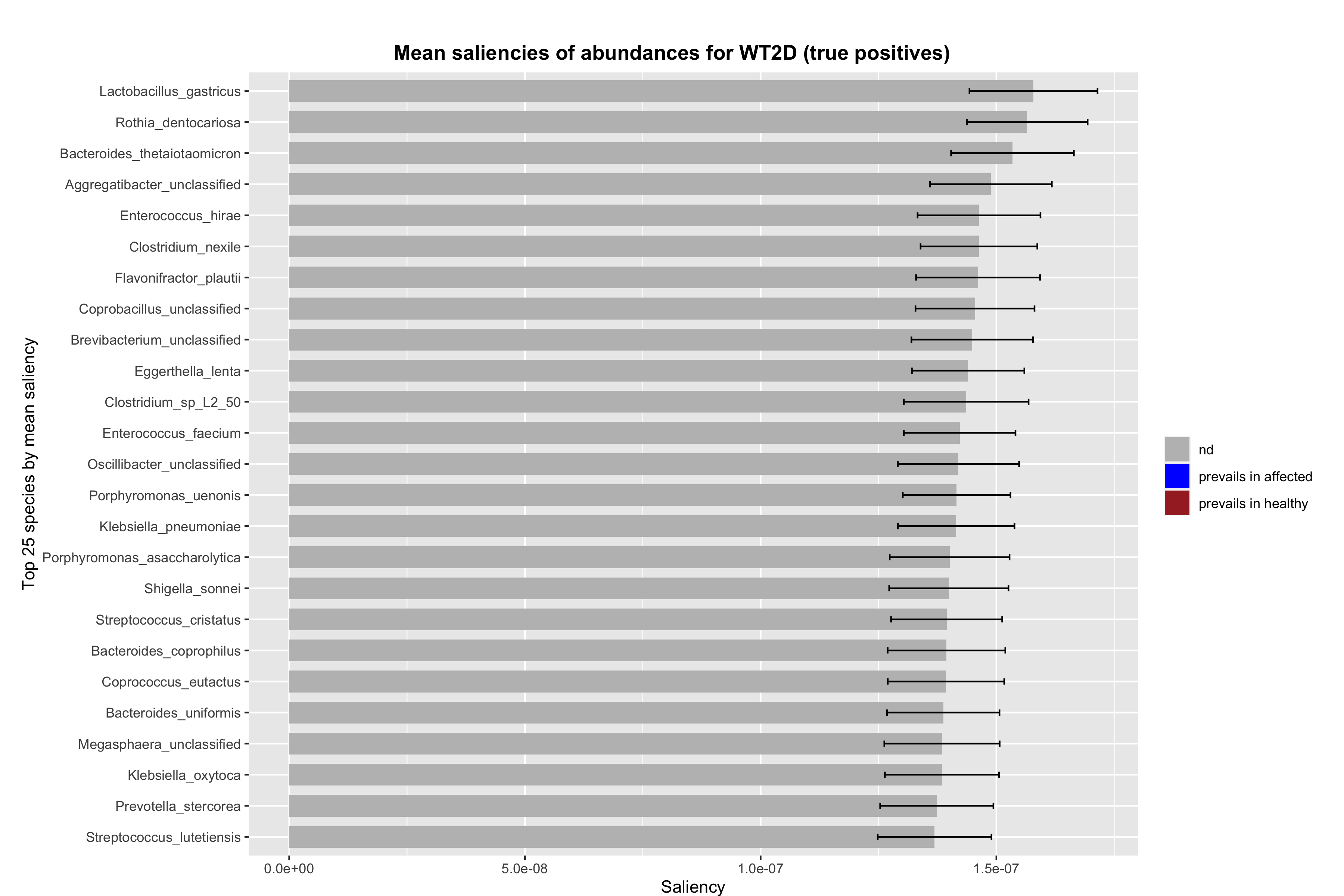

### abundance_violin_TP_saliencies_no_rescale_pval-0.1.png

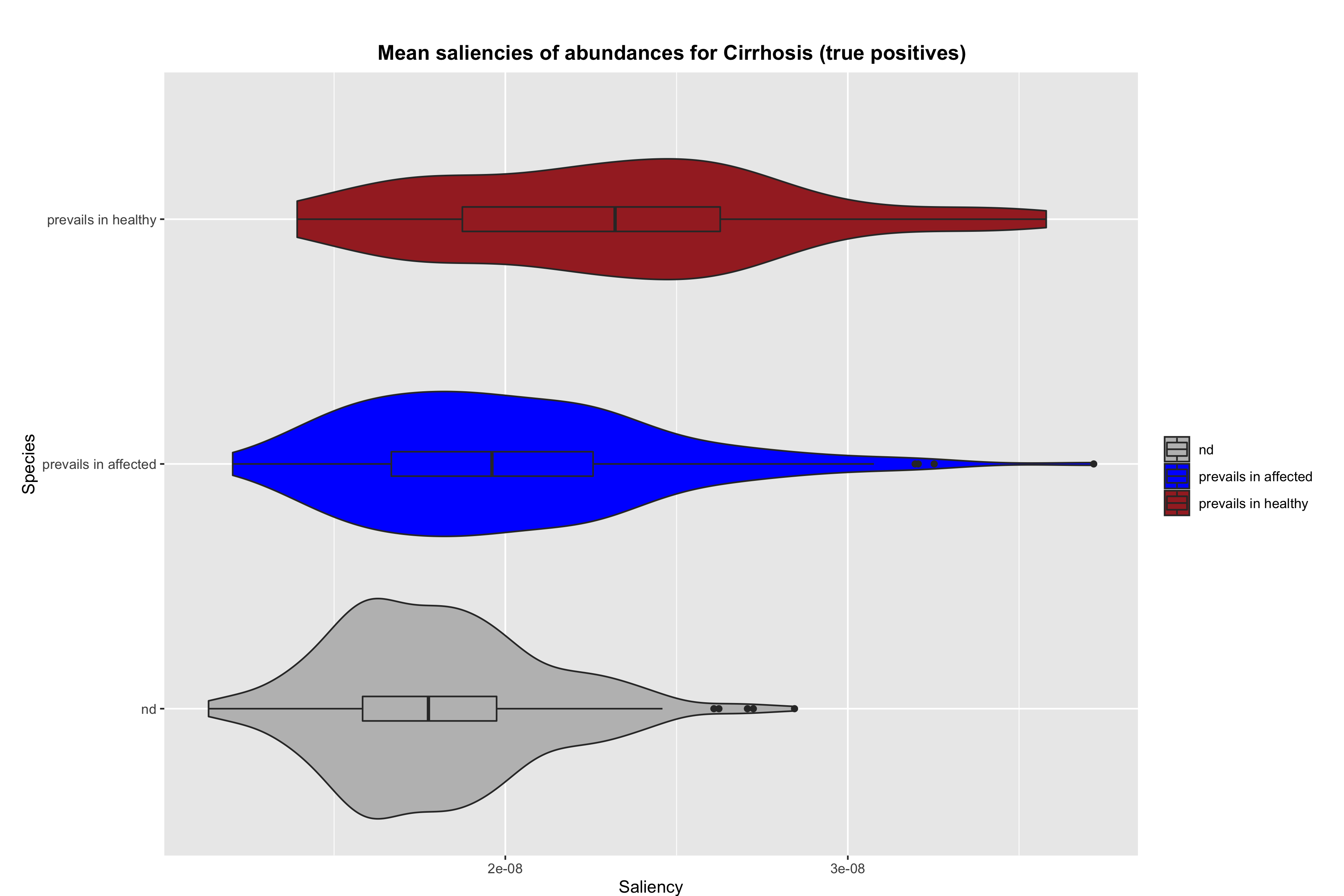

### abundance_violin_TP_saliencies_no_rescale_pval-0.1.png

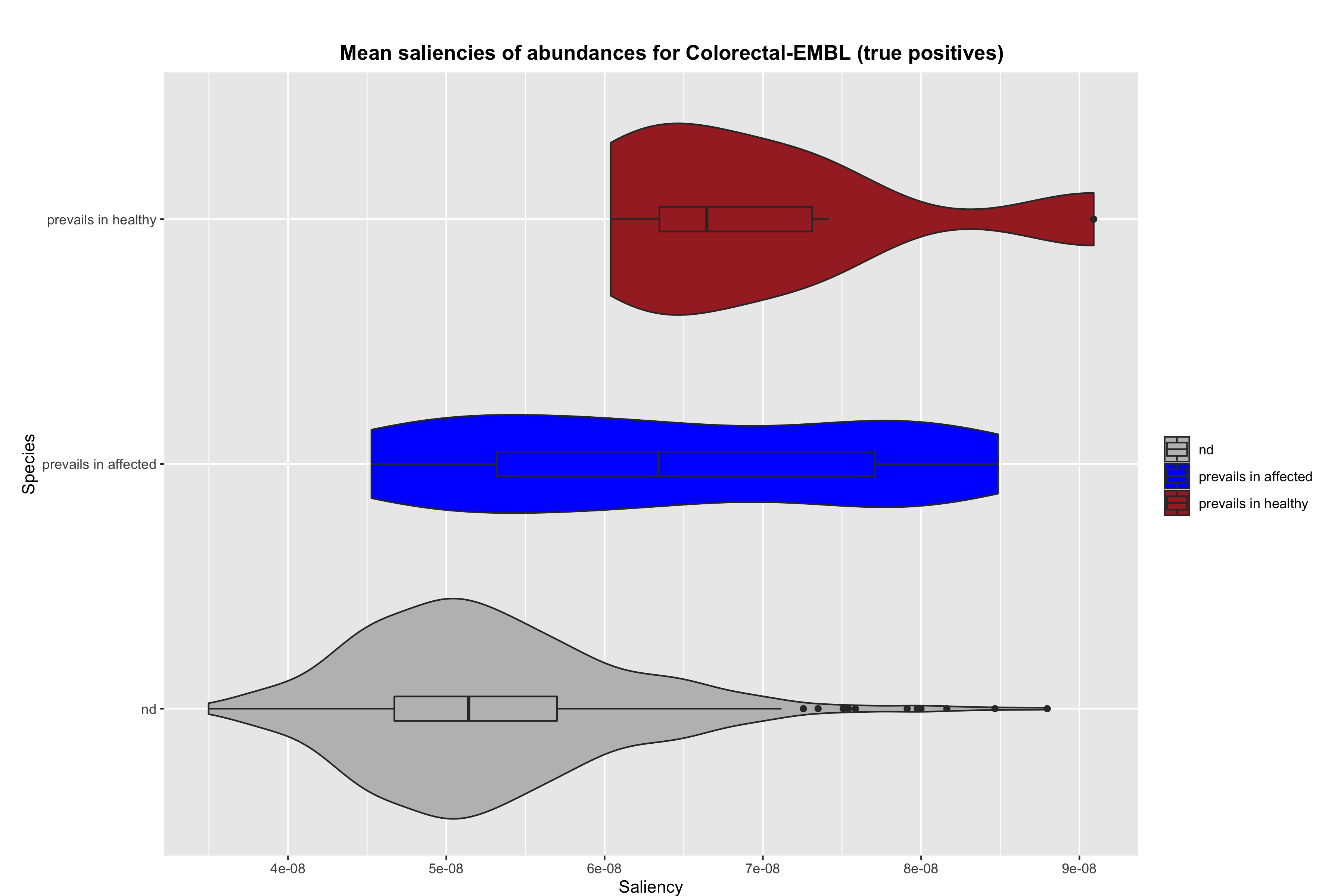

### abundance_violin_TP_saliencies_no_rescale_pval-0.1.png

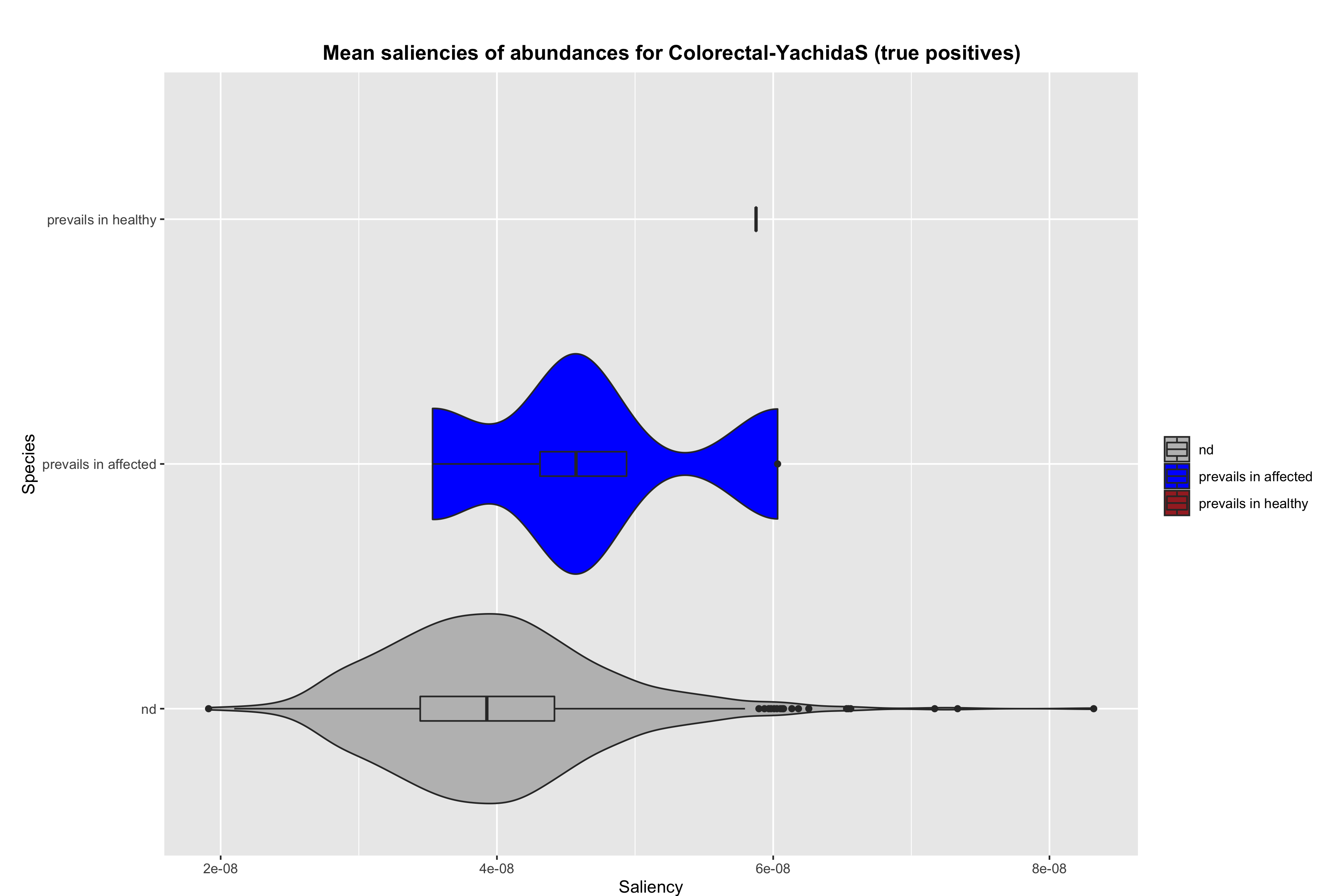

### abundance_violin_TP_saliencies_no_rescale_pval-0.1.png

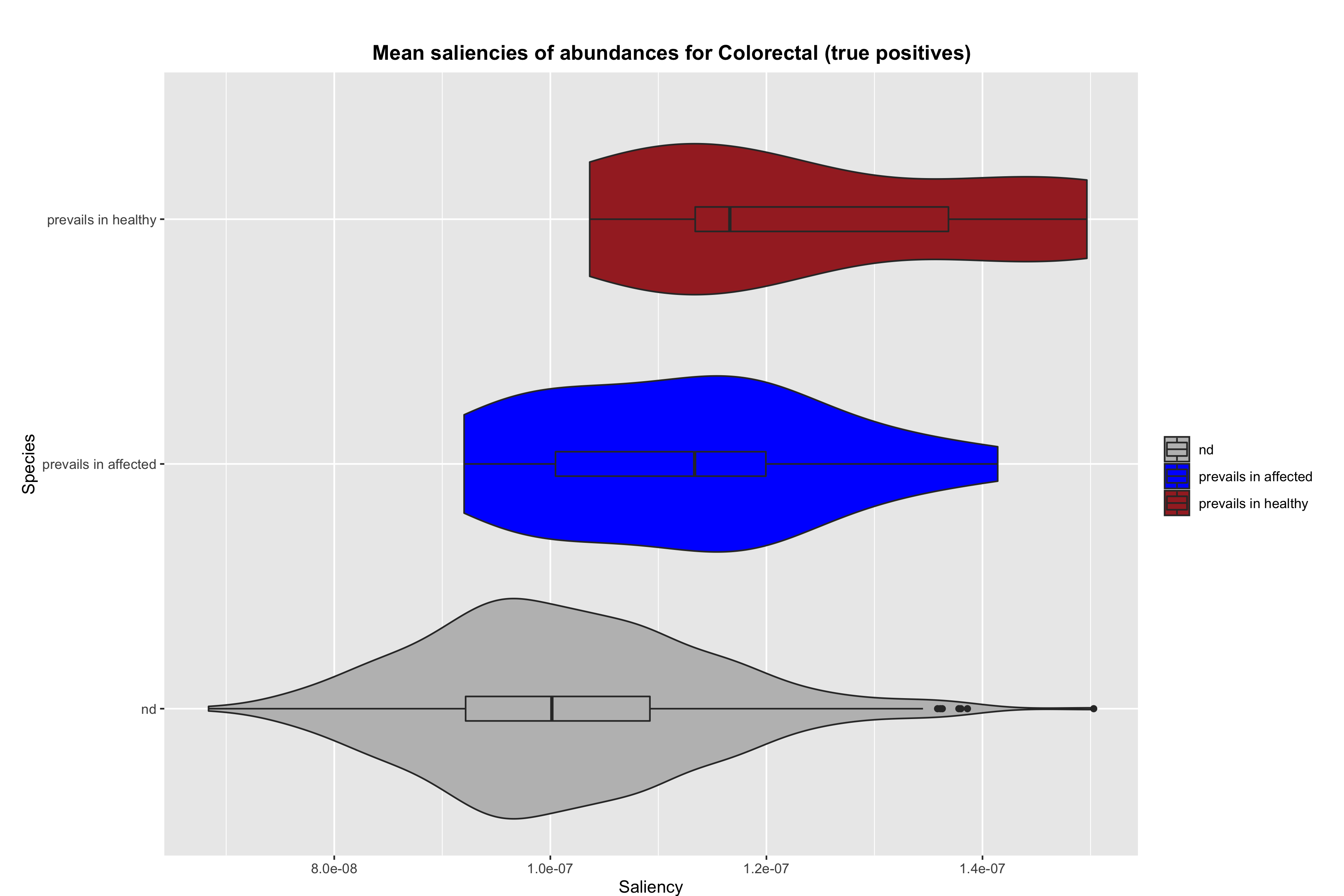

### abundance_violin_TP_saliencies_no_rescale_pval-0.1.png

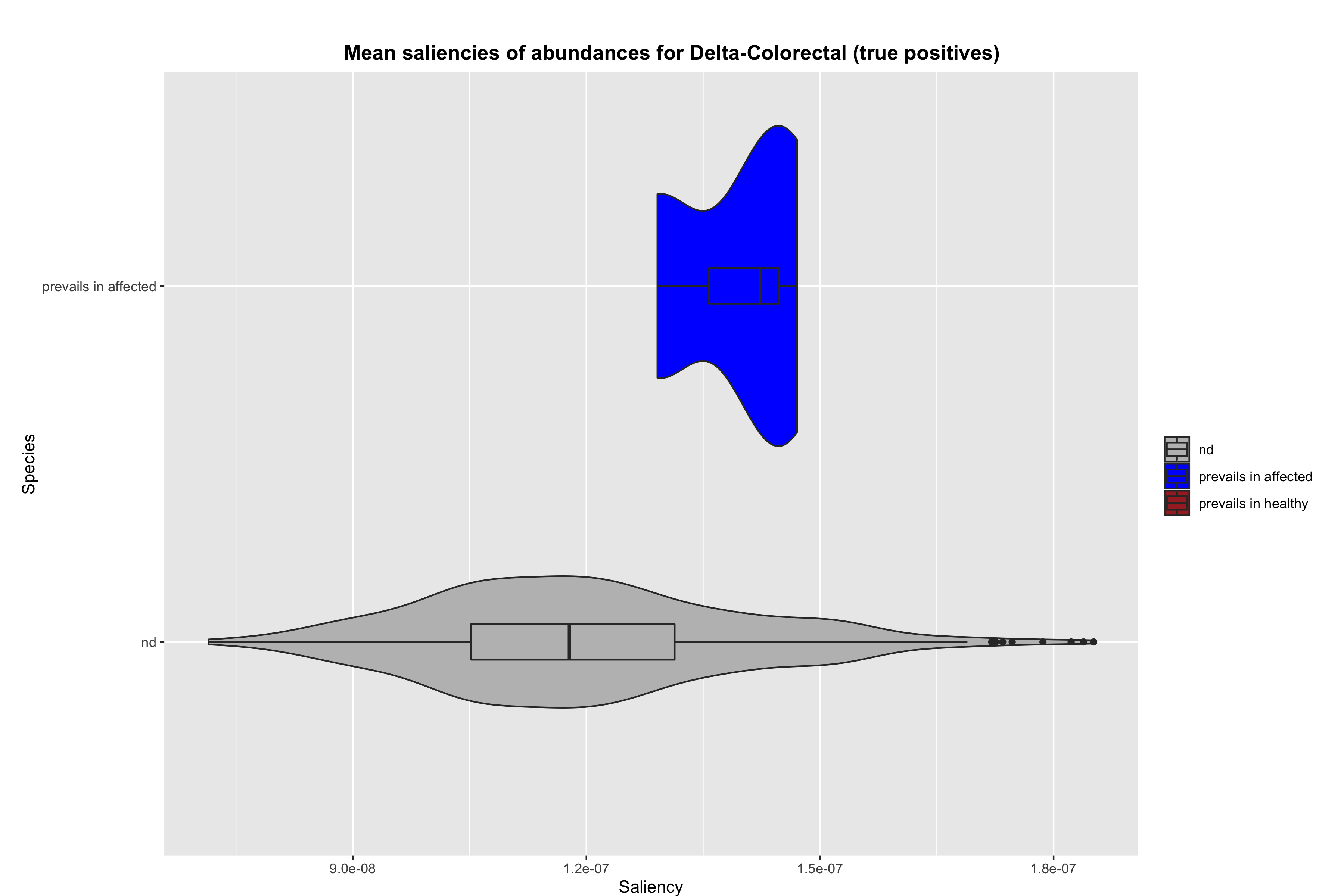

### abundance_violin_TP_saliencies_no_rescale_pval-0.1.png

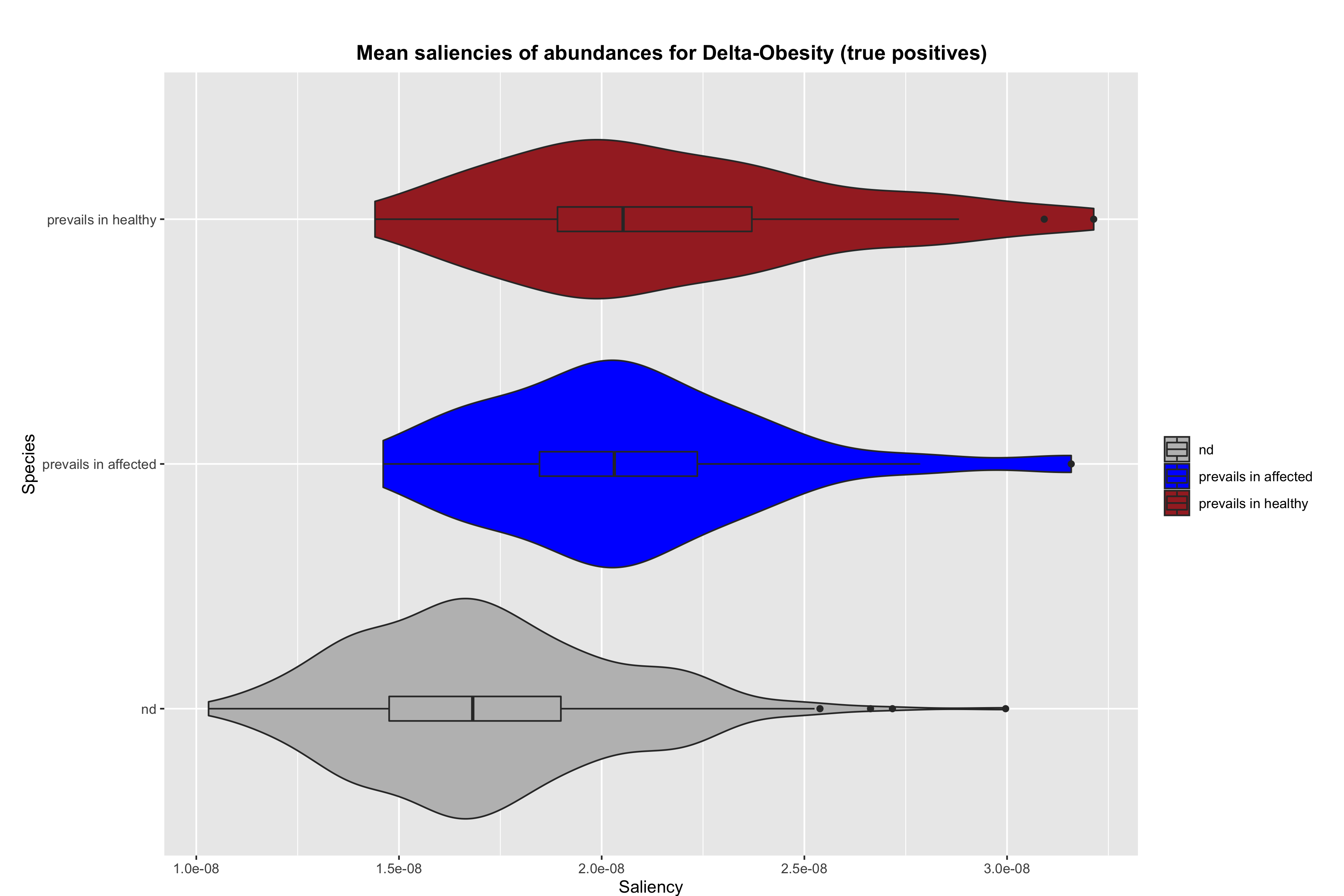

### abundance_violin_TP_saliencies_no_rescale_pval-0.1.png

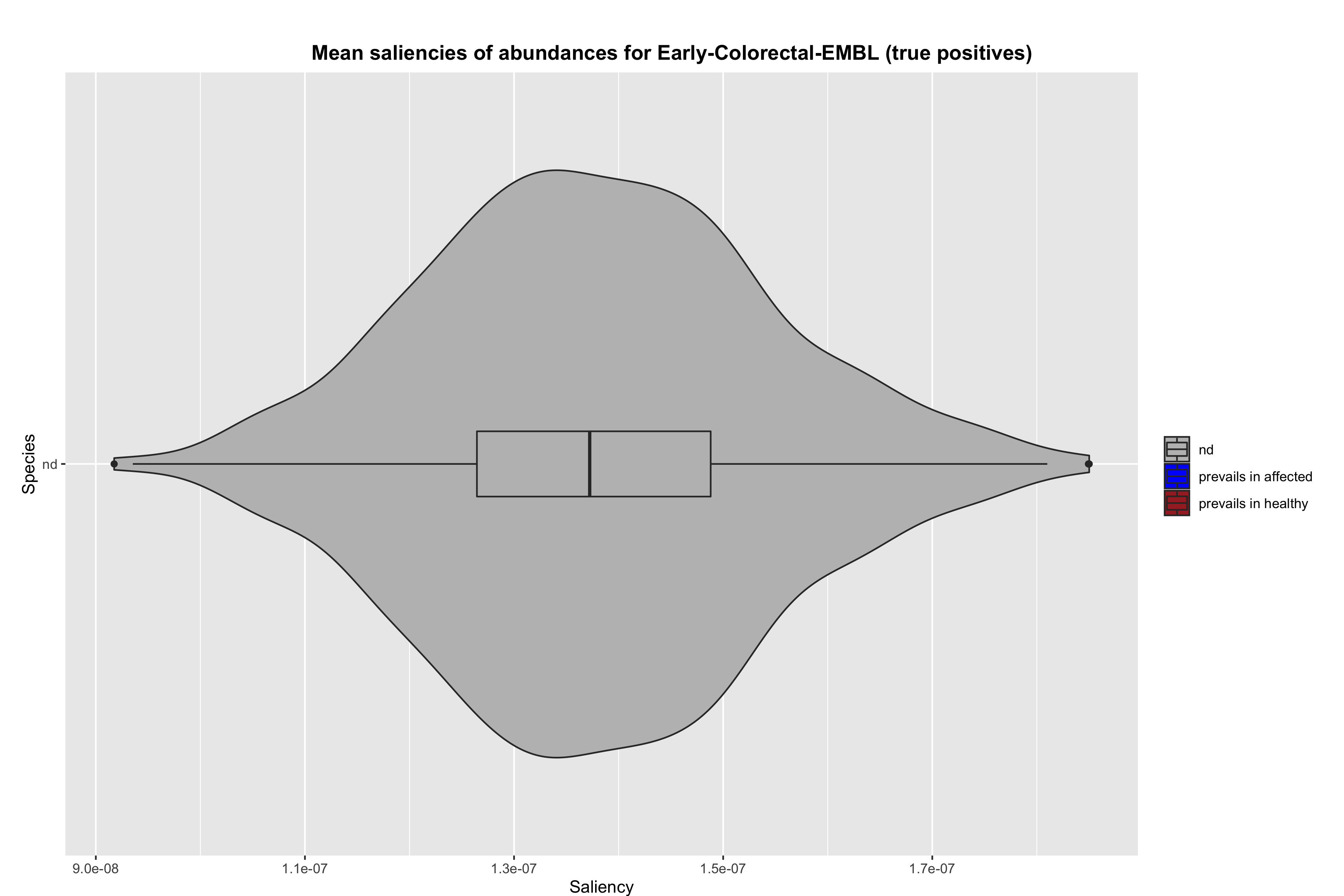

### abundance_violin_TP_saliencies_no_rescale_pval-0.1.png

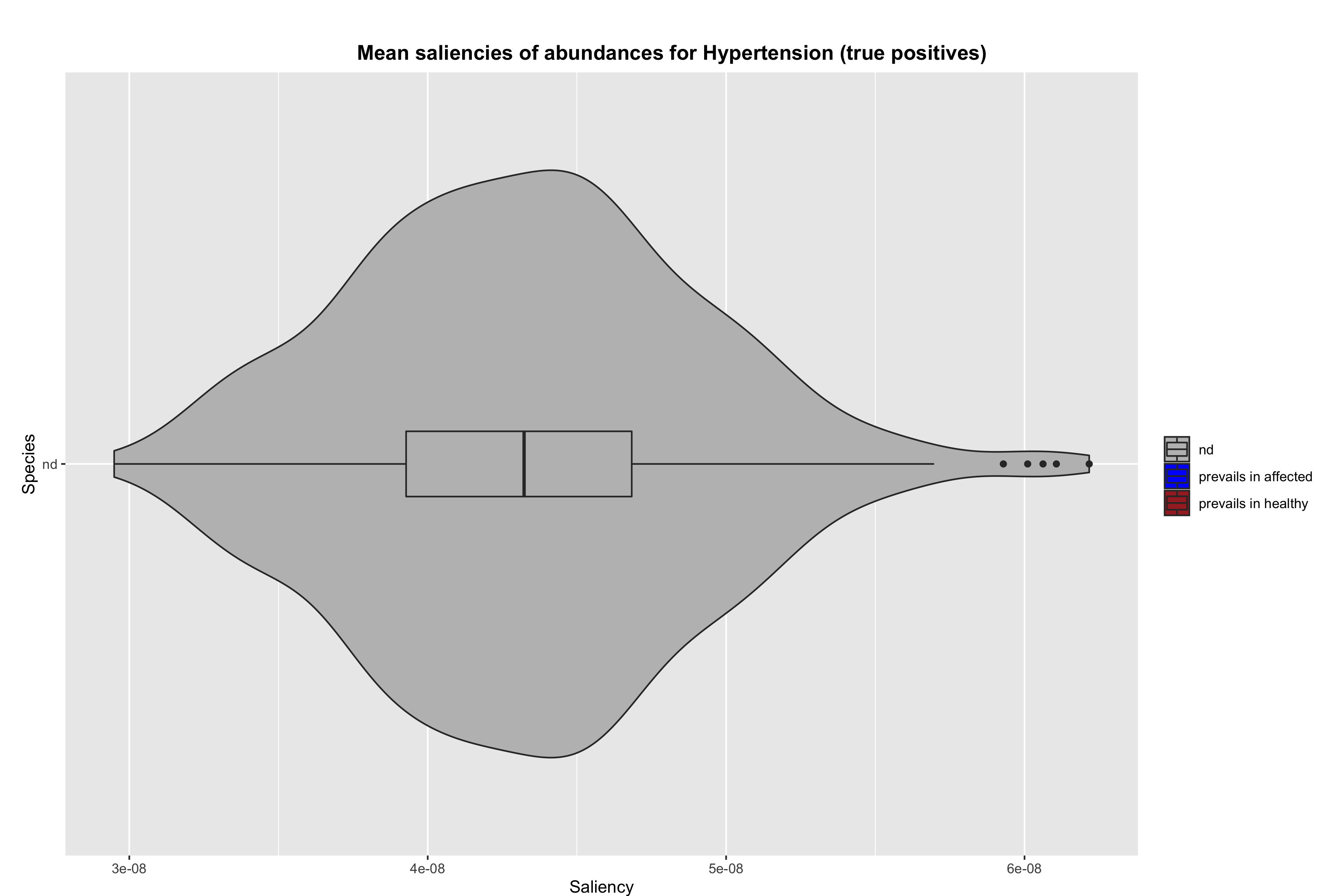

### abundance_violin_TP_saliencies_no_rescale_pval-0.1.png

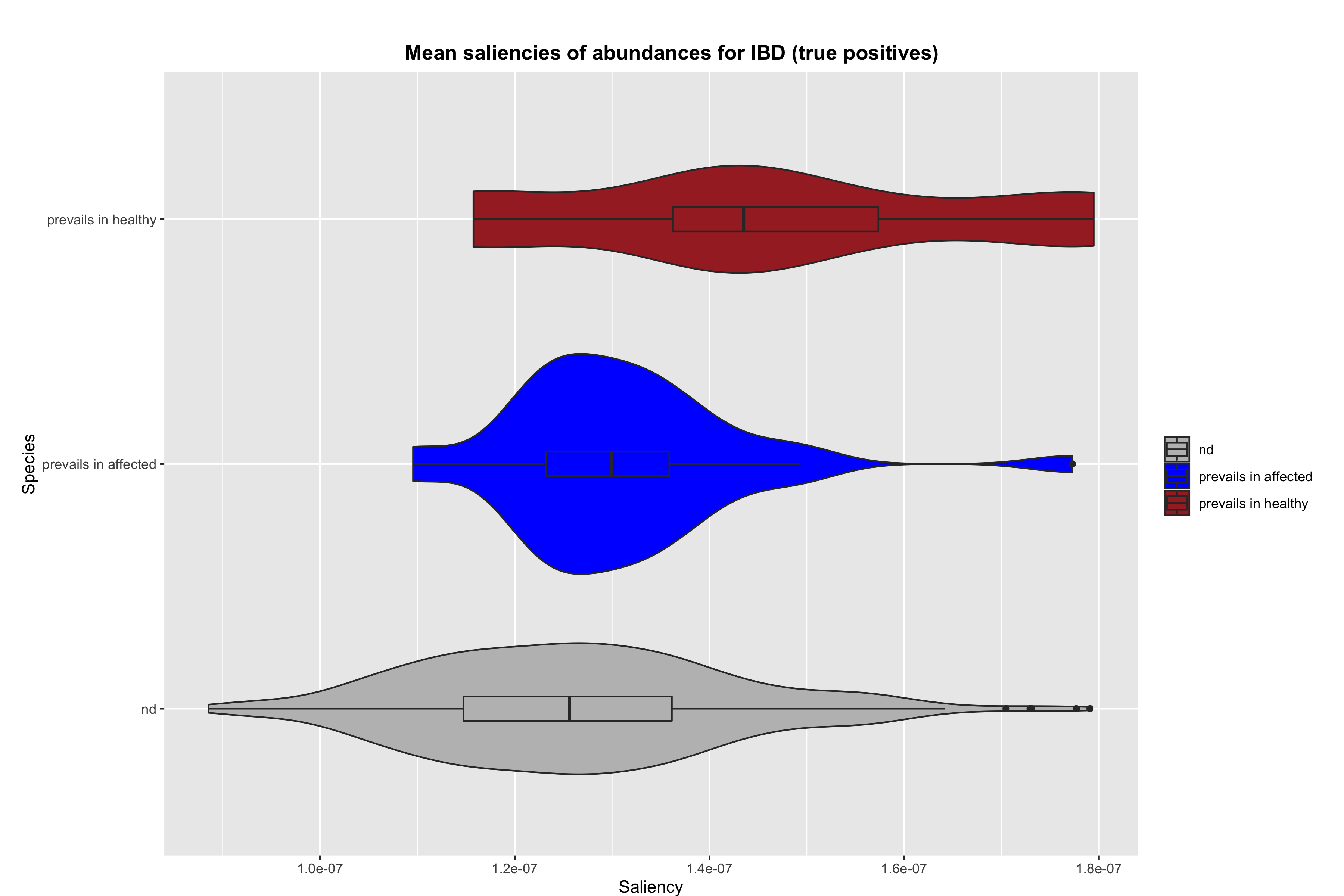

### abundance_violin_TP_saliencies_no_rescale_pval-0.1.png

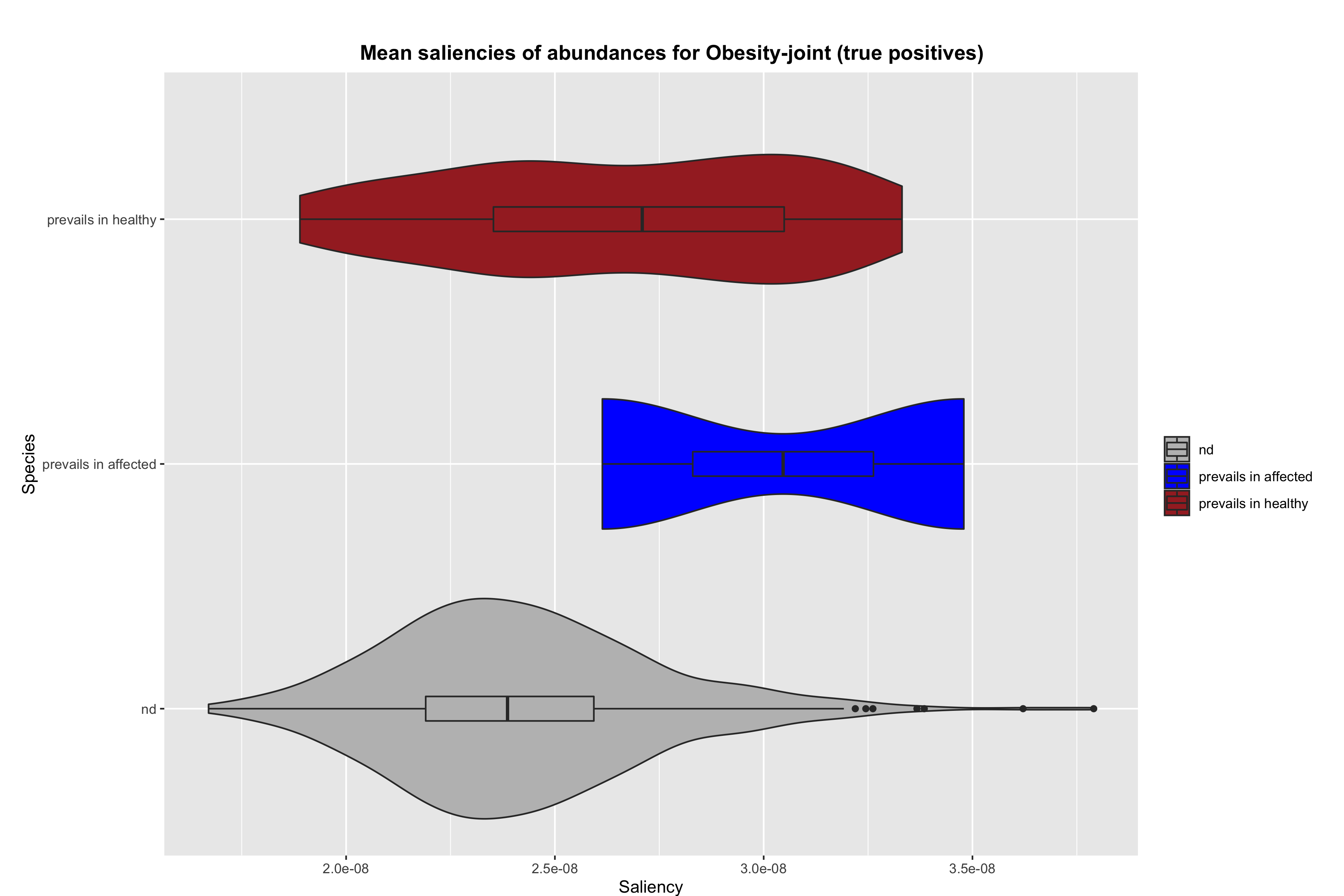

### abundance_violin_TP_saliencies_no_rescale_pval-0.1.png

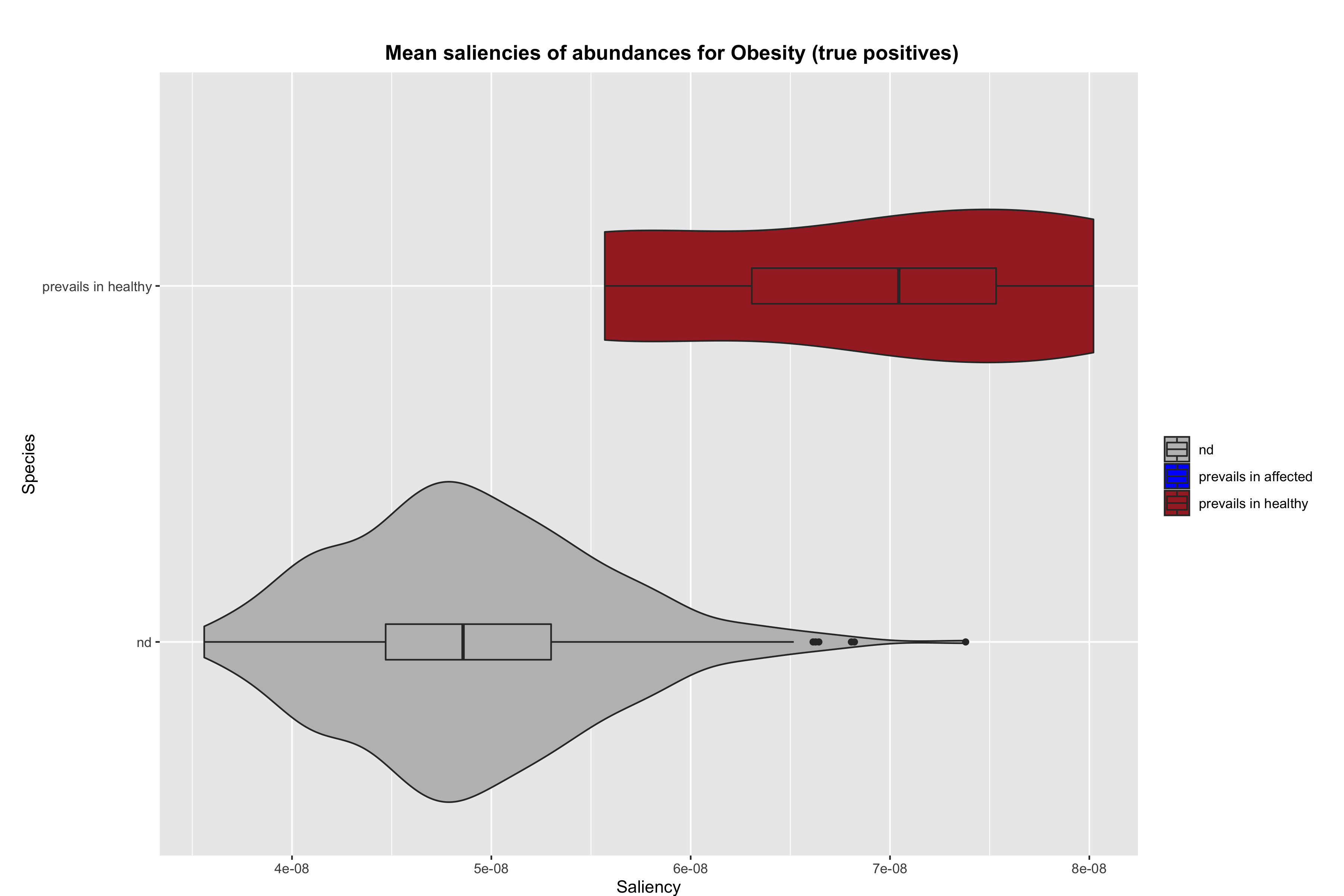

### abundance_violin_TP_saliencies_no_rescale_pval-0.1.png

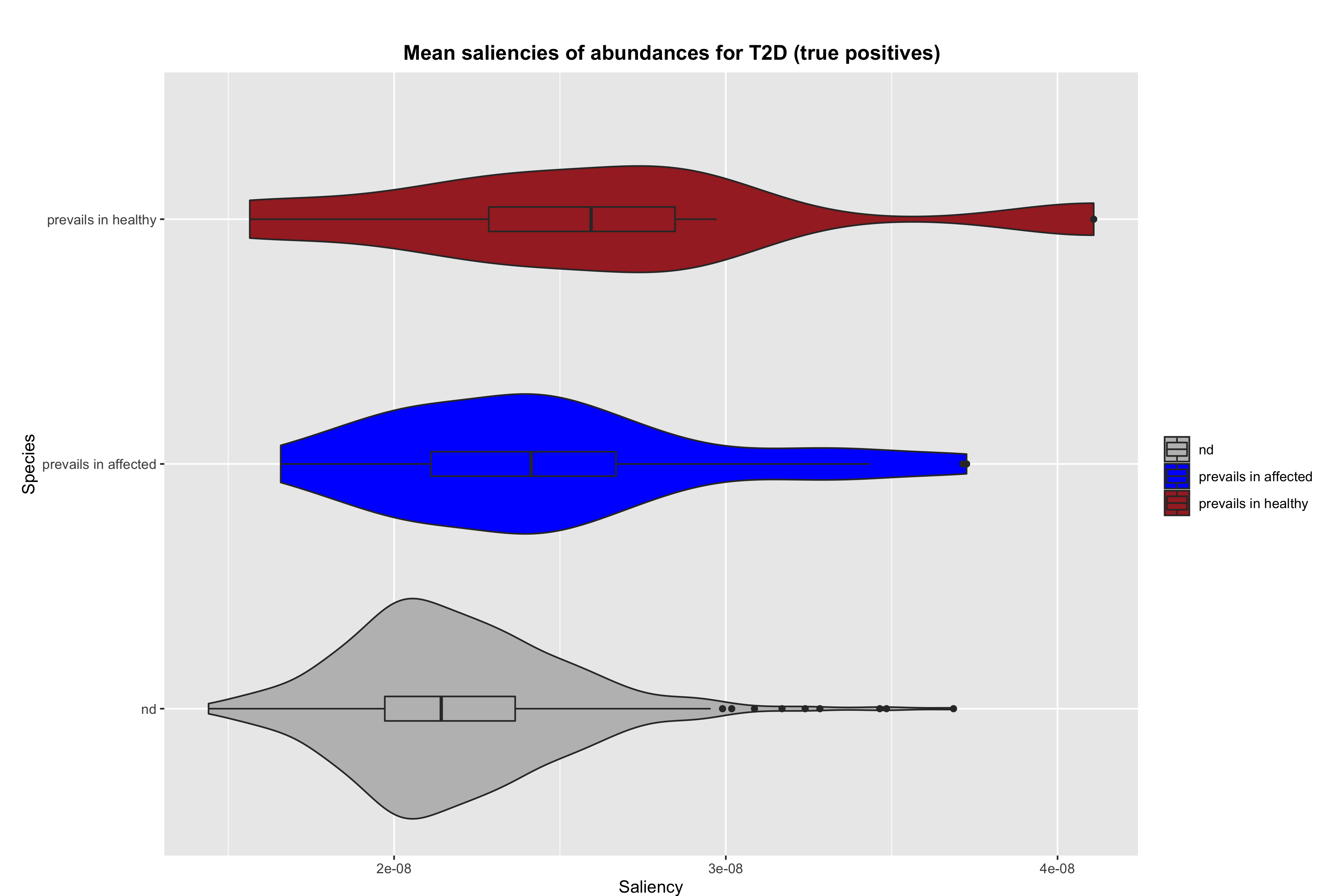

### abundance_violin_TP_saliencies_no_rescale_pval-0.1.png

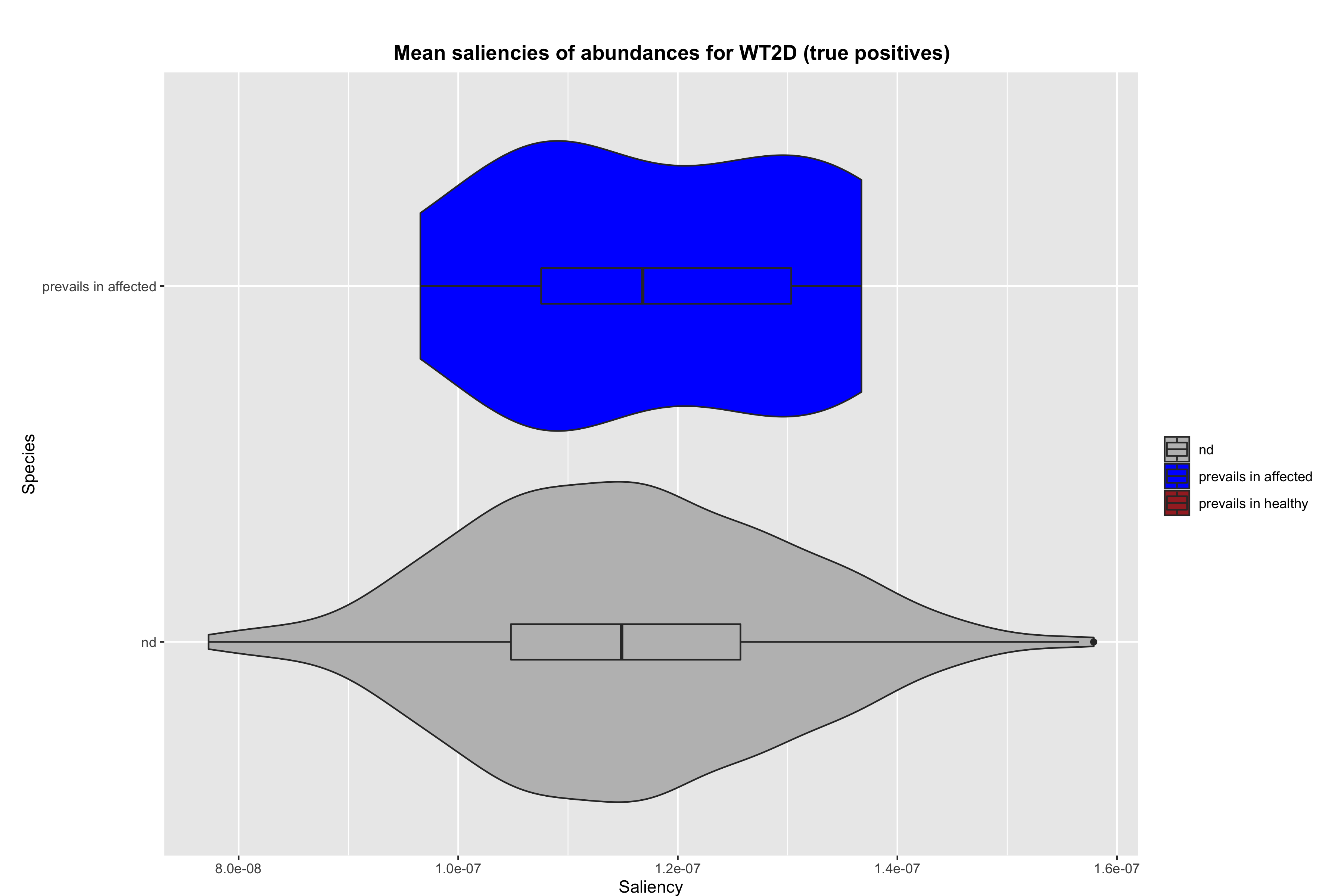

### markers_errbarplot_TP_saliencies_no_rescale.png

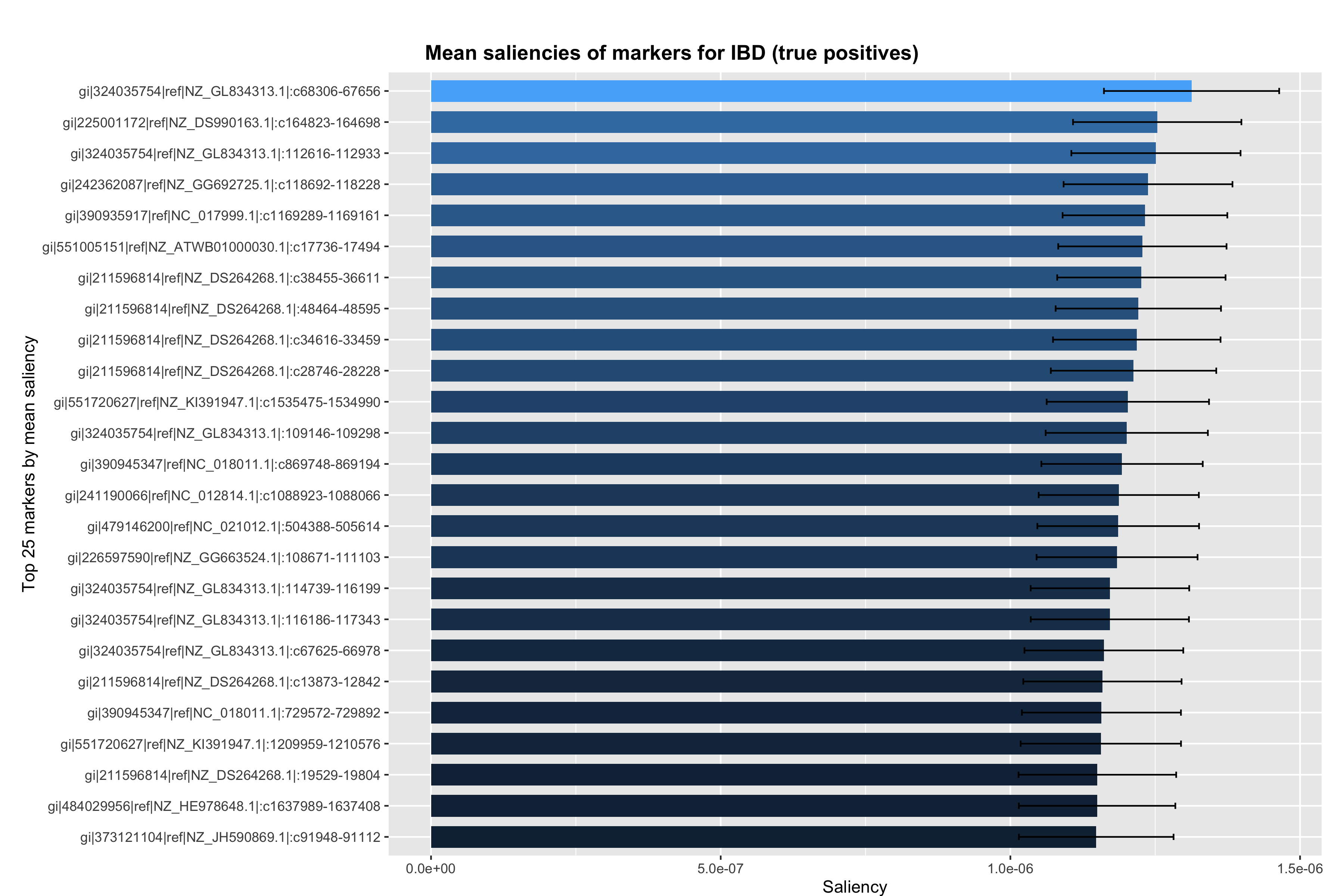

### markers_errbarplot_TP_saliencies_no_rescale.png

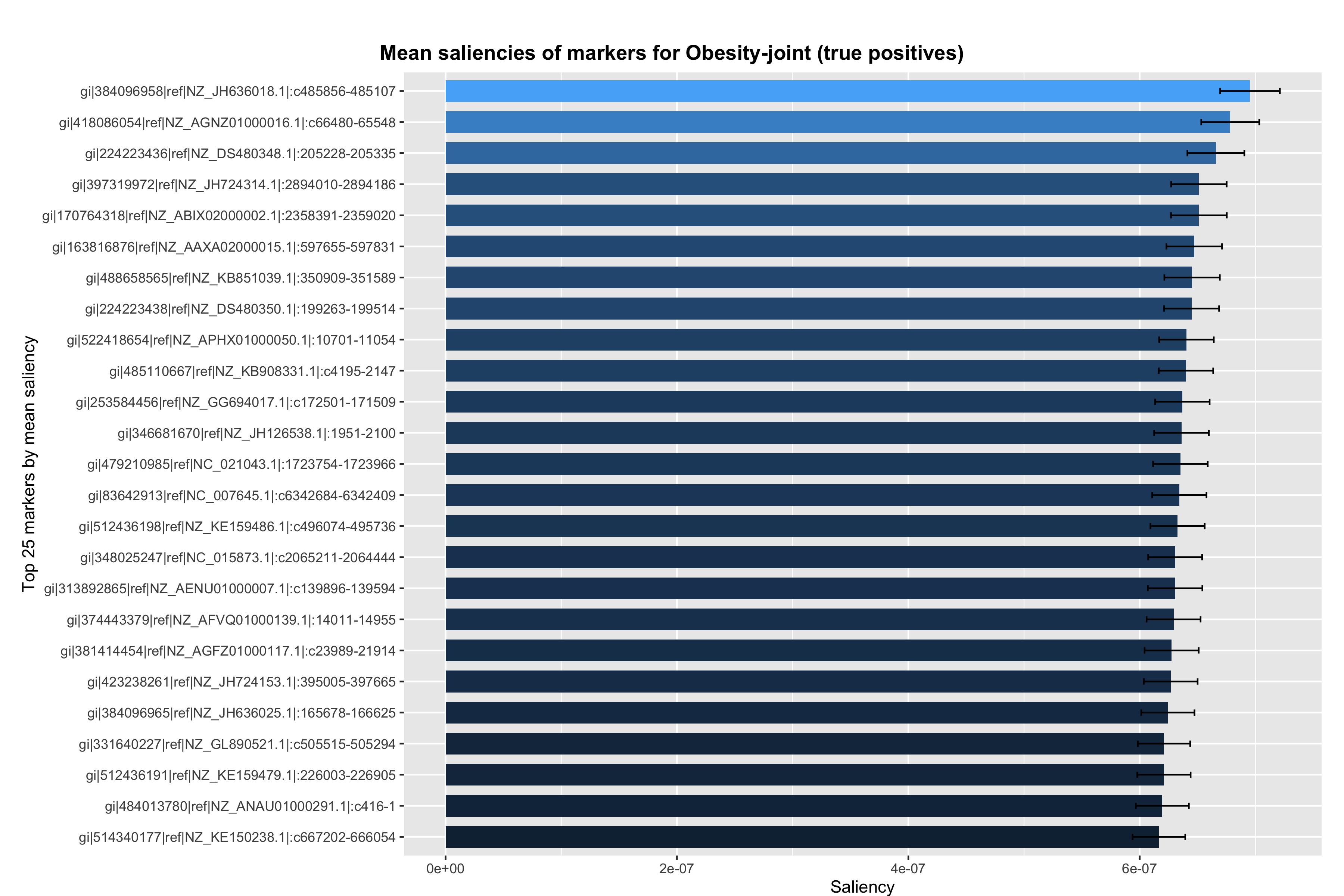

### markers_errbarplot_TP_saliencies_no_rescale.png

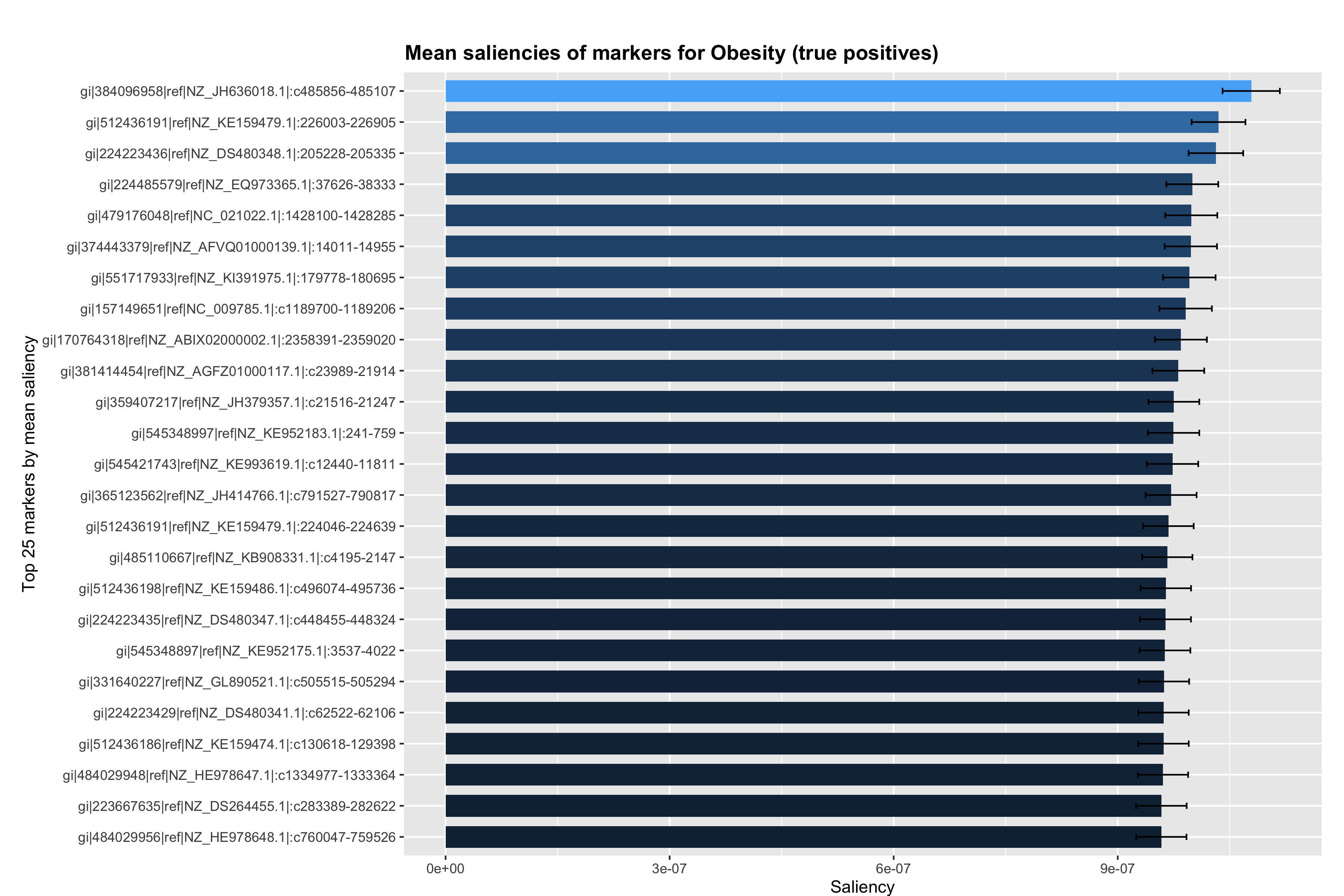

### markers_errbarplot_TP_saliencies_no_rescale.png

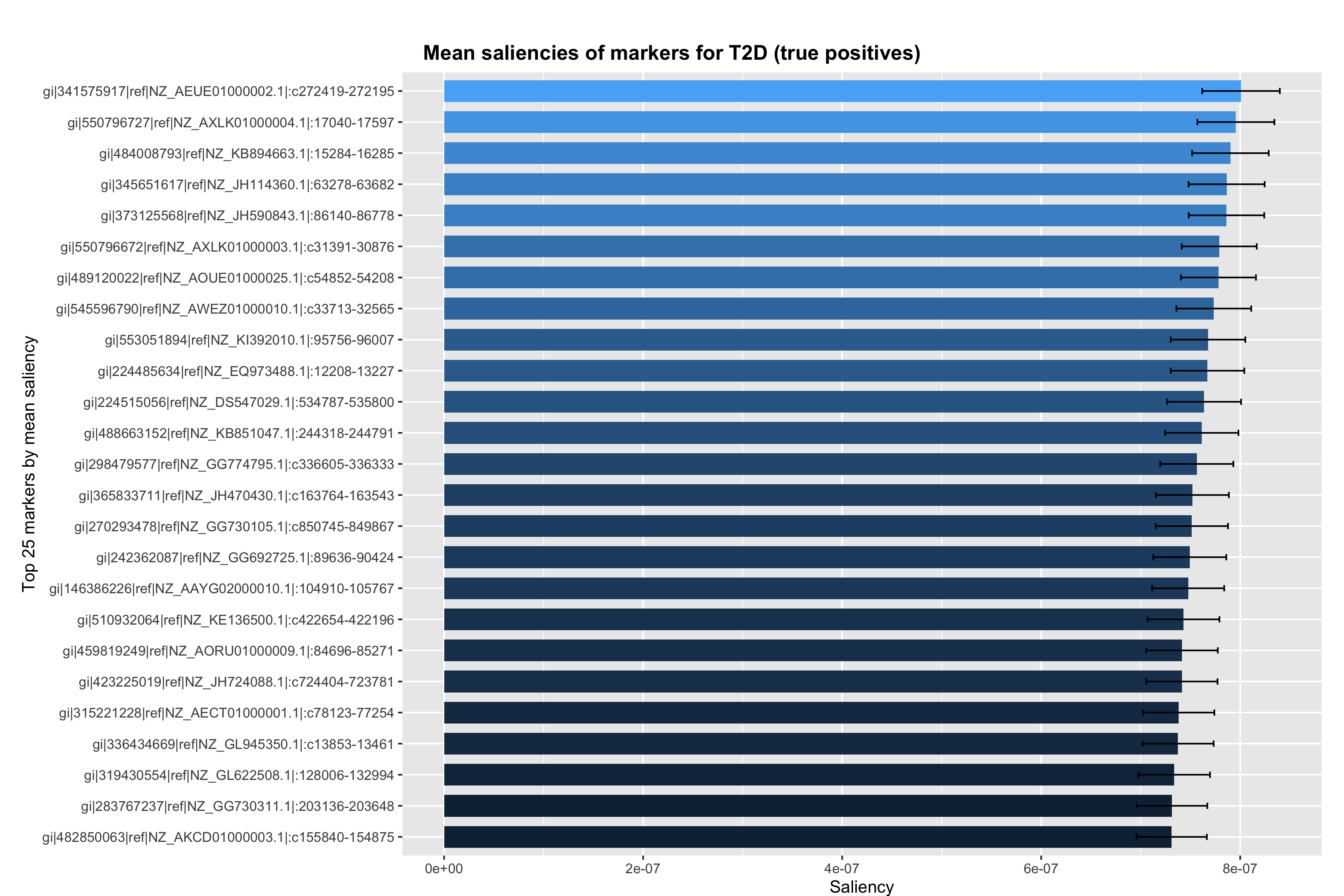

### markers_errbarplot_TP_saliencies_no_rescale.png

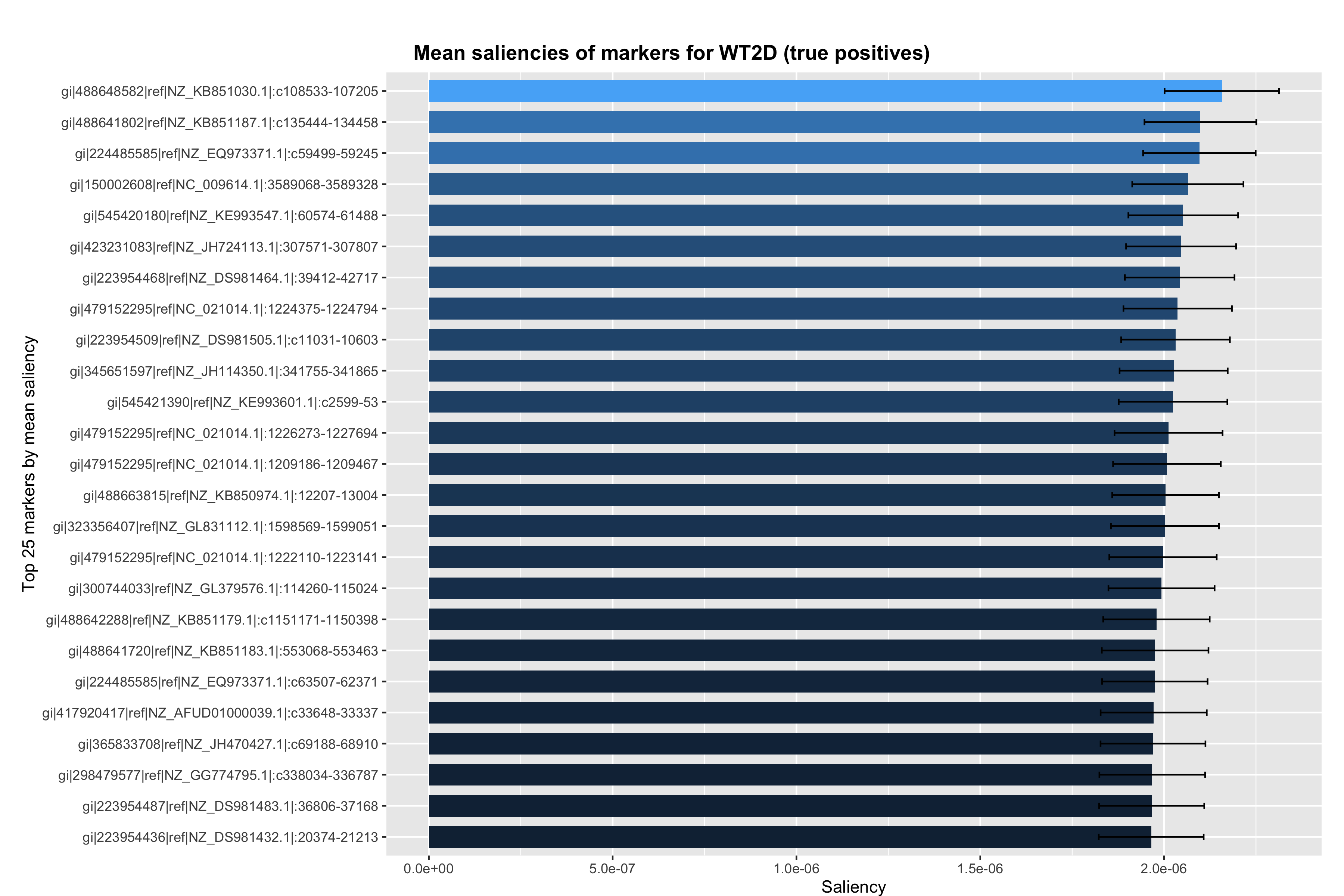
