## Supplementary material for "Microbiome-based disease prediction with multimodal variational information bottlenecks": S10 Table

**S7 Table. Experimental results for the Random Forest with default Scikit-learn implementation**

| | Random Forest - Default Scikit-learn implementation (RF-DEF) | | |
|---|---|---|---|
| **Dataset** | **A** | **M** | **A+M** |
| IBD | 0.878 (0.038) | 0.900 (0.018) | 0.878 (0.038) |
| EW-T2D | 0.796 (0.027) | 0.791 (0.041) | 0.796 (0.027) |
| C-T2D | 0.709 (0.022) | 0.743 (0.017) | 0.709 (0.022) |
| Obesity | 0.641 (0.026) | 0.567 (0.016) | 0.641 (0.026) |
| Cirrhosis | 0.890 (0.011) | 0.894 (0.012) | 0.890 (0.011) |
| Colorectal | 0.835 (0.038) | 0.785 (0.043) | 0.835 (0.038) |
| Obesity-joint | 0.798 (0.018) | 0.768 (0.030) | 0.798 (0.018) |
| Colorectal-EMBL | 0.866 (0.015) | 0.822 (0.024) | 0.866 (0.015) |
| Early-Colorectal-EMBL | 0.517 (0.048) | 0.521 (0.015) | 0.517 (0.048) |
| Hypertension | 0.652 (0.043) | 0.653 (0.025) | 0.652 (0.043) |

Experiments are executed five times with random independent training-test splits. Values in brackets refer to the standard error over the repeated experiments. Reported values are test ROC AUC.
